# AlphaFold predictions on whole genomes at a glance: a coherent view on packing properties, pLDDT values, and disordered regions

**DOI:** 10.1101/2024.11.16.623929

**Authors:** Frédéric Cazals, Edoardo Sarti

## Abstract

For model organisms, `AlphaFold` predictions show that 30% to 40% of amino acids have a (very) low pLDDT confidence score. This observation, combined with the method’s high complexity, commands a systematic analysis of `AlphaFold` predictions on whole genomes.

Consequently, using whole-genome predictions, we provide a coherent analysis on packing properties, pLDDT values, and their relationship with intrinsically disordered regions (IDRs). Our contributions are of two kinds. First, we introduce simple and explainable geometrical and topological statistics characterizing predictions. Second, we investigate four key biophysical and biological questions: (i) the clustering of `AlphaFold` predictions on whole genomes, (ii) the identification of high/low quality predicted domains, (iii) false positive/negative `AlphaFold` predictions with respect to IDRs, and (iv) the fragmentation of the polypeptide chain in terms of pLDDT values.

Altogether, our analysis provide novel insights into AlphaFold predictions across whole genomes, further enhancing the confidence assessment of the models.

## 1 Introduction

### 1.1 AlphaFold: complexity and predictions

`AlphaFold` ‘s ability to reliably predict protein structures from amino acid sequences has ushered in a new era in structural biology [1]. In recognition of this achievement, its leaders D. Hassabis and J. Jumper were jointly awarded the 2024 Nobel Prize in Chemistry. Each `AlphaFold` prediction is provided as a PDB file, with individual amino acids assigned a confidence score known as pLDDT (predicted Local Distance Difference Test), related to the Local Distance Difference Test [2]. The `AlphaFold` team defined four confidence categories based on pLDDT values: very high (0.9 ≤ pLDDT, dark blue), high (0.7 ≤ pLDDT *<* 0.9, cyan), low (0.5≤ pLDDT *<* 0.7, yellow), and very low (pLDDT *<* 0.5, orange).

On the experimental side, the Protein Data Bank currently holds approximately 230*K* experimentally determined structures, in contrast to the ∼ 200*M* protein sequences recorded in `UniProtKB`. This stark imbalance highlights the critical role of structure prediction in enabling genome-wide proteomic analysis (SWISS-MODEL Repository [3]). To address this, `AlphaFold` has been applied at scale, with predictions now available in the AlphaFold Protein Structure Database database–`AlphaFold-DB`, [4] and https://alphafold.ebi.ac.uk. A comprehensive analysis of this database reveals that for many model organisms, 30% to 40% of amino acids in predicted structures fall within the low or very low pLDDT confidence categories (Table 1, Section S6).

**Table 1:** Cumulated distribution function for pLDDT values of all a.a. of all proteins in a genome. See section S6 for statistics on median and mean values. Top: model genomes. Bottom: five *global health* proteomes. Min and max values per column stressed in bold.

| genome / pLDDT | 50 | 70 | 90 |
| --- | --- | --- | --- |
| AThaliana/all | 0.204 | 0.315 | 0.550 |
| CAlbicans/all | 0.213 | 0.312 | 0.570 |
| CElegans/all | 0.202 | 0.323 | 0.597 |
| DDiscoideum/all | **0.288** | 0.423 | **0.681** |
| DMelanogaster/all | 0.281 | 0.387 | 0.623 |
| DRerio/all | 0.246 | 0.343 | 0.593 |
| EColi/all | **0.029** | **0.078** | 0.275 |
| GMax/all | 0.217 | 0.338 | 0.577 |
| HSapiens/all | 0.284 | 0.382 | 0.666 |
| MJannaschii/all | 0.035 | 0.084 | **0.264** |
| MMusculus/all | 0.255 | 0.352 | 0.597 |
| OryzaSativa/all | 0.257 | **0.408** | 0.623 |
| RattusNorvegicus/all | 0.253 | 0.351 | 0.596 |
| SCerevisiae/all | 0.213 | 0.314 | 0.582 |
| SPombe/all | 0.187 | 0.289 | 0.558 |
| ZeaMays/all | 0.260 | 0.394 | 0.635 |
| SAureus/all | 0.045 | 0.096 | 0.295 |
| HPylori/all | 0.053 | 0.123 | 0.347 |
| MTuberculosis/all | 0.069 | 0.133 | 0.322 |
| Aeruginosa/all | **0.036** | **0.086** | **0.278** |
| PFalciparum/all | **0.460** | **0.584** | **0.804** |

From the methods’ perspective, `AlphaFold` uses two main modules, known as the `Evoformer` and the `structure module`. The overall algorithm is very complex, as evidenced by the ablation study which compares 10 variants to assess the merits of the main building blocks [1, SI]. The overall architecture heavily relies on transformers [5]. Beyond unsupervised approaches such as direct coupling analysis [6] or MSA transformers [7], the supervised inference of contacts by transformers within the `EvoFormer` is a key asset of `AlphaFold` [8]. In the context of the bias-variance trade-off in machine learning in general [9], and of the double descent phenomenon for large deep models [10], an important point is the balance between the number of parameters in `AlphaFold` and the pieces of information used to train it. The number of parameters has been estimated circa 100 millions. On the other hand, the number of non redundant sequences used in the self-distillation procedure is 350k [1, Section 1.3, SI], which, assuming a median size of 500 amino acids, results in about ∼ 1.75· 10^8^ amino acids. (Distillation is the process of transferring knowledge from a teacher to a student [11], a simplifying process bearing similarities to coreset calculations in a more algorithmic setting learning [12].)

### AlphaFold predictions: structural analysis

The structural features within `AlphaFold` predictions have been examined using the `Geometricus` algorithm[13], which represents protein structures through a series of steps: (i) a decomposition of the protein into structural fragments (based on *k*-mers or spherical neighborhoods), (ii) a discrete encoding of invariants (moments) of these fragments into shape-mers, and (iii) a counting step of shape-mers yielding an embedding of the protein. By clustering these embeddings – using non-negative matrix factorization – across 21 predicted proteomes from `AlphaFold-DB`, 250 structural clusters were identified, including 20 major clusters or superfamilies [14].

### AlphaFold predictions and intrinsic disorder

The pLDDT metric has been thoroughly analyzed, revealing its effectiveness as an indicator of result robustness. It has been observed that the low-pLDDT regions in `AlphaFold` predictions correlate with observed or predicted intrinsically disordered regions (IDRs) [15]. Indeed `AlphaFold` outperforms state-of-the-art methods such as IUPred2 on disordered region detection on a benchmark of `DisProt` targets and PDB controls [14]. Also, high mean pLDDT values are associated with minimal structural differences between `AlphaFold` and `trRosetta` models [14]. However, as indicated by a lack of association with B-factors, pLDDT does not appear to correlate with flexibility [16]. Although pLDDT can predict structural disorder, alternative estimators using `AlphaFold`-predicted structures, such as the relative solvent accessibility (RSA), provide an even more accurate measure of disorder [17].

### 1.2 Contributions

The significant percentage of amino acids with a (very) low pLDDT and the `AlphaFold` complexity call for further investigations to better understand the quality of predictions (Table 1). More specifically, using whole-genome predictions, our goal is to provide a coherent perspective on packing properties, pLDDT values, and their relationship with intrinsically disordered regions or proteins (IDPs/IDRs). Our contributions are of two kinds: first, we introduce two geometric/topological analysis methods; second, using these methods, we investigate four key biological questions.

## Methods

Topological data analysis provides a set of tools and concepts to extract stable features from high-dimensional data [18, 19]. To understand properties of `AlphaFold` predictions, we use the concept of *filtration*, namely a progressive encoding of a structure as a sequence of nested shapes starting with the empty set and ending with the entire structure, monitored by a real-valued parameter. In practice, we use two filtrations: the first one is associated with the number of neighbors of a *C*_*α*_ carbon–a local packing measure; the second one is based on the pLDDT value provided by `AlphaFold`. Distributions associated to these filtrations are used to compare structures using optimal transport theory [20, 21], and for pLDDT based analysis, to assess structures with respect to a null (random) model.

These tools are used to study four biological questions.

### Q1. Contacts, packing, and whole genome predictions

Given the prominent role played by contacts in the `EvoFormer`, our first analysis focuses on packing properties of predictions. An embedding of `AlphaFold` predictions based on shape-mers has been obtained using dimensionality reduction (t-SNE, [14, Fig. 2]), but the corresponding non linear coordinates are not easily exploited to study correlations with other properties.

In search of an embedding performing at once dimensionality reduction and clustering in the context of packing properties, we introduce the *arity*, an encoding based on percentiles of the distribution representing the number of neighbors of each amino acid. Such a distribution can be summarized by a small number of integers. Using two numbers yields the *arity map*, a 2D natural low dimensional embedding of all structures, which we use to investigate the quality of domains predicted by `AlphaFold` and the relationship with IDPs/IDRs.

### Q2. Predicted domains and their quality

The aforementioned complexity of `AlphaFold` and the commensurate information size raises the question of potential biases in the model. To address this, we examine the locations of predicted low-quality domains. To understand our approach, it is important to recall that protein domain classification is a cornerstone of protein science [22, 23, 24]. A particularly noteworthy effort is the recently published ECOD structural domain classification of the entire human proteome [25], which leveraged `AlphaFold` predictions to enrich its classification with several new structural domains categorized by confidence scores as high or low. A domain is classified as low-confidence if it appears globular by its predicted aligned error (PAE) description but lacks a strong homologous hit (via a HMM calculation) within the `ECOD` reference–[25] and prodata.swmed.edu/ecod/.

We investigate whether the training of `AlphaFold` introduces biases in the prediction of low-quality domains by analyzing their distribution across the arity map. Notably, we identify a specific region of the arity map that is disproportionately populated by such domains.

### Q3. Predictions and intrinsically disordered proteins/regions

The per-residue pLDDT scores on the `AlphaFold-DB` human proteome prediction yields a bimodal distribution where the two peaks identify ordered and disordered regions [26]. Nonetheless, there are exceptions: conditionally folded IDRs are usually predicted as stably folded regions, and studies seems to imply that 10% of the low-pLDDT regions are not related to disorder [26].

To complement these findings, we show that although most of the `AlphaFold` disorder predictions are supported by experimental evidence, predictions with certain structural characteristics (a specific region in the arity map) are much richer in false positive IDR detection than others.

### Q4. pLDDT values and fragmentation of AlphaFold reconstructions

The pLDDT value is a key statistics to assess `AlphaFold` predictions, but its coherence along the primary structure has not been studied. We provide this analysis via a multiscale characterization of stretches along the sequence with coherent pLDDT values. This analysis relies on the stability of connected components of amino acids (a.a.) along the protein sequence, upon building a suitable filtration which consists of inserting the a.a. by decreasing pLDDT values.

## 2 Methods

### 2.1 Packing analysis

**Arity and arity signature**. A natural parameter to identify compact regions in proteins in the number of neighbors of a *C*_*α*_ carbon. Let the *arity* of a *C*_*α*_ carbon be the number of neighboring *C*_*α*_s within a distance range *r*. As classically done, we take *r* = 10Å to account for non-covalent contacts. We note in passing that such neighbors are sufficient to robustly infer protein domains using a direct application of spectral clustering[27]. Assuming the polypeptide chain has *n* amino acids, define:

- *L*_*a*_ = {*a*_*n*_, …, *a*_*n*_}: arities of the *n C*_*α*_ carbons;
- *L* = [*A*_1_, …, *A*_*m*_]: unique arities sorted by increasing value;

The number of *C*_*α*_ with a given arity is used to define the histogram 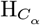. We compare two normalized such histograms (unit cumulated sum) using the mass transport solver from [28] implemented in the *Python Optimal Transport* package [29].

To simplify the presentation provided the 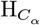 and the associated 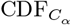, we define the following signature:

#### Definition. 1

*(Arity signature* 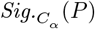 *of a polypeptide chain P*.) *Consider the cumulative distribution functions associated with the number of C*_*α*_ *carbons, that is* 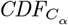.*Consider a list P*_*K*_ = {*p*_1_, …, *p*_*K*_} *of increasing percentiles. For p*_*k*_ ∈ *P*_*K*_, *let* `arity`*_C*_*α*_(*p*_*k*_) *be the smallest arity such that the CDF is larger than p*_*k*_, *that is*

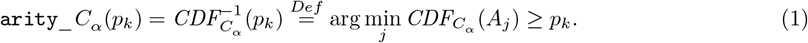

*The C*_*α*_ signature *is the increasing sequence*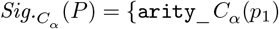,…, `arity`*_C*_*α*_(*p*_*K*_)}.

Using two quantiles {*q*_1_ = 0.25, *q*_2_ = 0.75}, the arity signature refers to the two arity values required to gather 25% and 75% percent of the number of amino acids in the chain (Fig. 1 for prototypical folds).

**Figure 1:**
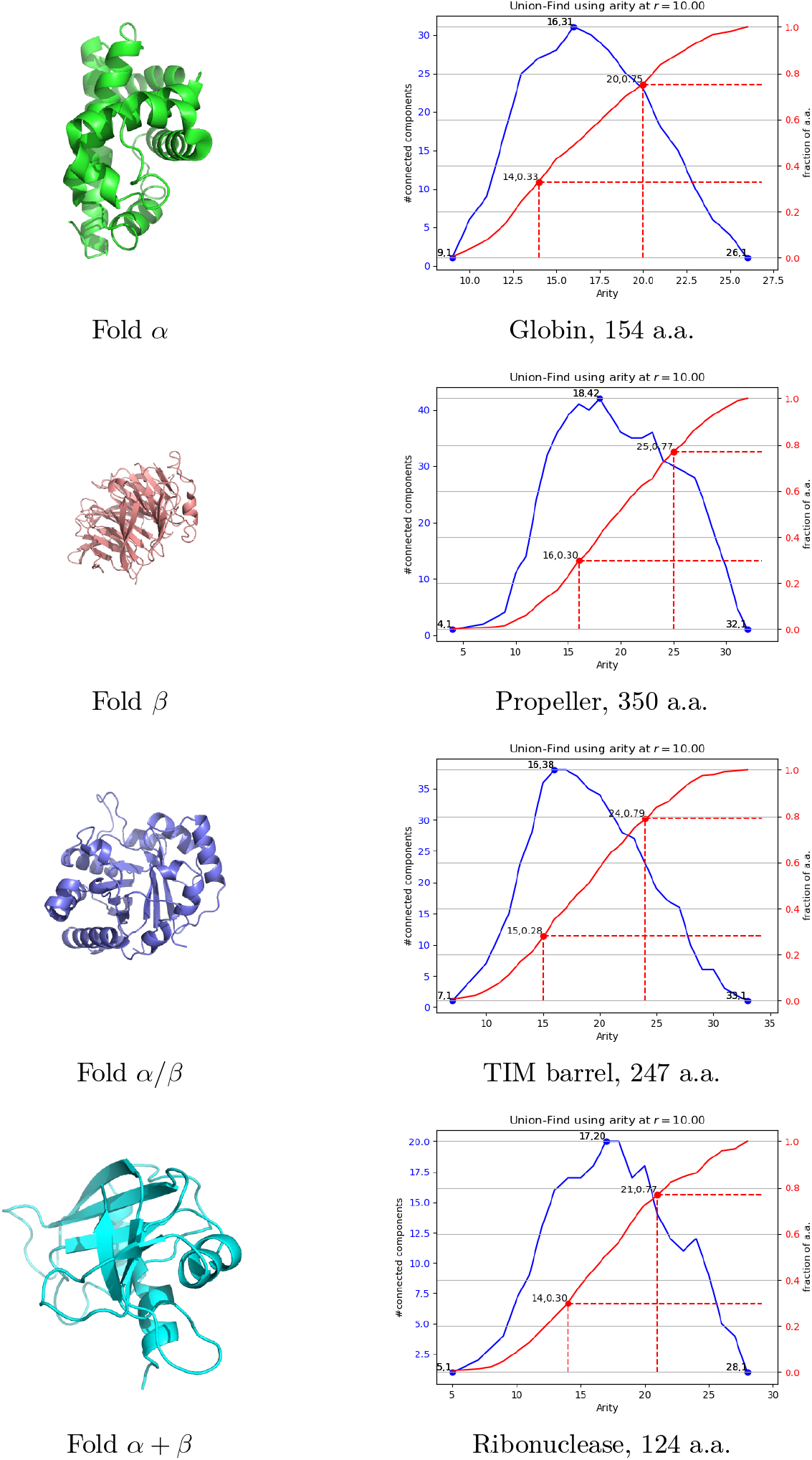
Arity distribution and arity signature for prototypical folds. Fold *α*: 101m-globin-alpha; 154 amino acids. Quantile-arity signature [(0.25, 14), (0.5, 17), (0.75, 20)] Fold *β*: 1erj-propeller-beta; 350 amino acids. Quantile-arity signature [(0.25, 16), (0.5, 20), (0.75, 25)] Fold *α/β*: 8tim-TIMbarrel-alphaSlashBeta; 247 amino acids. Quantile-arity signature [(0.25, 15), (0.5, 19), (0.75, 24)] Fold *α* + *β*: 1a5p-ribonuclease-alphaPlusBeta; 124 amino acids. Quantile-arity signature [(0.25, 14), (0.5, 17), (0.75, 21)]

**Figure 2:**
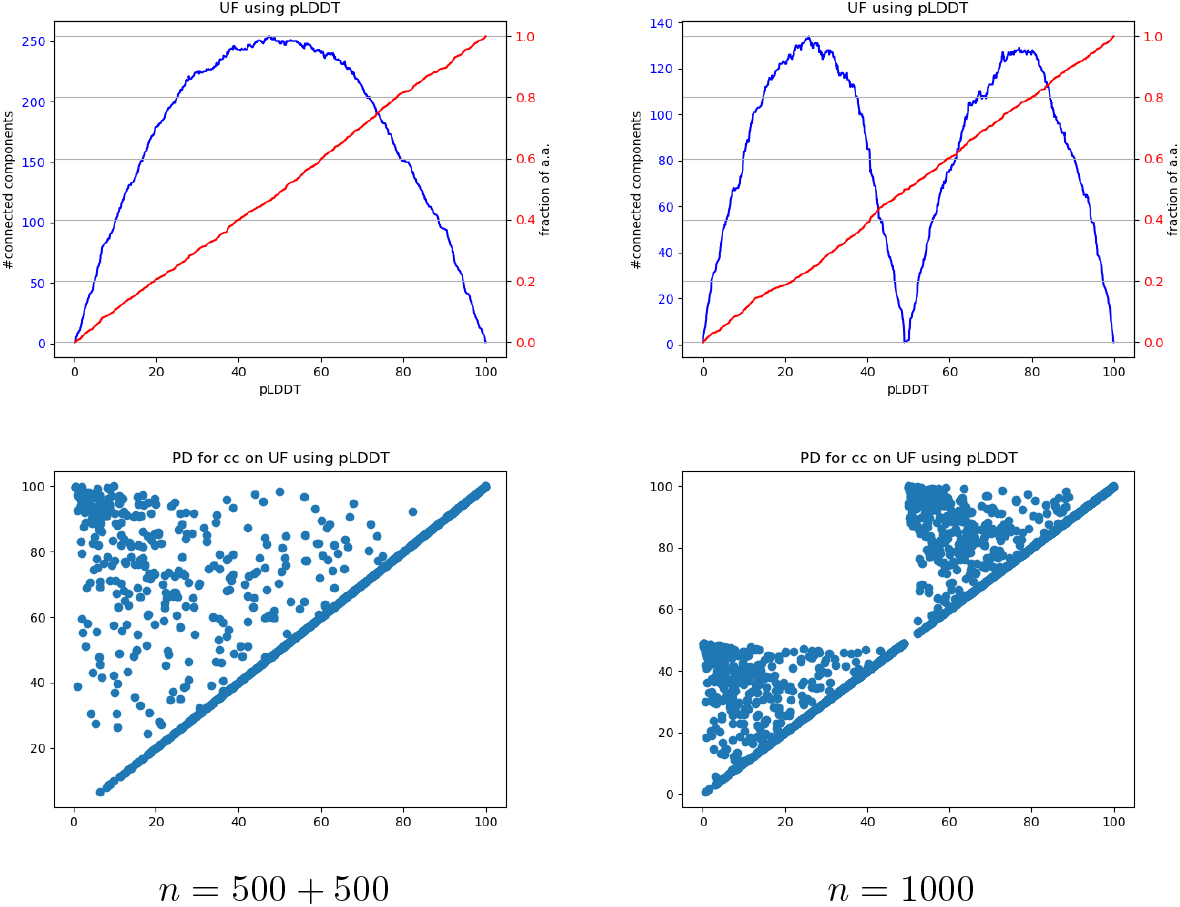
Filtration 𝒢_pLDDT_ and Union-Find: evolution of the number of connected components *N*_cc_(pLDDT) for fictitious proteins with one domain and two domains, respectively. Left: the *n* = 1000 a.a. are assigned random pLDDT values in [0, 100]; Right: the first 500 a.a. are assigned random pLDDT values in [0, 49], and the last 500 a.a. are assigned random pLDDT values in [50, 100]. **(Top row)** Blue curve: function *N*_cc_(pLDDT); red curve: fraction of amino acids. **(Bottom row)** persistence diagrams.

For a collection of structures, typically all predictions in a genome, we define:

#### Definition. 2

*Given two quantiles q*_1_(= 0.25) *and q*_2_(= 0.75), *the* arity map *is the map whose x and y axis are the arities at q*_1_ *and q*_2_. *A* bin/cell *of the map hosts all structures with a prescribed arity signature*.

Upon processing the *C*_*α*_ carbons by increasing arity, we are also interested in the number of connected components of the primary structure defined by all amino acids / *C*_*α*_s whose arity is less than a given threshold *A*_*i*_. We now formalize this idea using topological persistence.

### 2.2 Persistence based analysis on the primary structure

#### Filtrations and persistence diagrams

We aim to evaluate the *coherence*, with respect to a parameter generically denoted as *u*, of the *n* amino acids of a protein sequence in an `AlphaFold` prediction. Practically, *u* will be the arity or the pLDDT, so that we expose the method for a general value *u*.

We consider the polypeptide chain as a path graph with *n* vertices and *n* − 1 edges. From a topological data analysis perspective [18], we use a filtration 𝒢_*u*_ encoding a sequence of nested subgraphs of this path graph. Here, the calligraphic letter 𝒢 indicates that the filtration is applied to the path graph, and *u* is the value carried by each each amino acid. Upon inserting a.a. by increasing *u* value, we connect the a.a. being processed to its neighbors along the sequence if already inserted–Algorithm 1. The filtration starts with the empty graph and ends with the complete path graph.

As *u* varies, we track the number of connected components (c.c.) in the graph 𝒢 _*u*_ using the Union-Find algorithm [30]. This results in a function *ν* = *N*_cc_(*u*) representing the number of connected components at each value of *u*, with *ν* in the interval [1, ⌈*n/*2⌉].

The *persistence* of a connected component in 𝒢 _*u*_ is the time elapsed between its birth and the moment when it merges with a c.c. that was created earlier. The *persistence diagram* (PD) is the diagram whose *x*-axis (resp. y-axis) is the birth (resp. death) date [18]. The points of the PD are denoted *P* = {*c*_1_, …, *c*_*m*_}, and referred to as *critical points*. The persistence of point *c*_*i*_ reads as `p`_𝒢_ (*c*_*i*_) = death_𝒢_ (*c*_*i*_) − birth_𝒢_ (*c*_*i*_). We also denote *m*^*′*^ the number of critical points whose persistence is positive–*i*.*e*. critical points away from the diagonal, and define the fraction of positive critical points as 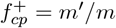.

Assuming that the persistences belong to a discrete set, we denote 𝒫 [·] the probability mass function ℙ: 𝒫 → [0, 1]. To assess the diversity of persistence values, we consider the normalized entropy of this probability distribution.

Finally, we consider the *salient / persistent* local maxima of the function *N*_cc_(*u*). We identify such maxima using persistence again on the filtration ℋ_*ν*_ associated with super-level sets 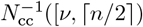 of the function *N*_cc_(*u*). (The calligraphic letter ℋ indicates that the filtration is on the height function *N*_cc_(*u*).) The persistence `p` (*c*_*i*_) = death_*ℋ*_ (*c*_*i*_) − birth_*ℋ*_ (*c*_*i*_) of a local maximum of *N*_cc_(*u*) is the elevation drop leading to a saddle point also connected to a more elevated local maximum. We simplify the function *N*_cc_(*u*) using the persistence simplification based on the Morse-Smale-Witten chain complex [31]. In particular, we denote PLM(*t*_*ν*_) the number of persistent local maxima of *N*_cc_(*u*) at persistence threshold *t*_*ν*_. In practice, we will use a relative threshold *t*_p_ ∈ (0, 1) to define *t*_*ν*_ = *n* * *t*_p_ – with *n* the number of amino acids.

We summarize the previous discussion with the following:

##### Definition. 3

*Using the filtration 𝒢*_*u*_ *coding subsets of the path graph connecting the n amino acids of a polypeptide chain, we define:*

- *The* fraction *of positive critical points:* 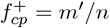.
- *The* mean persistence: 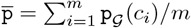,
- *The* normalized persistence entropy:

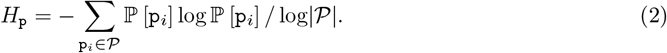

*Using the filtration* ℋ_*ν*_ *consisting of super-level sets of the function N*_*cc*_(*u*), *we define:*

- *The maximum number of connected components observed:* 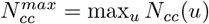
- *The number of persistent local maxima of the function N*_*cc*_(*u*) *at persistence threshold t*_*ν*_: *PLM*(*t*_*ν*_).

We are now in position to define the two filtrations of interest.

#### Arity based filtration 𝒢_arity_

Using the arity as filtration parameter yields the filtration 𝒢_arity_, whose connected components are maintained using Algorithm 2–a simple variant of Algorithm 1 handling all *C*_*α*_ carbons with a given arity in a batch.

As an illustration, we consider the evolution of connected components for prototypical folds (Fig. 1).

#### pLDDT based filtration

The filtration 𝒢_pLDDT_ uses –pLDDT as parameter *u*. We indeed process pLDDT values in increasing order, to handle high quality regions first. Assuming that all –pLDDT values are different, the PD therefore contains exactly *n* −1 critical points corresponding to the *n*− 1 edges connecting the *n* residues. A number of these have a null persistence and correspond to *accretion*–a *C*_*α*_ carbon inserted in the Union-Find data structure, and merged immediately to a neighbor on the left or the right. These account for the definition of the fraction 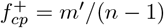 (Def. 3).

##### Example 1

*Consider first a fictitious protein with random pLDDT values in* [0, 100]. *The insertion of individual a*.*a. results in a well behaved curve peaking at the median pLDDT value, and whose maximum is conjectured to be 1/4 of the number of a*.*a. (Fig. 2, Fig. S1; conjecture 1). The persistence diagram also exhibits a characteristic triangular shape with a concentration of critical points near its apex, corresponding to the death of connected components upon moving forward to the full path graph in* G_*pLDDT*_. *Simulations also give the following approximate values:* 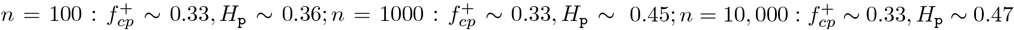.

*Consider next the following fictitious protein: (i) a first group of a.a. corresponding to an unstructured region on the N-ter side, with low confidence (say pLDDT* ∈ [0, 49]; *and (ii) a second group of a*.*a. corresponding to a structured region on the C-ter side, with high confidence (say pLDDT*∈ [50, 100]*). Along the insertion, one observes the formation of the former and the latter regions (Fig. 2)*.

## 3 Results

### 3.1 Q1. Contacts, packing, and whole genome predictions

#### Charting whole genomes with the arity map

The arity is instrumental at distinguishing ordered, disordered, and mixed proteins. While proteins with classical *α* and *β* folds exhibit an arity in the range 15-20, a mode in the interval 5-7 is observed for unstructured regions (Fig. 1, Fig. 3). See also Sec. S1.2 for numerous additional examples.

**Figure 3:**
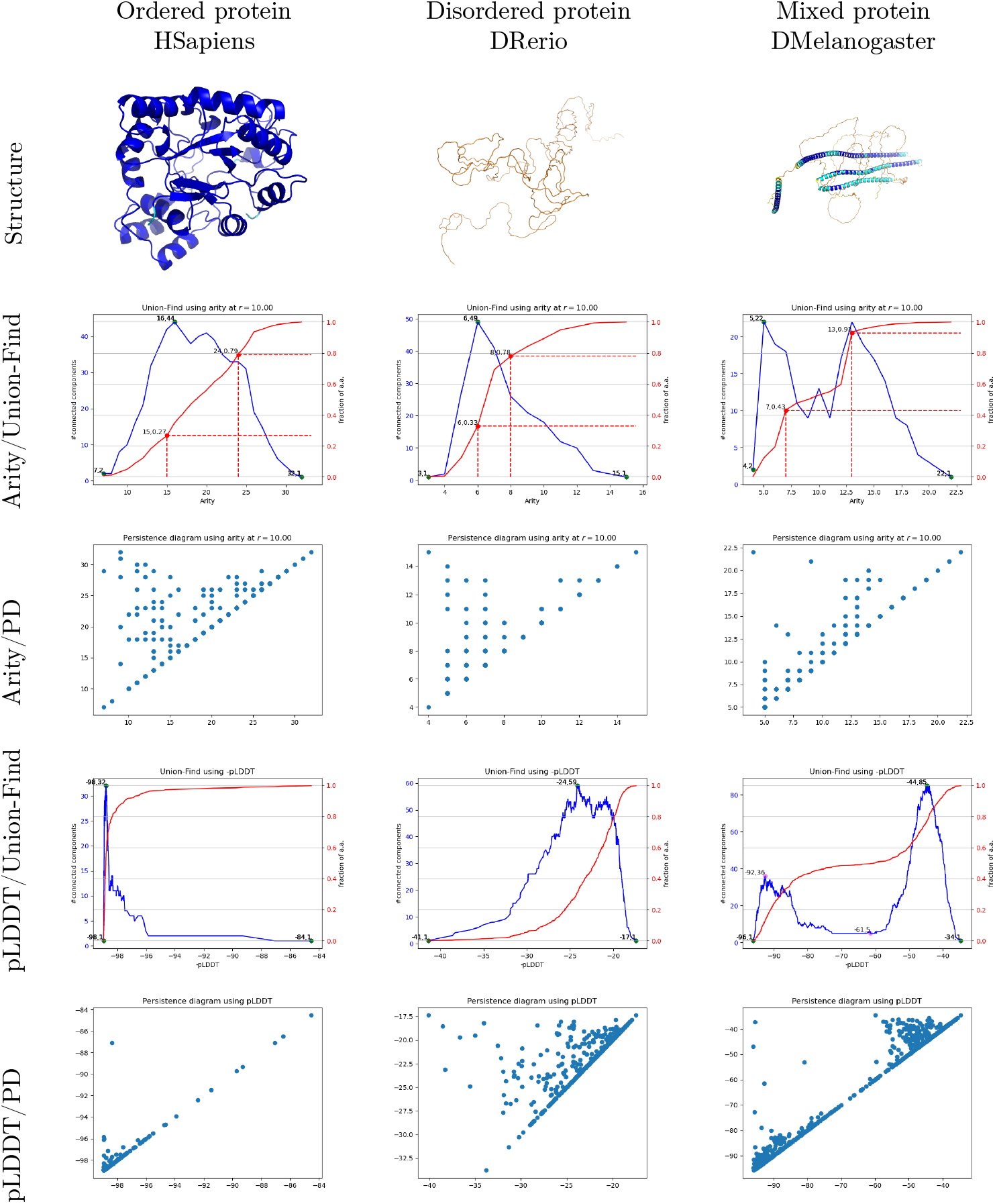
Filtrations 𝒢_arity_ and 𝒢_pLDDT_ on three prototypical examples corresponding to ordered, disordered and mixed proteins: evolution of the number of connected components and persistence diagrams. **(Left)** (H. Sapiens; AF-P15121-F1-model_v4; 316 amino acids; *H*_p_ ∼ 0.04.) **(Middle)** (Zebrafish; AF-A0A0G2L439-F1-model_v4; 449 amino acids; *H*_p_ ∼ 0.27.) **(Right)** (D. Melanogaster; AF-Q9VQS4-F1-model_v4; 781 amino acids; *H*_p_ ∼ 0.28.) Second row, 𝒢_arity_: the blue curve codes the variation of the number of connected components; the red one is the cumulated fraction of *C*_*α*_ carbons whose `arity` is less than a threshold. Third row, 𝒢_arity_: persistence diagram for the connected components. Fourth row, 𝒢_pLDDT_: same as the second row, using pLDDT instead of `arity`. Fifth row, 𝒢 _pLDDT_: same as the third row, using pLDDT instead of `arity`.

We process all proteins of a given genome and build the *arity map* (H. Sapiens, Fig. 4(A)). The interpretation of this map provides a whole genome analysis at a glance. The *x* (resp. *y*) axis `arity`_*C*_*α*_(*q*_1_) (resp. `arity`_*C*_*α*_(*q*_2_)) is geared towards loosely (resp. tightly) packed regions. For a given cell of this map, the distance to the diagonal *y* = *x* encodes the coherence of the overall packing of the region of the protein selected by the quantiles *q*_1_ and *q*_2_. For example, for bins/cells on the diagonal, a unique arity values suffices to move from a fraction of amino acids *< q*_1_ to a fraction *> q*_2_, stressing the coherence of the overall 3D structure. This is never the case for native protein structures, which always present different quantiles *q*_1_ and *q*_2_ (Fig. S12, in red) due to their interchange between rigid and flexible regions. But such a collapse is observed on `AlphaFold` predictions (Fig. 4, AF-Q8IWJ2-F1).

**Figure 4:**
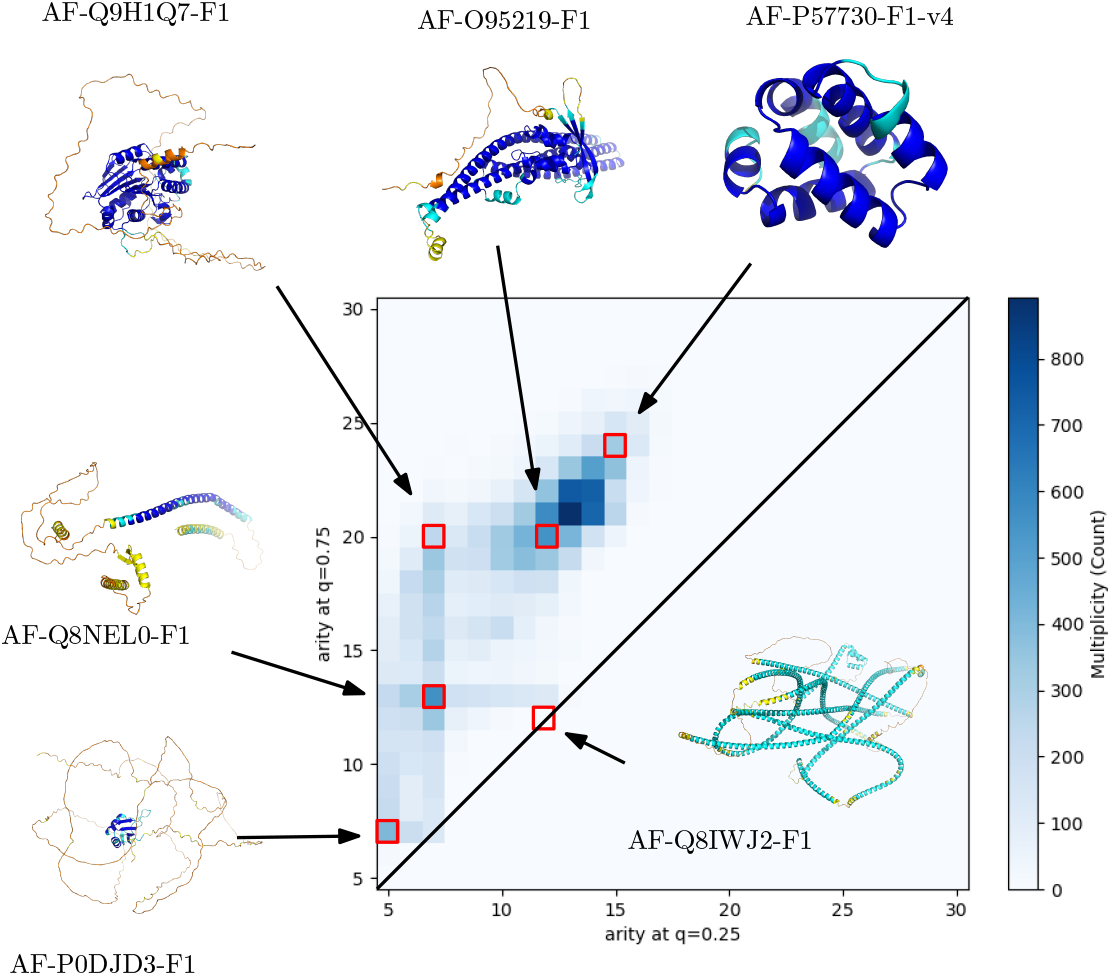
H. Sapiens: arity map discriminates well-folded, intrinsically disordered, and low-confidence structures. Upper triangle of the arity map: protein count with for a prescribed arity signature. Each bin corresponds to all `AlphaFold` predictions harboring a given arity signature.

**Figure 5:**
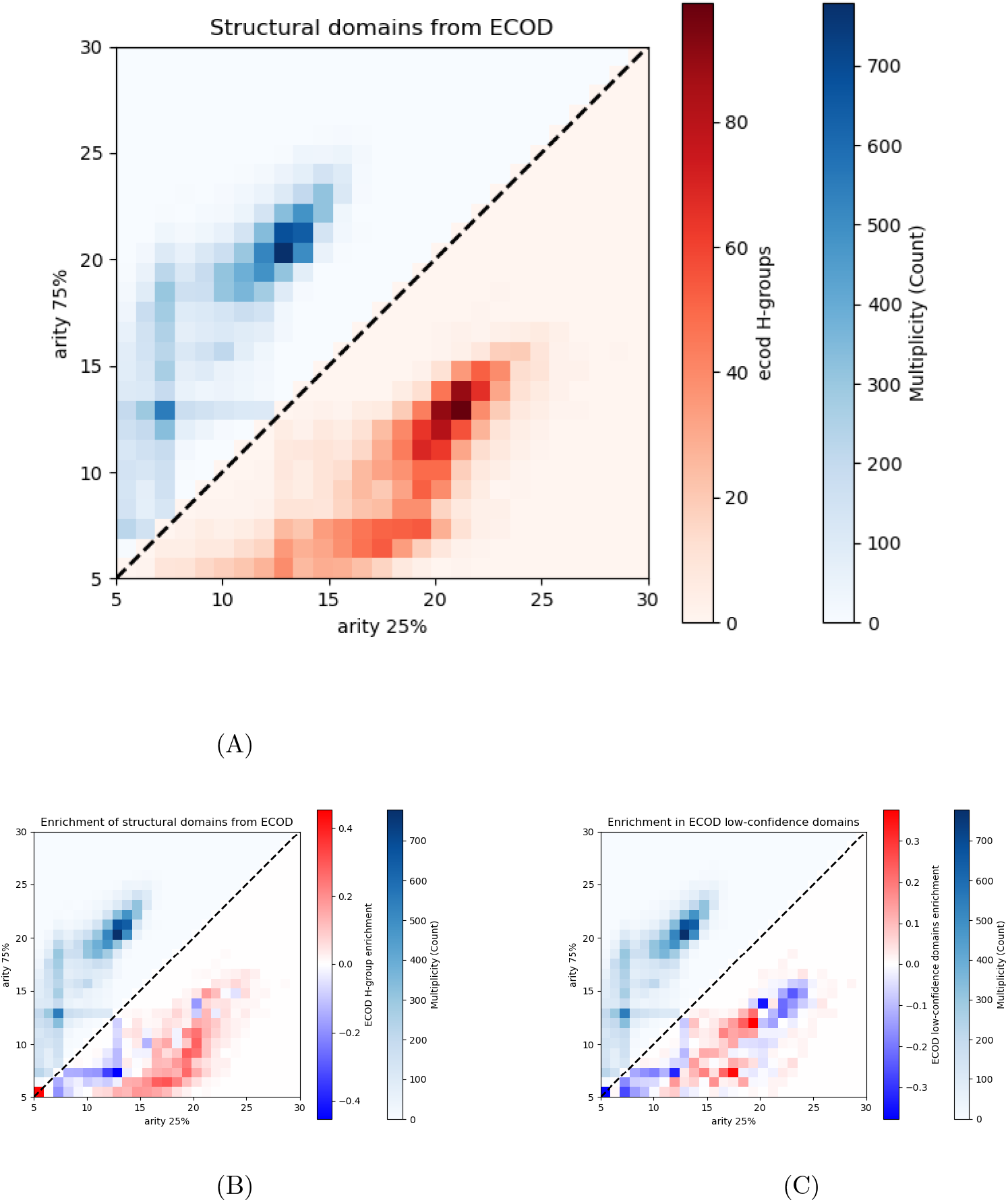
H. Sapiens: arity map of the human proteome vs ECOD domain enrichment in human proteins. The upper triangle always reports the human proteome arity map for reference. **(A)** Number of unique ECOD H-groups for each arity signature. **(B)** Enrichment difference (with respect to the proteome) of unique ECOD H-groups. **(C)** Enrichment difference (with respect to the proteome) of non-redundant ECOD low-confidence domains.

To further analyse proteins in a cell of the arity map, we perform a hierarchical clustering of arity histograms using the Earth Mover distance as metric [20, 28]. As expected, the clustering unveils the finer structure of those proteins (Fig. S10(B,C,D)).

We produced the arity map for all models genomes from `AlphaFold-DB` (H. Sapiens, Fig. 4; other species, Sec. S5). In passing, we note that arity and pLDDT strongly correlate (Fig. S11), confirming that the arity is also a good proxy for prediction robustness. Yet, pLDDT is not able to show the finer details of the structural repartition that emerges from the arity map.

These maps also appear to be bimodal. The most pronounced mode corresponds to well folded proteins, in the arity map region [14, 18] × [20, 25]. The second mode corresponds to the region [4, 8] × [10, 15], with the particularly populated cell 7 ×13. These modes are in general separated by low density regions. The inspection of genomes beyond the model ones yields different patterns, though. For example, *P. Falciparum* exhibits a continuum between the previous two modes, which is consistent with the overall low quality of predictions noted above.

We further investigate the hot spot at 7 ×13 of the human genome (Fig. S10). Contrary to the rest of the [4, 8] × [10, 15] region, this hot spot does not find any counterpart in the arity plot of the human proteins of known structure (Fig. S12, in red) – and is not present either in the plot of experimentally-confirmed disordered proteins (Fig. 6, discussed later). These `AlphaFold-DB` predictions therefore represent disordered structures or inaccurate predictions.

**Figure 6:**
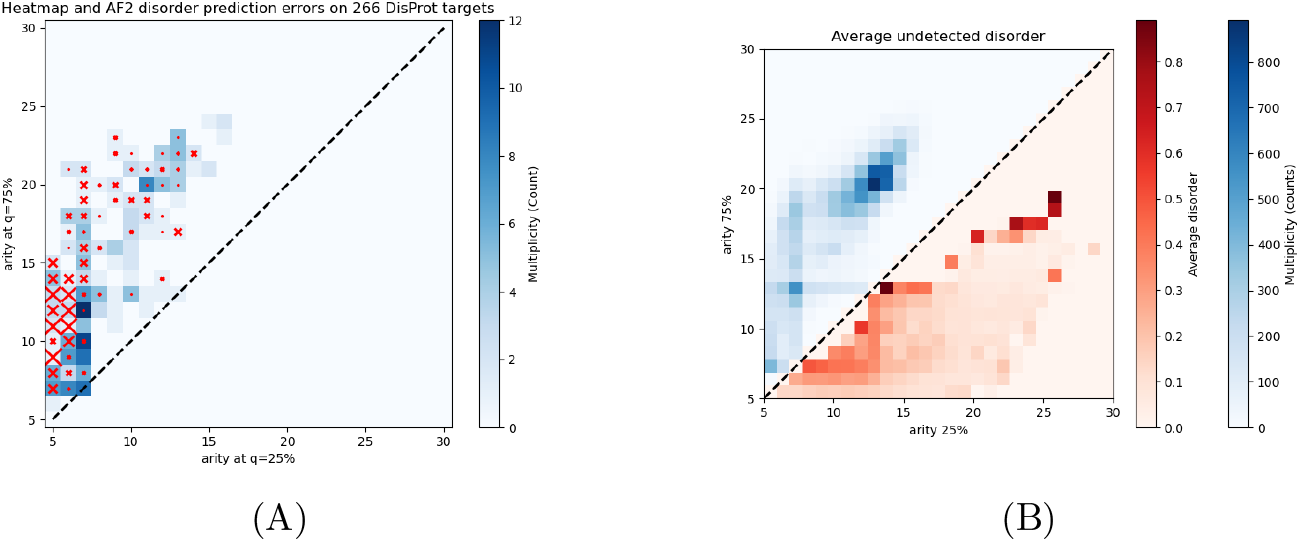
H. Sapiens: arity map discriminates IDRs. **(A)** `DisProt` **analysis on 266 target proteins lacking structural data**. The size of the red crosses over the arity map indicates the fraction of amino acids incorrectly predicted as disordered–pLDDT *<* 50% but lacking a disorder annotation in `DisProt`. (No cross: never, pixel-size cross: more than 70% of the times). The leftmost region of the plot shows a high enrichment in such cases. **(B)** `AIUPred` **analysis**. The upper triangle reports the human proteome arity for reference.

The connexion with IDRs/IDPs is the object of our Question 3.

#### Arity based filtration

The variability of arities coding the compactness of the atomic environments around *C*_*α*_s allows incrementally *building* the protein by inserting the a.a. by order of increasing arity and connecting each a.a. to its predecessor (N-ter) and successor (C-ter) if already inserted (Section 2.1, Algorithm 2).

The case of well folded proteins follows this pattern as focusing on salient extrema of the function *N*_cc_(`arity`) yields a unimodal arity curve with a salient extremum in the range 15-20 (Fig. 3(Left column, second row, blue curve)). The same holds for unstructured proteins, excepts that the local extremum of *N*_cc_(`arity`) is attained at values ∼5 − 7 (Fig. 3(Middle column, second row, blue curve)). The signature is more informative for proteins having both high and low quality regions (Fig. 3(Right column, second row, blue curve)). For such proteins, the curve of connected components is typically bimodal, since one successively forms the regions with low and high arity.

Concluding, the ability to detect the persistent local extrema on these curves gives a direct indication on the structural elements encountered in the predictions.

### 3.2 Q2. Predicted domains and their quality

We investigate potential learning biases in `AlphaFold` predictions at the sequence and structure levels.

At the sequence level, counting the number of homologous structures in `UniProtKB` for each arity signature exhibits a clear trend between the arity signature and the average number of homologous sequences (Section S1.2, Fig. S13(A)). However, mapping these homologs to their cluster representatives in the current release of `Uniclust30` erases this bias (Fig. S13(B)), simply showing that structured proteins have more homologous sequences in nature. We also note that the quality of predictions in terms of average pLDDT shows a gradient towards high arity values (Fig. S12). Since both the `AlphaFold` direct and distillation training algorithms construct MSAs of non-redundant sequences with sequence identity *<* 30%, we conclude that a large and diverse input MSA is necessary but not sufficient for guaranteeing high-quality predictions.

We proceed with the investigation of potential biases at the structural level using the recently published `ECOD` structural domain classification of the entire human proteome [25]. Using `AlphaFold-DB` predictions, `ECOD` could enrich its classification with several new structural domains. At the level of H-groups in `ECOD` hierarchy, each identifier refers to a cluster of sequence-homologous, same-topology domains in tight relation with the SCOP superfamily level. For each arity signature, we thus collect the unique `ECOD` H-group identifiers and the corresponding folds. The resulting 2D-histogram shows a clear correlation with the protein count and the number of `ECOD` folds (Fig. S14(A), upper triangle for count and lower triangle for `ECOD` folds). In order to remove the correlation and study the residual signal, we consider the signed difference of the two rescaled histograms. (The rescaling ensures that the two maxima are equal.) This difference map shows two main domains (Fig. S14(B), red and blue regions). On the one hand, the region of the main peak *i*.*e*. 13 × 20 until the 7 × 13 peak (excluded) shows an enrichment in `ECOD` domains. On the other hand, the region of the 7× 13 peak towards the bottom left of the arity map (IDP region, see next Section) gets depleted.

`ECOD` also provides a set of low-confidence domains associated with low pLDDT and PAE (predicted aligned error) values [25], enabling the calculation of the enrichment relative to this dataset. Remarkably, the former *red* region gets split, the enrichment in low confidence domains being confined to the 7 × 13 signature as well as the arity signatures [5, 15] × [14, 20]. This extended region contains structures richer in IDRs, but still having well-formed domains. Conversely, the [5, 7] × [5, 12] region is poor both in standard and low-confidence well-formed domains, hinting at a complete dominance of IDRs over structured regions. These observations call for a further inspection of IDPs/IDRs in the arity map.

### 3.3 Q3. Predictions and intrinsically disordered proteins/regions

Several recent works assume that in `AlphaFold`’s predictions, regions with low pLDDT and/or not displaying secondary nor tertiary structure are to be considered intrinsically disordered rather than just the product of insufficient or unexploitable information. The tools developed in the previous sections can help assessing whether and when a correlation between unstructured regions in `AlphaFold`’s prediction correlate with experimental evidences of disordered regions.

#### Analysis using DisProt

We first consider the `DisProt` database, which collects disordered regions from more than 1300 target protein sequences. Many of the benchmarked sequences correspond to proteins of known structure, present in the PDB. For this test, we focus on the 266 targets lacking structural data and that have a predicted structure on `AlphaFold-DB`. Consider all a.a. of all proteins in a given arity map cell. We compute the fraction of false positive of these – a.a. with a pLDDT *<* 50% but lacking a disorder annotation in `DisProt`. Notably, `DisProt` is known not to provide complete annotations, thus in principle `AlphaFold` could discover new unannotated IDRs on the `DisProt` targets. The simplest hypothesis consists of expecting an incompleteness proportional to the extent of the predicted disordered regions–regardless of other structural characteristics of `AlphaFold` predictions. However, while `AlphaFold` predicts a large number of amino acids as part of IDRs for the region of the arity map `arity`_*C*_*α*_(0.25) *<* 7, most of these are not annotated as such in `DisProt` (Fig. 6(A)). Predictions populating that part of the arity plot are thus likely to be of poor quality: this is not surprising, as the previous section details that proteins in this region often have a low number of homologous structures.

#### Analysis using AIUPred

Finally, we reconsider the 7× 13 cell of the arity map of the human genome, hosting candidates for divergent `AlphaFold` predictions (Fig. S10). We ran the state-of-the-art software `AIUPred` [32] to predict IDRs on the whole human genome, and for each protein computed the number of residues characterized by pLDDT ≥ 0.5 and a disorder score of *>* 0.5 according to `AIUPred` (Fig. 6(B)). The triangle delimited by ([6, 8], [6, 13], [12, 13]) is enriched in predicted structures not clearly recognized by `AlphaFold`. This region is somehow complementary to the one highlighted by `DisProt`, and interestingly, the only neighborhood that both displays false positive and false negative IDR predictions is centered in 7 × 13.

### 3.5 Q4. pLDDT values and fragmentation of AlphaFold reconstructions

We use the filtration 𝒢_pLDDT_ and the associated values (Def. 3, Fig. 3, Fig. 7) to characterize the coherence of pLDDT values along the backbone.

**Figure 7:**
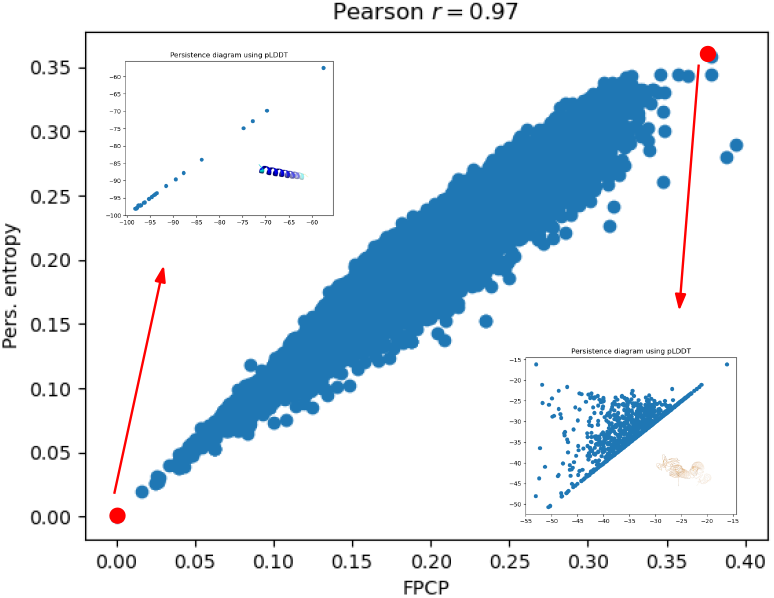
The persistence entropy *H*_p_ and the fraction of positive critical points (FPCP) 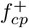 of 𝒢_pLDDT_ show a continuum between highly coherent and random-like structures in terms of pLDDT. Illustrations for HSapiens, with insets displaying the structures achieving the minimum and maximum *H*_p_ values.

#### Specific proteins with different profiles

To appreciate the understanding yielded by our statistics, and in particular the persistence entropy *H*_p_ and the variation of the number of connected components *N*_cc_(pLDDT), let us consider various protein profiles, namely well structured, disordered, and mixed (Fig. 3). It appears that the function *N*_cc_(pLDDT) varies from unimodal (when considering persistent maxima) to multimodal. The former occurs for well structured or unstructured proteins (Fig. 3(Left, Middle)), and the latter for mixed cases (Fig. 3(Right)). For example, the *D. Melanogaster* protein AF-Q9VQS4-F1-model_v4 exhibits a clear bimodal profile with two persistent maxima (Fig. 3(Right)). On this example, the plateau at five connected components corresponds to five helices–four long ones plus a shorter one. The persistence diagram associated with the construction of the filtration also exhibit the dynamics of connected components (Fig. 3, fourth row). Using the same example, the PD of *D. Melanogaster* quantifies the dynamics of the high (resp. low) confidence region of the molecule on the left (resp. right) part of the diagram. We note that the apex of the triangles are not as populated as those of the null model (Fig. S1), showing a lesser randomness of pLDDT values, which is consistent with the lesser entropy *H*_p_.

Let us now analyse these quantities in more detail at the genome scale.

#### Persistence entropy

At the genome scale, we observe a strong correlation between the three pairs of variables amidst 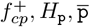 (Fig. 7, Fig. S15). The maxima of 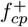 are consistent with those a the null random model, while those of *H*_p_ are slightly below those of the same null model. (NB: average values of p are irrelevant for the null model.) For all organisms, the minimum *H*_p_ is observed for very regular structures for which all pLDDT values are within a small range (Fig. S16(Second row)). Instead, the maxima are observed for highly disordered structures whose a.a. span a wide range of pLDDT values (Fig. S16(Third row)). For such cases, the typical triangular shape in the PD is also observed. In considering the overall pLDDT statistics per organism (Table 1, SI Section S6), it appears that shifting the pLDDT values towards 100 results in a lesser fraction of positive critical points and in a smaller entropy (Fig. S15).

#### Maximum value 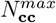 **of the function** *N*_**cc**_(**pLDDT**)

The maximum number of local maxima observed *N*^*max*^ normalized by the protein size (Fig. S17) in general lower than that of the null model – conjectured to be equal to 1*/*4, even though some cases reach or even surpass that bound. This is accounted for by two reasons. First, accretion is at play–see definition above. Second, pLDDT values come in batches, so that local insertions into *stretches* along the sequence prevent the maximum number of local maxima to rise. These batches are identified by the persistent local maxima discussed next.

#### Persistent local maxima PLM of the function *N*_**cc**_(**pLDDT**)

We first note that the number of persistent local maxima is in general less than 10 (Fig. S15(Third column)). Also, the correlation between *H*_p_ (or 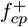)and PLM is quite low, which is expected. One can indeed have a significant number of persistent critical points localized in a narrow range of pLDDT values (Fig. S18). This phenomenon is well known to geographers defining peaks on mountains: two tall peaks can be separated by a low lying pass, but be located at a short flying distance [33]. Nevertheless, as pointed out above, persistent local maxima identify coherent stretches along the sequence in terms of pLDDT values.

#### Model assessment and database queries

In the model assessment perspective, querying the encoding provided by the number of a.a., the persistence entropy *H*_p_, the fraction of positive critical points 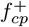 and the number of persistent local maxima PLM enables the identification of cases incurring a large fragmentation. Such cases exhibit a wide range of pLDDT values aggregated in coherent stretches along the sequence, yielding a large entropy and a (relatively) high number of persistent local maxima of the function *N*_cc_(pLDDT) (Fig. 8).

**Figure 8:**
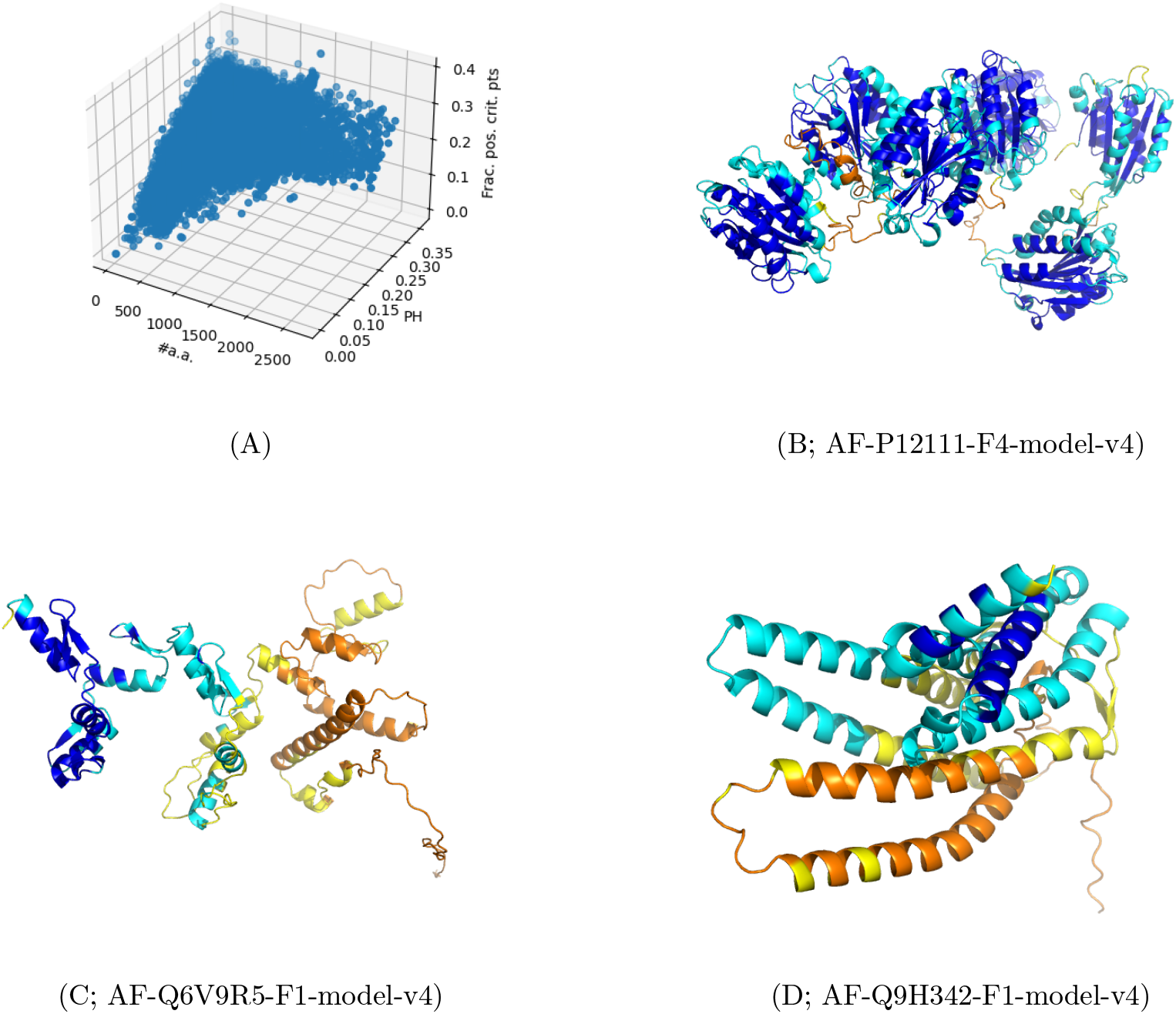
Illustration of model fragmentation: wide range of pLDDT values aggregated in coherent stretches of pLDDT values along the sequence. **(A)** Scatter plot with protein size persistence entropy *H*_p_ × fraction of positive critical points 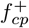.**(B, C, D)** For HSapiens, at persistence threshold *t*_p_ = 0.025, 86 structures are characterized by *H*_p_ ≥0.25, #*a*.*a*. ≥200, PLM ≥3. Three of them are displayed.

## 4 Discussion and outlook

`AlphaFold` has significantly advanced protein structure prediction, delivering remarkably high accuracy across a wide range of targets. A key feature of its predictions is the pLDDT score, which offers an estimate of local confidence for each amino acid of the predicted structure. This score is computed by `AlphaFold`’s structure module–a component that has since been widely adopted in other structure prediction tools. pLDDT has also been shown to anticorrelate with intrinsically disordered regions, surpassing the accuracy of dedicated state-of-the-art disorder predictors. Despite its impressive overall performance, `AlphaFold` is not without flaws: there have been consistent reports of clear prediction errors–cases where incorrect structures are assigned high average pLDDT scores [34, 35, 36]–yet identifying a shared cause behind these failures has remained elusive. Only recently have large-scale, first-principles analyses begun to emerge for protein language models (pLMs) [37].

In this context, our work introduces a novel representation of structural elements in proteins with two powerful descriptors at the amino acid level, which are formalized as topological filtrations.

The first filtration describes the 3D packing properties and the corresponding cumulative distribution counting the number of *C*_*α*_ carbons with a prescribed number of neighbors. This distribution facilitates the comparison of structures using optimal transportation, and the persistent components of the filtration can be used to compare structural motifs from different proteins. To simplify this representation, the distribution can be reduced to two integers, forming the *arity map*, which simultaneously performs clustering of coherent structures and dimensionality reduction. The assignment of structures to cells of the arity map charts the space of predictions from the structural standpoint, and provides an embedding of structures that enables the study of certain properties *as a function* of the arity. In particular, the coherence of observations for neighboring cells being unveils important trends in two directions. First, the study of domains yields the identification of a specific region of the arity map populated with low quality structures. The region is enriched in low-confidence domains recently published in the `AlphaFold` enriched version of the `ECOD` database. Most of these domains may be termed *hallucinations* since they may appear valid, but are incorrect and/or fabricated. Second, the investigation of IDRs/IDPs using the arity map also hints at another specific region with a high rate of false positive IDRs. While the performances of `AlphaFold` for IDP prediction have been praised, the notion of isolated structure for IDPs is iconoclastic since dynamics play a key role, and future work will likely address the generation of ensembles.

The second filtration uses pLDDT values along the sequence. This filtration and the associated statistics characterize cases with a wide range of pLDDT values aggregated in coherent stretches along the sequence, yielding a large entropy and a (relatively) high number of persistent local maxima of the function counting connected components. These analysis make it possible, in particular, to isolate regions of a model with a coherent quality (pLDDT) profile.

Overall, we believe that the statistics developed in this work complement those already delivered within `AlphaFold-DB`, and will contribute to understand limitations of `AlphaFold` regarding low quality domains and false positive IDRs.

## Acknowledgments

This work has been supported by the French government, through the 3IA Côte d’Azur Investments (ANR-19-P3IA-0002), and the ANR project Innuendo (ANR-23-CE45-0019).

We wish to thank V. Mallet for insightful comments.

## References

[1] J. Jumper, R. Evans, A. Pritzel, T. Green, M. Figurnov, O. Ronneberger, K. Tunyasuvunakool, R. Bates, A. Žídek, A. Potapenko, et al. Highly accurate protein structure prediction with AlphaFold. Nature, 596(7873):583–589, 2021.

[2] V. Mariani, M. Biasini, A. Barbato, and T. Schwede. lDDT: a local superposition-free score for comparing protein structures and models using distance difference tests. Bioinformatics, 29(21):2722–2728, 2013.

[3] Stefan Bienert, Andrew Waterhouse, Tjaart AP De Beer, Gerardo Tauriello, Gabriel Studer, Lorenza Bordoli, and Torsten Schwede. The swiss-model repository—new features and functionality. Nucleic acids research, 45(D1):D313–D319, 2017.

[4] Mihaly Varadi, Stephen Anyango, Mandar Deshpande, Sreenath Nair, Cindy Natassia, Galabina Yordanova, David Yuan, Oana Stroe, Gemma Wood, Agata Laydon, et al. Alphafold protein structure database: massively expanding the structural coverage of protein-sequence space with high-accuracy models. Nucleic acids research, 50(D1):D439–D444, 2022.

[5] Ashish Vaswani, Noam Shazeer, Niki Parmar, Jakob Uszkoreit, Llion Jones, Aidan N Gomez, łukasz Kaiser, and Illia Polosukhin. Attention is all you need. Advances in neural information processing systems, 30, 2017.

[6] F. Morcos, A. Pagnani, B. Lunt, A. Bertolino, D. Marks, C. Sander, R. Zecchina, J. Onuchic, T. Hwa, and M. Weigt. Direct-coupling analysis of residue coevolution captures native contacts across many protein families. PNAS, 108(49):E1293–E1301, 2011.

[7] Roshan M Rao, Jason Liu, Robert Verkuil, Joshua Meier, John Canny, Pieter Abbeel, Tom Sercu, and Alexander Rives. MSA transformer. In International Conference on Machine Learning, pages 8844–8856. PMLR, 2021.

[8] Bernard Moussad, Rahmatullah Roche, and Debswapna Bhattacharya. The transformative power of transformers in protein structure prediction. Proceedings of the National Academy of Sciences, 120(32):e2303499120, 2023.

[9] T. Hastie, R. Tibshirani, and J. Friedman. The Elements of statistical learning: data mining, inference and prediction. Springer, 2001. htf-esldm-01.

[10] Mikhail Belkin, Daniel Hsu, and Ji Xu. Two models of double descent for weak features. SIAM Journal on Mathematics of Data Science, 2(4):1167–1180, 2020.

[11] Qizhe Xie, Minh-Thang Luong, Eduard Hovy, and Quoc V Le. Self-training with noisy student improves imagenet classification. In Proceedings of the IEEE/CVF conference on computer vision and pattern recognition, pages 10687–10698, 2020.

[12] A. Maalouf, G. Eini, B. Mussay, D. Feldman, and M. Osadchy. A unified approach to coreset learning. IEEE Trans. Neural Networks Learn. Syst., 35(5), 2024.

[13] Janani Durairaj, Mehmet Akdel, Dick de Ridder, and Aalt DJ van Dijk. Geometricus represents protein structures as shape-mers derived from moment invariants. Bioinformatics, 36(Supplement_2):i718–i725, 2020.

[14] Mehmet Akdel et al. A structural biology community assessment of alphafold2 applications. Nature Structural & Molecular Biology, 29(11):1056–1067, 2022.

[15] Kathryn Tunyasuvunakool et al. Highly accurate protein structure prediction for the human proteome. Nature, 596(7873):590–596, August 2021.

[16] Oliviero Carugo. PLDDT values in AlphaFold2 protein models are unrelated to globular protein local flexibility. Crystals (Basel), 13(11):1560, November 2023.

[17] Damiano Piovesan, Alexander Miguel Monzon, and Silvio CE Tosatto. Intrinsic protein disorder and conditional folding in AlphaFoldDB. Protein Sci., 31(11):e4466, November 2022.

[18] H. Edelsbrunner and J. Harer. Computational topology: an introduction. AMS, 2010.

[19] Jean-Daniel Boissonnat, Frédéric Chazal, and Mariette Yvinec. Geometric and topological inference, volume 57. Cambridge University Press, 2018.

[20] Y. Rubner, C. Tomasi, and L.J. Guibas. The earth mover’s distance as a metric for image retrieval. International Journal of Computer Vision, 40(2):99–121, 2000.

[21] C. Villani. Topics in optimal transportation. Number 58. AMS, 2003.

[22] Antonina Andreeva, Dave Howorth, John-Marc Chandonia, Steven E Brenner, Tim JP Hubbard, Cyrus Chothia, and Alexey G Murzin. Data growth and its impact on the scop database: new developments. Nucleic acids research, 36(Suppl_1):D419–D425, 2007.

[23] Ian Sillitoe, Natalie Dawson, Tony E Lewis, Sayoni Das, Jonathan G Lees, Paul Ashford, Adeyelu Tolulope, Harry M Scholes, Ilya Senatorov, Andra Bujan, et al. CATH: expanding the horizons of structure-based functional annotations for genome sequences. Nucleic acids research, 47(D1):D280– D284, 2019.

[24] Hua Cheng, R Dustin Schaeffer, Yuxing Liao, Lisa N Kinch, Jimin Pei, Shuoyong Shi, Bong-Hyun Kim, and Nick V Grishin. ECOD: an evolutionary classification of protein domains. PLoS computational biology, 10(12):e1003926, 2014.

[25] R Dustin Schaeffer, Jing Zhang, Kirill E Medvedev, Lisa N Kinch, Qian Cong, and Nick V Grishin. ECOD domain classification of 48 whole proteomes from AlphaFold structure database using DPAM2. PLoS Comput. Biol., 20(2):e1011586, February 2024.

[26] Alderson, T. Reid and Pritisanac, Iva and Kolaric, Desika and Moses Alan M. and Forman-Kay, Julie D. Systematic identification of conditionally folded intrinsically disordered regions by alphafold2. Proceedings of the National Academy of Sciences, 120(44), October 2023.

[27] F. Cazals, J. Herrmann, and E. Sarti. Simpler protein domain identification using spectral clustering. Proteins: structure, function, and bioinformatics, NA(NA), 2025.

[28] Nicolas Bonneel, Michiel Van De Panne, Sylvain Paris, and Wolfgang Heidrich. Displacement interpolation using lagrangian mass transport. In Proceedings of the 2011 SIGGRAPH Asia conference, pages 1–12, 2011.

[29] Rémi Flamary, Nicolas Courty, Alexandre Gramfort, Mokhtar Z Alaya, Aurélie Boisbunon, Stanislas Chambon, Laetitia Chapel, Adrien Corenflos, Kilian Fatras, Nemo Fournier, et al. Pot: Python optimal transport. Journal of Machine Learning Research, 22(78):1–8, 2021.

[30] Thomas H Cormen, Charles E Leiserson, Ronald L Rivest, and Clifford Stein. Introduction to algorithms. MIT press, 2022.

[31] F. Cazals and D. Cohen-Steiner. Reconstructing 3D compact sets. Computational Geometry Theory and Applications, 45(1-2):1–13, 2011.

[32] Gábor Erdős and Zsuzsanna Dosztányi. AIUPred: combining energy estimation with deep learning for the enhanced prediction of protein disorder. Nucleic Acids Research, page gkae385, 2024.

[33] C. Thöni. Criteria to define summits in the Swiss alps: prominence and culminance height. Les Alpes, 1:26–28, 2003.

[34] Thomas C. Terwilliger, Dorothee Liebschner, Tristan I. Croll, Christopher J. Williams, Airlie J. McCoy, Billy K. Poon, Pavel V. Afonine, Robert D. Oeffner, Jane S. Richardson, Randy J. Read, and Paul D. Adams. Alphafold predictions are valuable hypotheses and accelerate but do not replace experimental structure determination. Nature Methods, 21(1):110–116, November 2023.

[35] Devlina Chakravarty, Myeongsang Lee, and Lauren L. Porter. Proteins with alternative folds reveal blind spots in alphafold-based protein structure prediction. Current Opinion in Structural Biology, 90:102973, February 2025.

[36] Jeffrey P. Bonin, James M. Aramini, Ying Dong, Hao Wu, and Lewis E. Kay. Alphafold2 as a replacement for solution nmr structure determination of small proteins: Not so fast! Journal of Magnetic Resonance, 364:107725, July 2024.

[37] Zhidian Zhang, Hannah K. Wayment-Steele, Garyk Brixi, Haobo Wang, Dorothee Kern, and Sergey Ovchinnikov. Protein language models learn evolutionary statistics of interacting sequence motifs. Proceedings of the National Academy of Sciences, 121(45), October 2024.

