## Supporting Information for "AlphaFold predictions on whole genomes at a glance: a coherent view on packing properties, pLDDT values, and disordered regions"

### S1 Methods

#### S1.1 Algorithms

**Algorithm 1 Union-Find on the polypeptide chain and filtration $\mathcal{G}_u$.** Particular case: using pLDDT as value for the parameter $u$ yields the filtration $\mathcal{G}_{\text{pLDDT}}$.

**procedure** BUILD_PATH_GRAPH_FILTRATION($[(j, u_j)]_{i=1,\ldots,n}$)
  Form the list $[(j, u_j)], j = 1, \ldots, n$, for the $n$ amino acids
  Let $L$ be this sorted list ascending $u_j$ values
  **for** $(j, u_j) \in L$ **do**
    UF.make_set($c_j$)
    if $j > 1$ and $c_{j-1}$ exists in the UF data structure: UF.union($c_j, c_{j-1}$)
    if $j < n$ and $c_{j+1}$ exists in the UF data structure: UF.union($c_j, c_{j+1}$)
    $cc_j \leftarrow$ UF.num_cc
    $nn_j \leftarrow nn_j + 1$
  **end for**
  **return** $\{(cc_i, nn_i)\}$ and the associated persistence diagram
**end procedure**

**Algorithm 2 Union-Find on the polypeptide chain using arities yields the filtration $\mathcal{G}_{\texttt{arity}}$.**

**procedure** BUILD_ARITY_FILTRATION($\{a_i\}_{i=1,\ldots,n}$)
  Compute all individual arities
  Compute the the sorted list $L = [A_1, \ldots, A_m]$ of unique arities
  **for** $A_i \in L$ **do**
    **for** $c_j \in A_{C_\alpha}^{-1}(A_i)$ **do**
      UF.make_set($c_j$)
      if $j > 1$ and $c_{j-1}$ exists in the UF data structure: UF.union($c_j, c_{j-1}$)
      if $j < n$ and $c_{j+1}$ exists in the UF data structure: UF.union($c_j, c_{j+1}$)
    **end for**
    $cc_i \leftarrow$ UF.num_cc
    $nn_i \leftarrow$ UF.num_nodes
  **end for**
  **return** $\{(cc_i, nn_i)\}$ and the associated persistence diagram
**end procedure**

#### S1.2 Methods: pLDDT filtrations for fictitious proteins

The simulations (Fig. 2, Fig. S1) suggest the following conjecture:

**Conjecture. 1** *For a $n$-nodes graph/path, the expectation of the maximum of the number of connected components yielded by the incremental construction of Algorithm 1 is equal to $n/4$.*

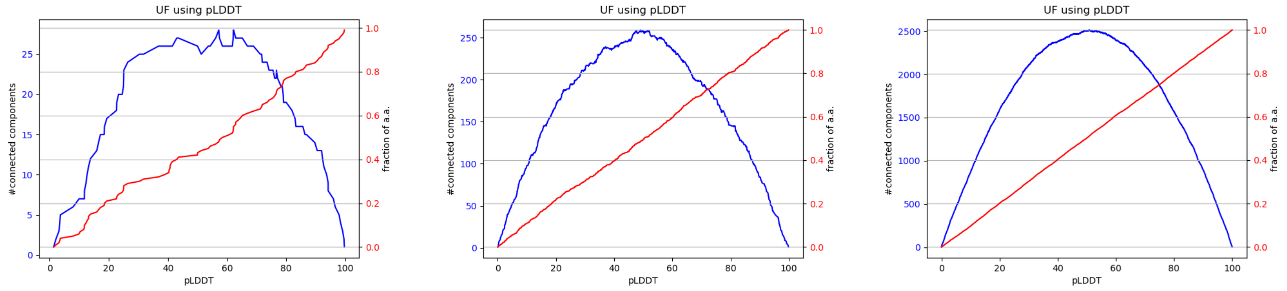

$n = 100$ $n = 1000$ $n = 10000$

Figure S1: **Union-Find and filtration $\mathcal{G}_{\mathbf{pLDDT}}$: evolution of the number of connected components $N_{\mathbf{cc}}(\mathbf{pLDDT})$ upon inserting $n$ values drawn uniformly at random in $[0, 100]$.** Conventions identical to Fig. 2.

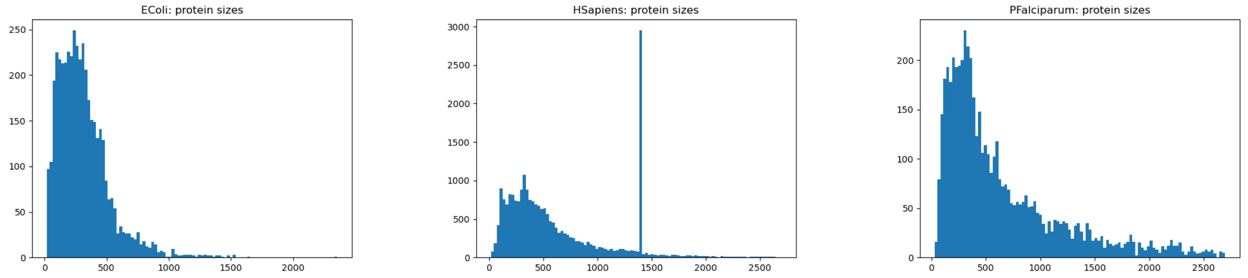

EColi HSapiens PFalciparum

Figure S2: **Distribution of protein sizes: illustration for three genomes.** For HSapiens, the peaks corresponds to the addition of sequences explained as follows "*... as well as 1400-residue fragments to cover longer human proteins*" [4].

### S2 Q1. Contacts, packing, and whole genome predictions

When the pLDDT values span a small range, the correlation with arity is in general low (structured protein, Fig. S4(B); unstructured protein, Fig. S6)(B). On the other hand, when the pLDDT spans a large range of values, from very low to very high, a significant Pearson correlation coefficient is generally observed (see examples Fig. S5(B), Fig. S7(B), Fig. S8(B), Fig. S9(B)).

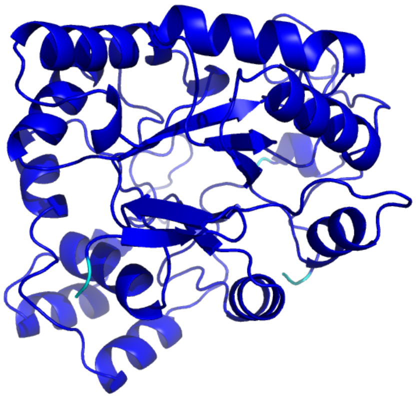

(A) 3D view

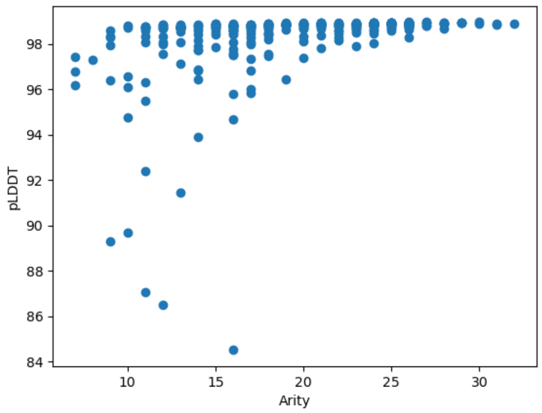

(B) Arity at $r = 10$ x pLDDT. Pearson $r = 0.36$

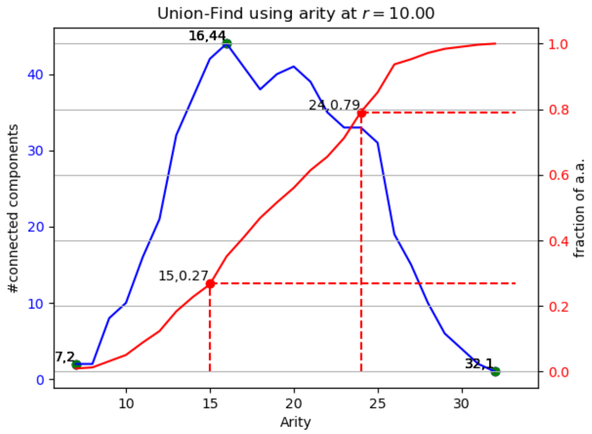

(C) Union-Find using arity at $r = 10.00$

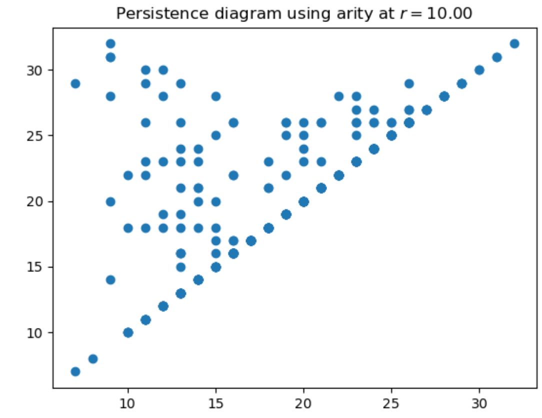

(D) Persistence diagram using arity at $r = 10.00$

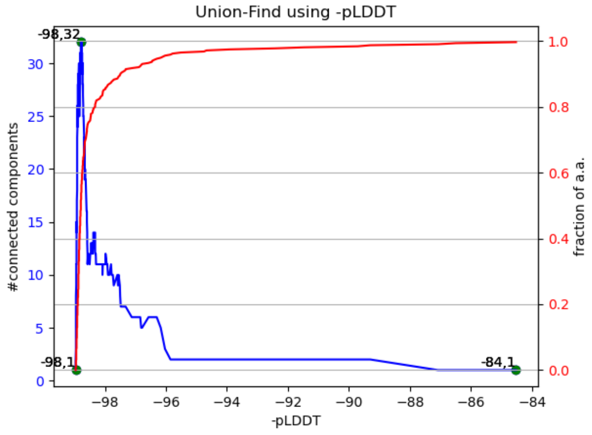

(E) Union-Find using -pLDDT

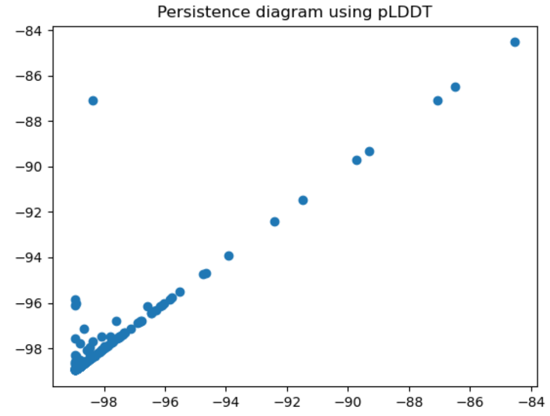

(F) Persistence diagram using pLDDT

Figure S3: **human-AF-P15121-F1-model_v4; 316 amino acids. Quantile-arity signature [(15, 0.25, 0.27), (24, 0.75, 0.79)]**

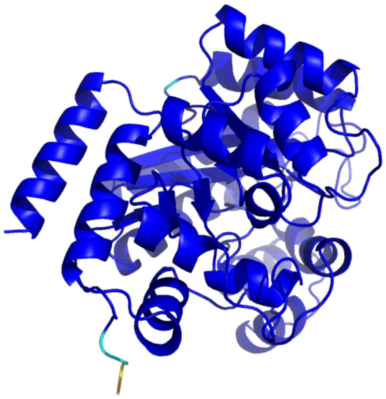

(A) 3D view

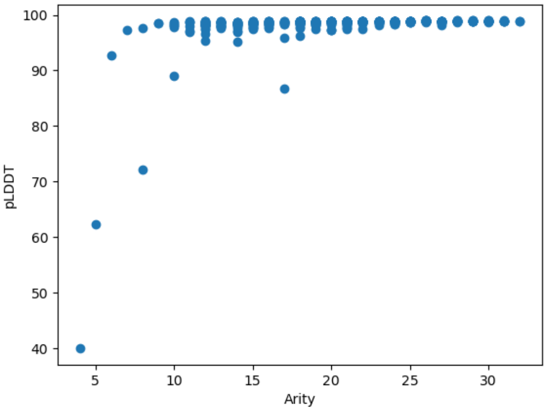

(B) Arity at $r = 10$ x pLDDT. Pearson $r = 0.31$

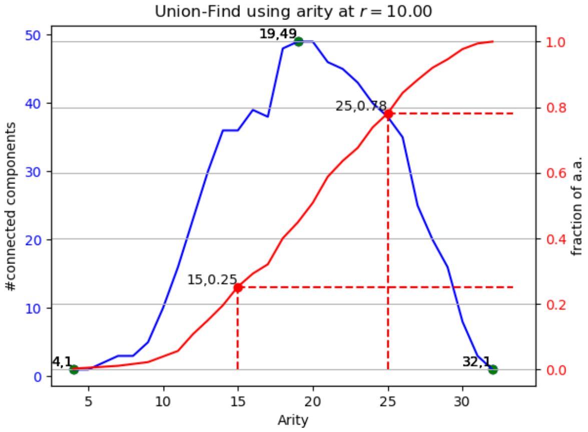

(C) Union-Find using arity at $r = 10.00$

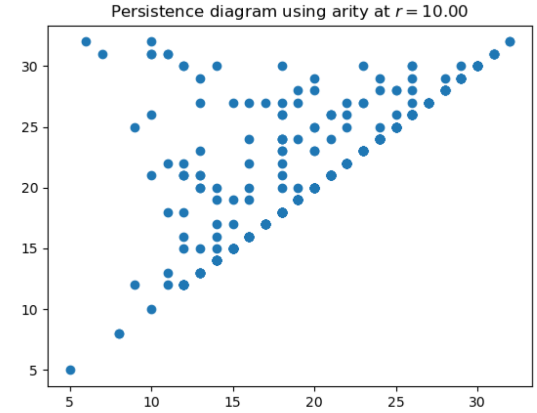

(D) Persistence diagram using arity at $r = 10.00$

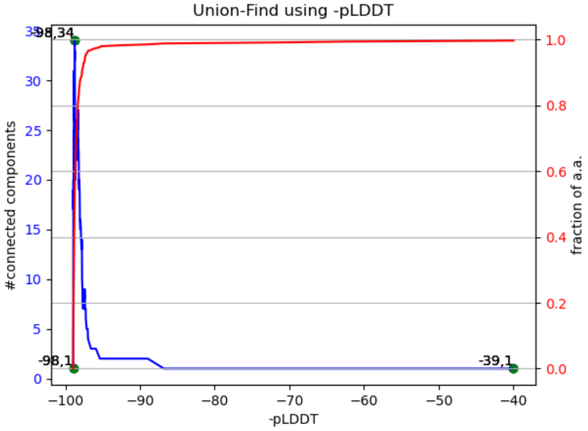

(E) Union-Find using -pLDDT

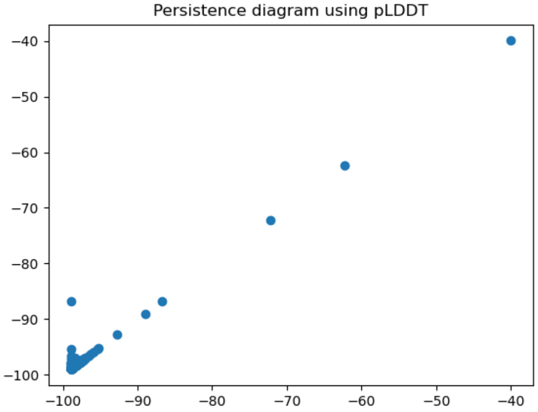

(F) Persistence diagram using pLDDT

Figure S4: **rat-AF-Q920P6-F1-model_v4; 352 amino acids. Quantile-arity signature [(15, 0.25, 0.25), (25, 0.75, 0.78)]**

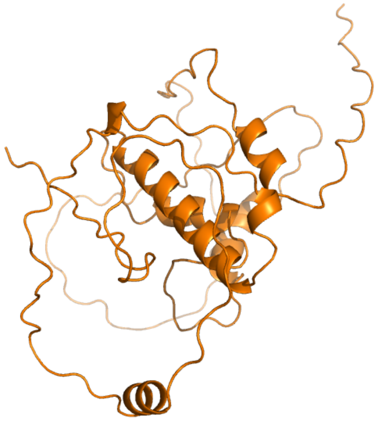

(A) 3D view

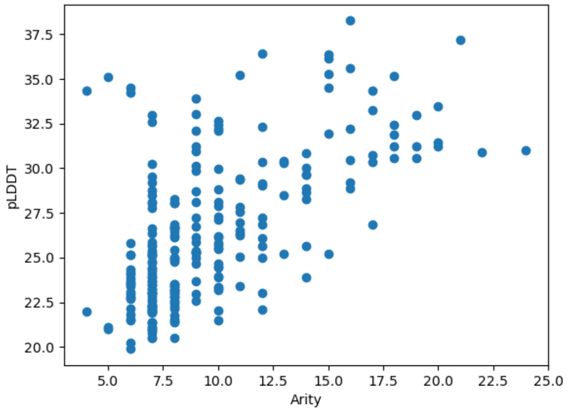

(B) Arity at $r = 10$ x pLDDT. Pearson $r = 0.61$

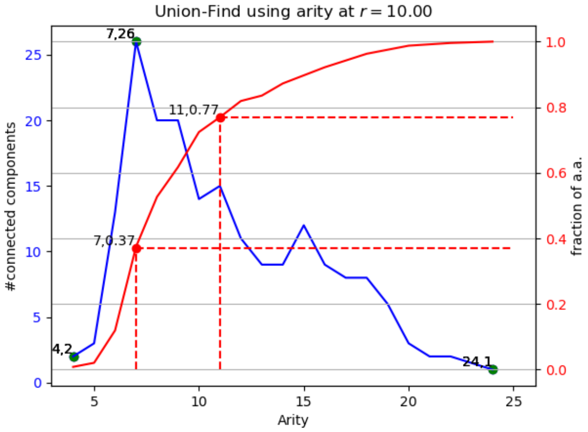

(C) Union-Find using arity at $r = 10.00$

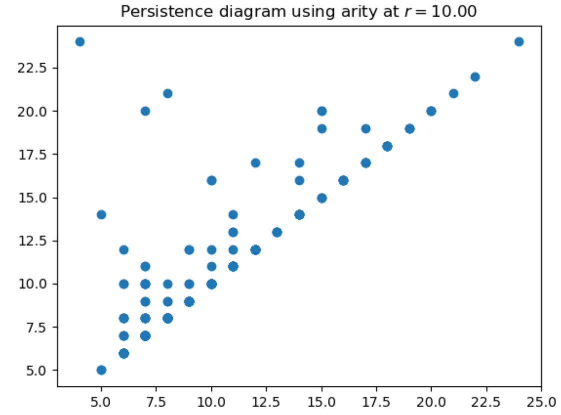

(D) Persistence diagram using arity at $r = 10.00$

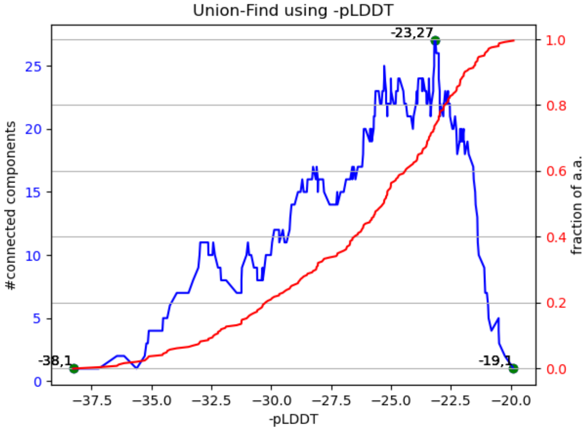

(E) Union-Find using -pLDDT

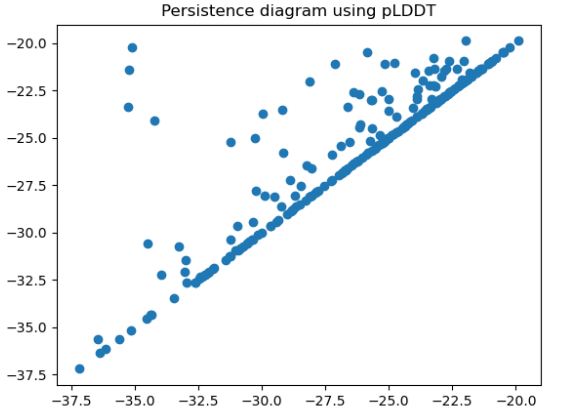

(F) Persistence diagram using pLDDT

Figure S5: **human-AF-Q96MU5-F1-model_v4; 243 amino acids. Quantile-arity signature [(7, 0.25, 0.37), (11, 0.75, 0.77)]**

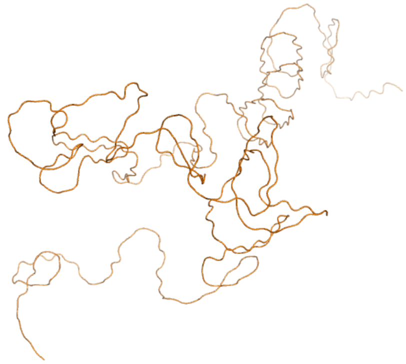

(A) 3D view

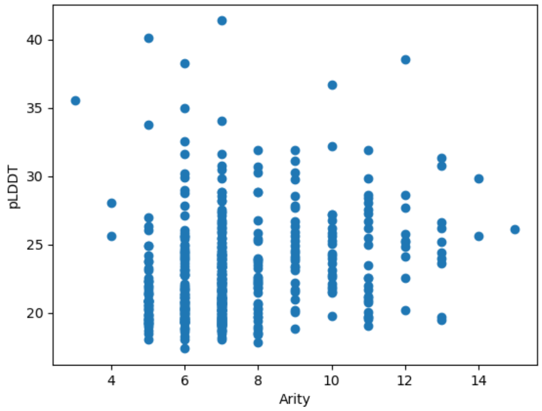

(B) Arity at $r = 10$ x pLDDT. Pearson $r = 0.23$

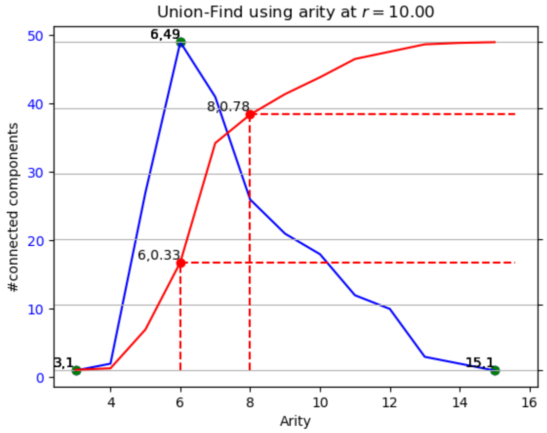

(C) Union-Find using arity at $r = 10.00$

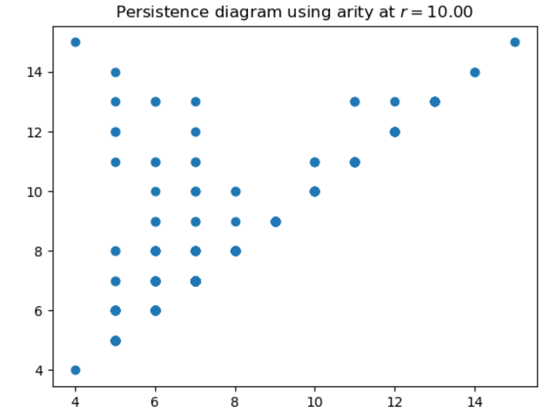

(D) Persistence diagram using arity at $r = 10.00$

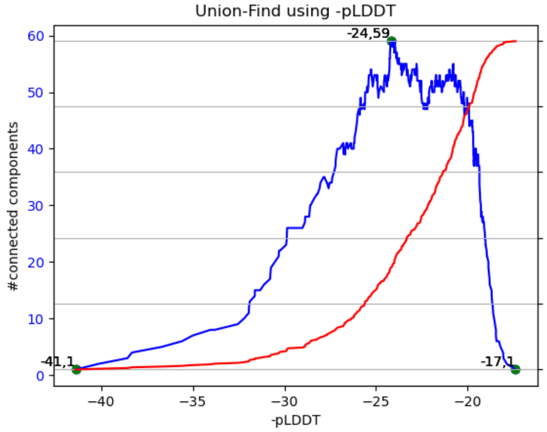

(E) Union-Find using -pLDDT

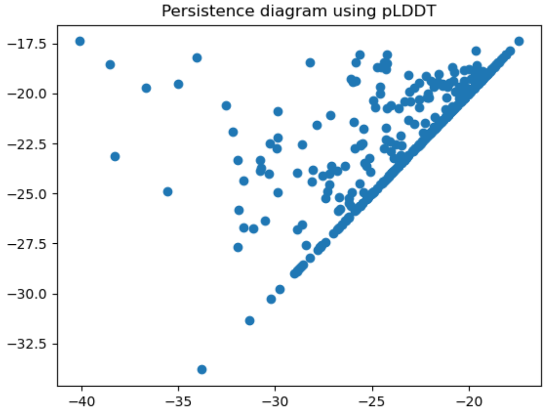

(F) Persistence diagram using pLDDT

Figure S6: **zebrafish-AF-A0A0G2L439-F1-model_v4; 449 amino acids. Quantile-arity signature [(6, 0.25, 0.33), (8, 0.75, 0.78)]**

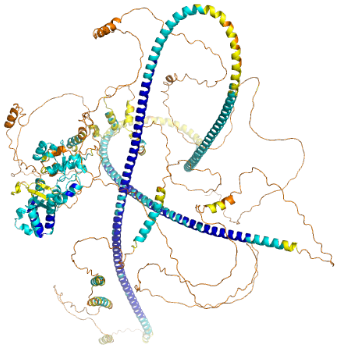

(A) 3D view

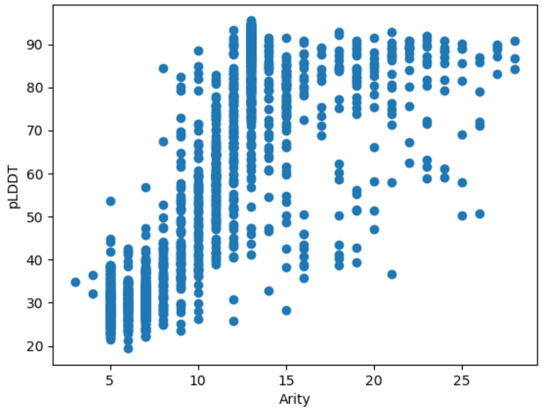

(B) Arity at $r = 10$ x pLDDT. Pearson $r = 0.75$

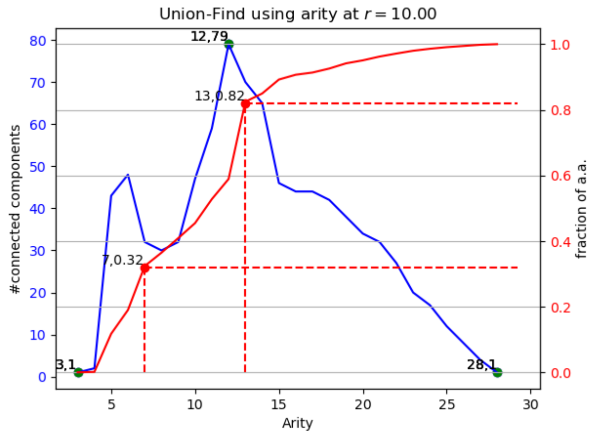

(C) Union-Find using arity at $r = 10.00$

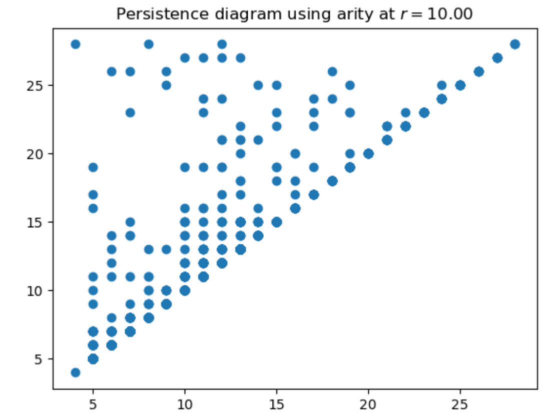

(D) Persistence diagram using arity at $r = 10.00$

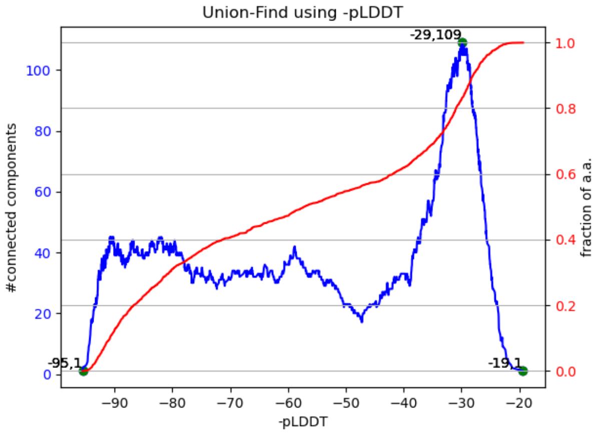

(E) Union-Find using -pLDDT

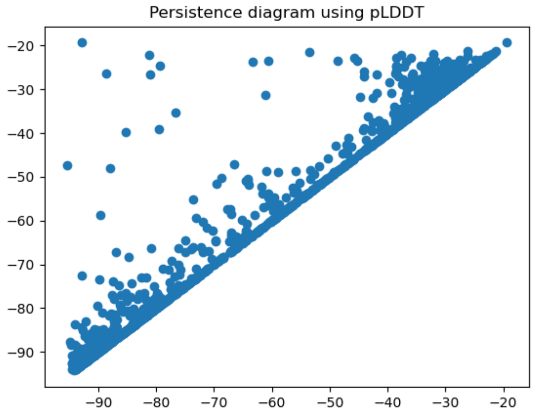

(F) Persistence diagram using pLDDT

Figure S7: **nemat-AF-A0A1C3NSL9-F1-model_v4; 1644 amino acids. Quantile-arity signature [(7, 0.25, 0.32), (13, 0.75, 0.82)]**

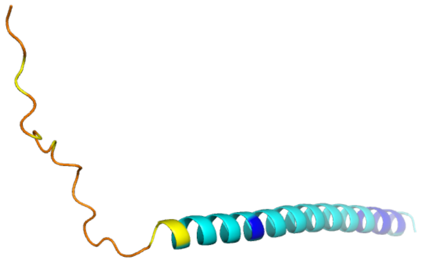

(A) 3D view

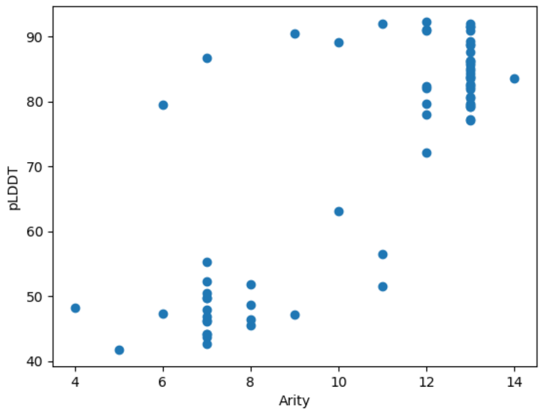

(B) Arity at $r = 10$ x pLDDT. Pearson $r = 0.83$

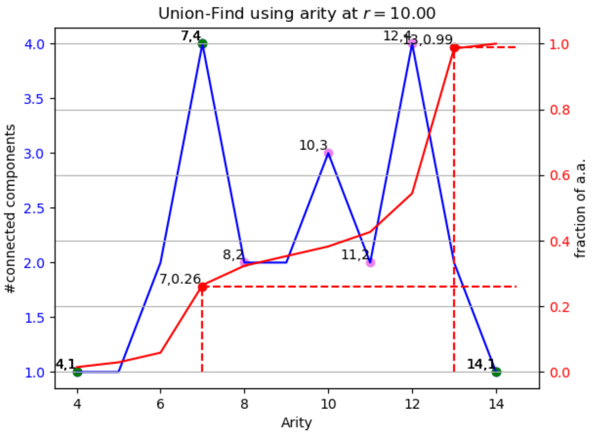

(C) Union-Find using arity at $r = 10.00$

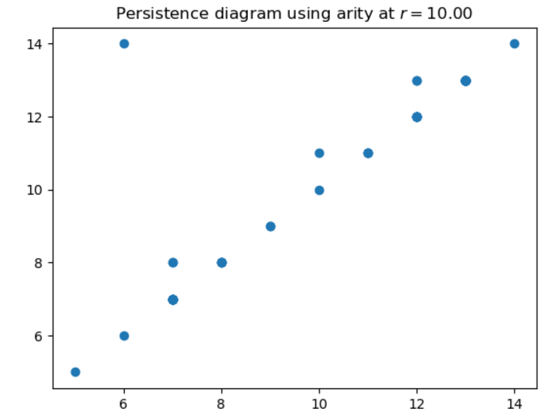

(D) Persistence diagram using arity at $r = 10.00$

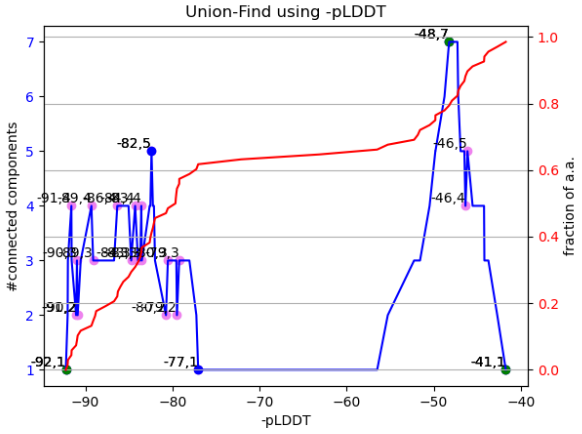

(E) Union-Find using -pLDDT

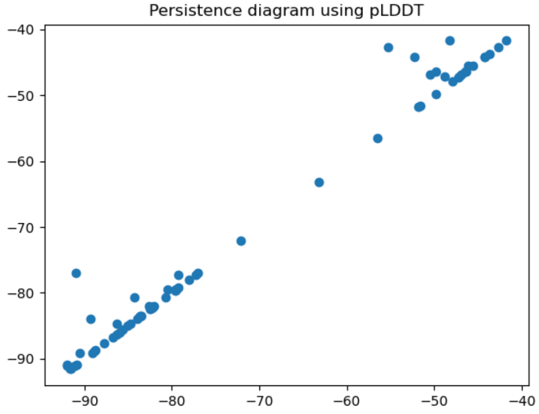

(F) Persistence diagram using pLDDT

Figure S8: **nemat-AF-Q7YTK8-F1-model_v4; 68 amino acids. Quantile-arity signature [(7, 0.25, 0.26), (13, 0.75, 0.99)]**

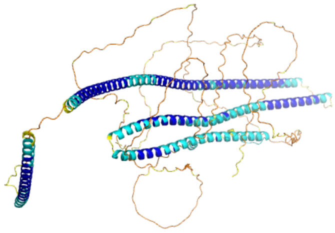

(A) 3D view

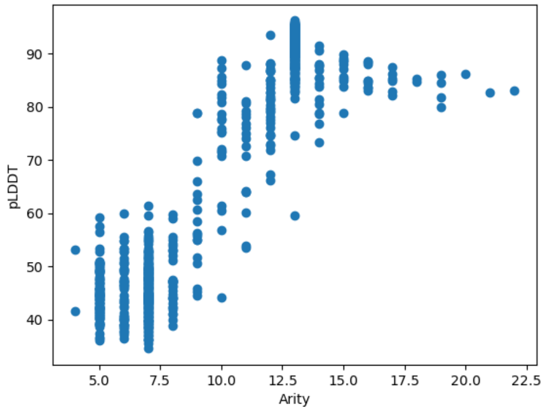

(B) Arity at $r = 10$ x pLDDT. Pearson $r = 0.91$

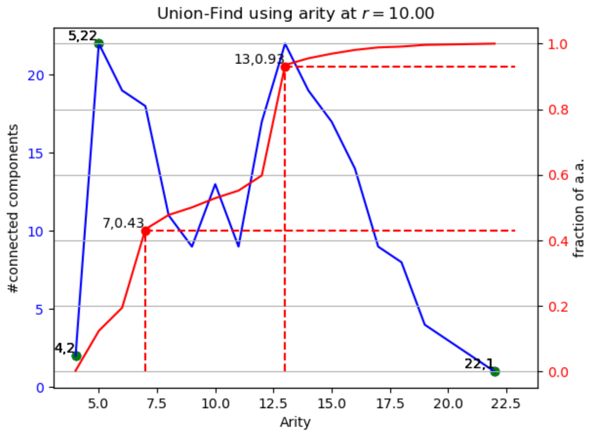

(C) Union-Find using arity at $r = 10.00$

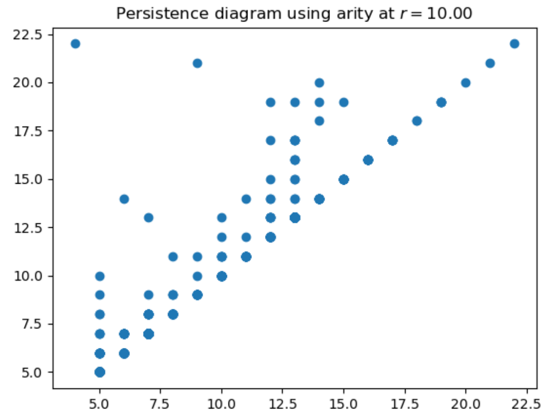

(D) Persistence diagram using arity at $r = 10.00$

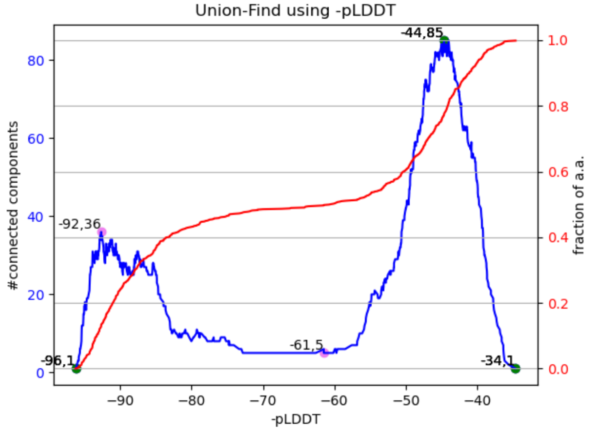

(E) Union-Find using -pLDDT

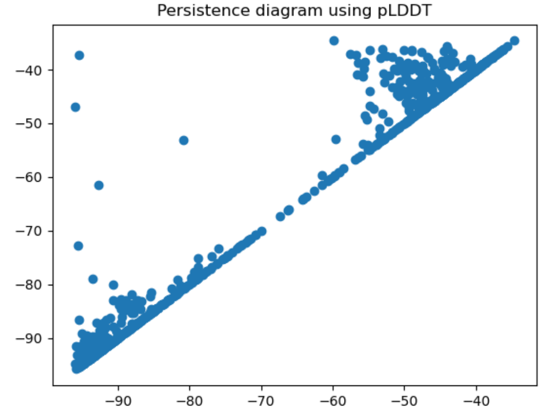

(F) Persistence diagram using pLDDT

Figure S9: **droso-AF-Q9VQS4-F1-model_v4; 781 amino acids. Quantile-arity signature [(7, 0.25, 0.43), (13, 0.75, 0.93)]**

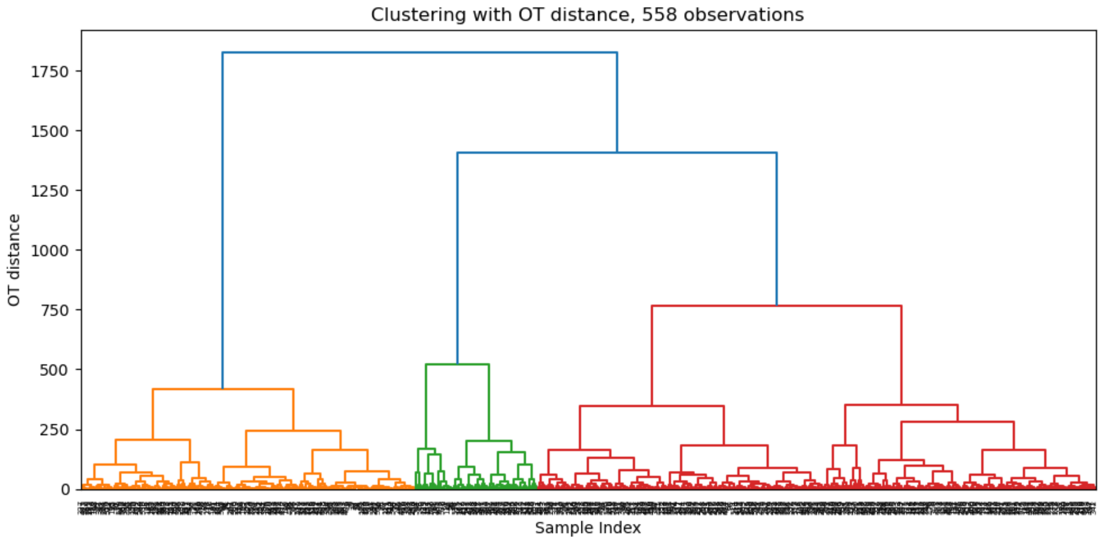

(A)

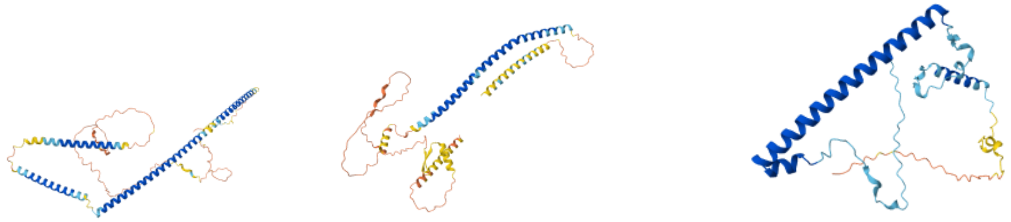

(B)

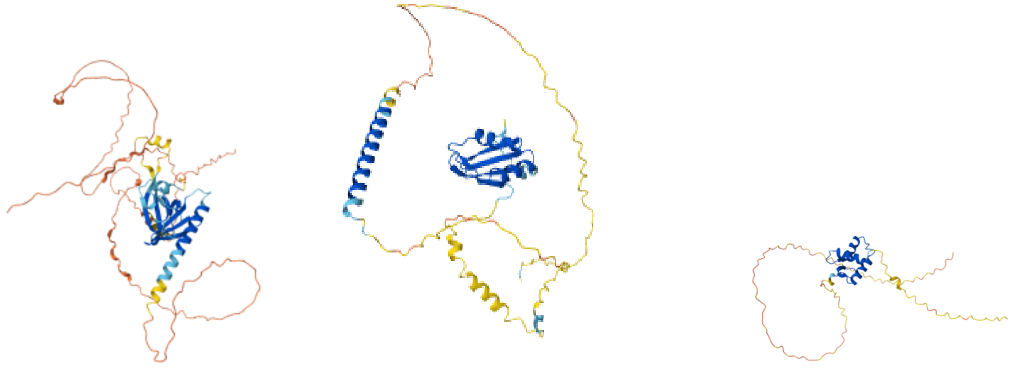

(C)

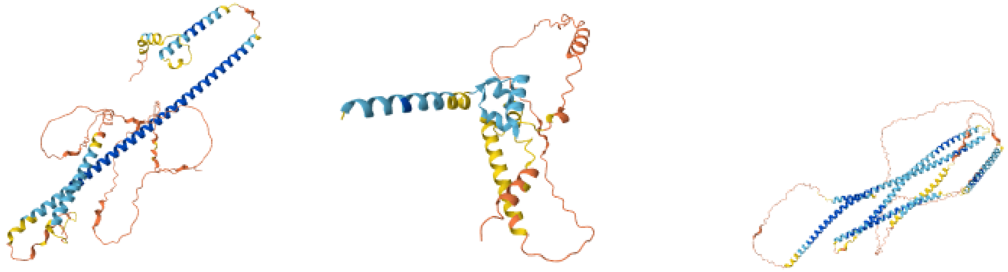

(D)

Figure S10: **H. Sapiens: hierarchical clustering of the 558 structures in the 7x13 cell of the arity map, with three random structures from every cluster.** **(A)** Ward hierarchical clustering of the structures using the optimal transportation distance on the normalized arity histograms $H_{C_\alpha}$ for each pair. **(B)** First cluster: independent helices; **(C)** Second cluster: independent helices and a small $\beta$ core; **(D)** Third cluster: packed helices.

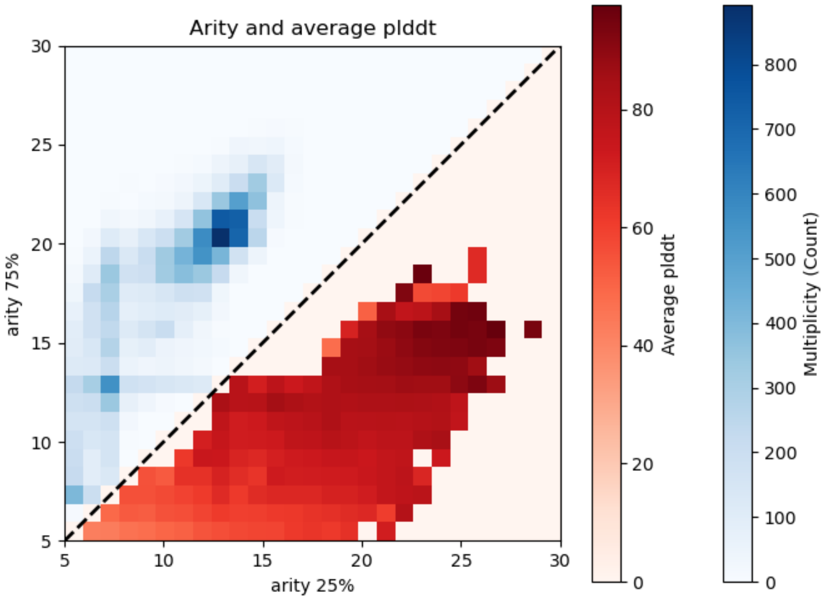

Figure S11: **pLDDT and arity in the human proteome.** The upper triangle of the heatmap shows the protein count on the arity plot. The lower triangle shows the average pLDDT per bin, with mirrored coordinates. As expected, pLDDT increases in the densely populated aread of well-folded proteins, and decreases towards the bottom-left part of the plot, where IDP/IDRs are thought to cluster

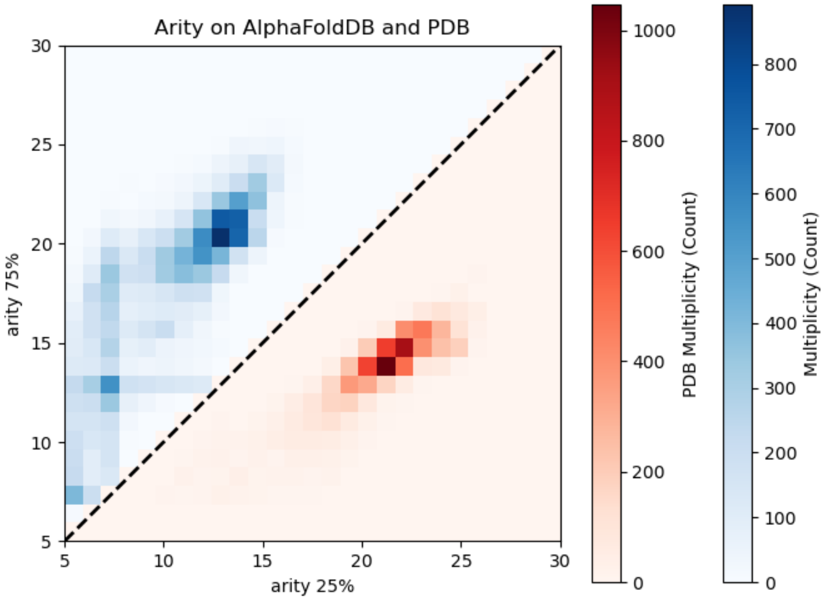

Figure S12: **Arity of AF2-predicted and experimentally determined human proteins.** The upper triangle of the heatmap shows the protein count of AF2-predicted human proteins on the arity plot. The lower triangle shows the same quantity over the experimentally determined human proteins available on the PDB. The experimental structures only group in correspondance of the main peak of the upper triangle, which is supposed to contain well-folded protein structures.

### S3 Q2. Predicted domains and their quality

**Counting homologs.** For each predicted protein of each proteome, we run a homology search over `UniProtKB` (SwissProt + TrEMBL, 2024-10-20) with `mmseqs2 easy-search`. Given the very large amount of queries, we chose `MMseqs2` over other homology search strategies (BLASTP, HMMer3) because of its time efficiency and sensitivity. Much of MMseqs rapidity comes from the prefiltering steps, which limit the number of sequence comparisons performed by the algorithm. If run with standard parameters, this strategy risks to limit the maximum number of homologs the algorithm finds per query. In order to get a reliable homolog count, we allow the maximum hits threshold parameter `-max-seqs` to attain 1,000,000. In order to further optimize the speed, we also use the parameters `-max-rejected 10` (the homolog search for a given query stops after ten consecutive rejected alignments, exploiting the sequence sorting done by the prefiltering steps) and `-min-ungapped-score 30` (restrict the rejection condition during the prefilter alignments). The E-value threshold for a match is kept to the standard `0.001`. We remind that the E-value often calculated in homology search algorithms is an estimator bound between 0 and 1 of the Karlin-Altschul statistics. For E-values $< 0.001$, it approximates the p-value to refuse the null hypothesis that a sequence with a score at least as high as the score of the hit found is not a homolog of the query sequence.

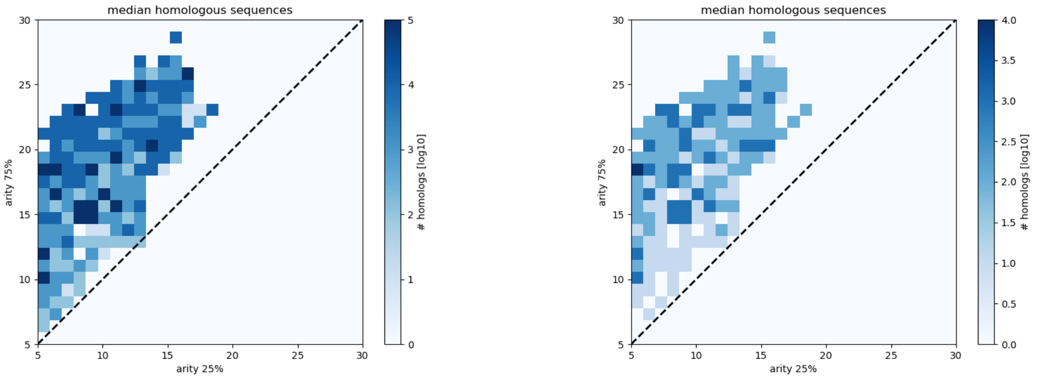

(A) (B)

Figure S13: **H. Sapiens: the number of homologous sequences correlates with arity, the number of `Uniclust` representatives does not**. **(A)** Per-structure median number of homologous `UniProtKB` sequences within each arity signature bin. Structured proteins (high-arity signatures, top right corner) usually find more homologous than partially or totally disordered proteins. **(B)** Per-structure median number of homologous `Uniclust` representative sequences within each arity signature bin. The correlation between arity and homology vanishes.

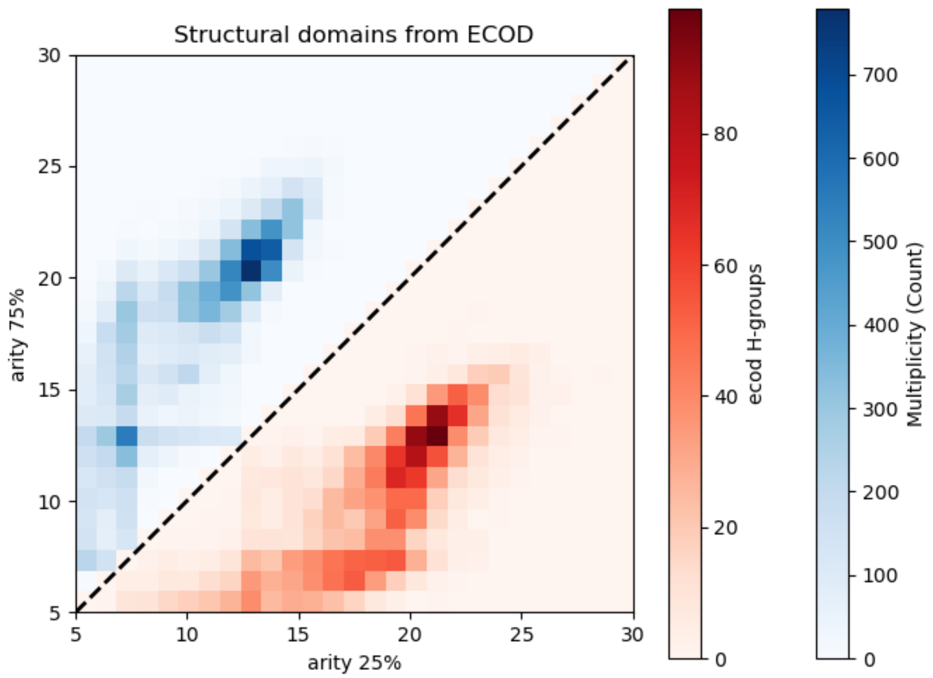

(A)

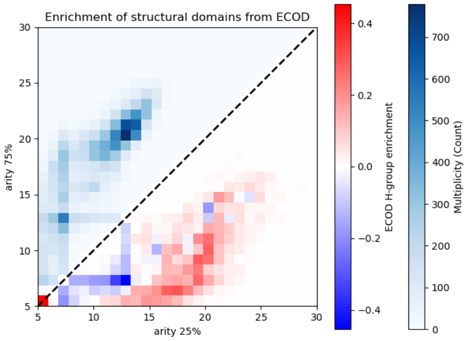

(B)

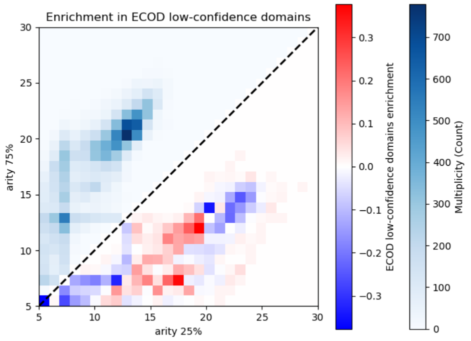

(C)

Figure S14: **H. Sapiens: arity map of the human proteome vs ECOD domain enrichment in human proteins**. The upper triangle always reports the human proteome arity map for reference. **(A)** Number of unique ECOD H-groups for each arity signature. **(B)** Enrichment difference (with respect to the proteome) of unique ECOD H-groups. **(C)** Enrichment difference (with respect to the proteome) of non-redundant ECOD low-confidence domains.

### S4 Q4. pLDDT values and fragmentation of `AlphaFold` reconstructions

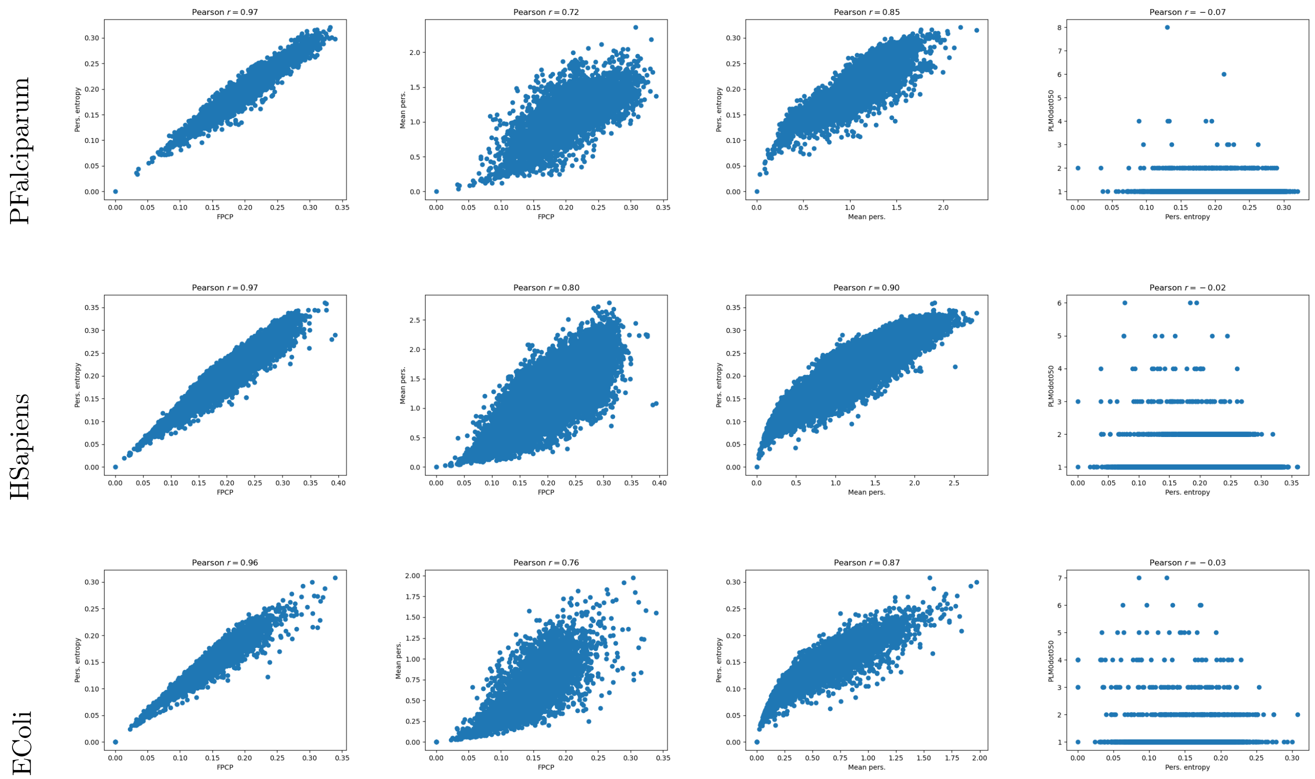

Figure S15: **PAeruginosa, HSapiens, Ecoli: correlations between the fraction of positive critical points $f^+_{cp}$, the persistence entropy $H_{\rm p}$, the mean persistence $\overline{\rm p}$, and the number of persistent local maxima PLM**$(t_{\rm p} = 0.05)$**.** Along these three examples: the fraction of positive critical points increases and so does the persistence entropy. Compare with Table 1.

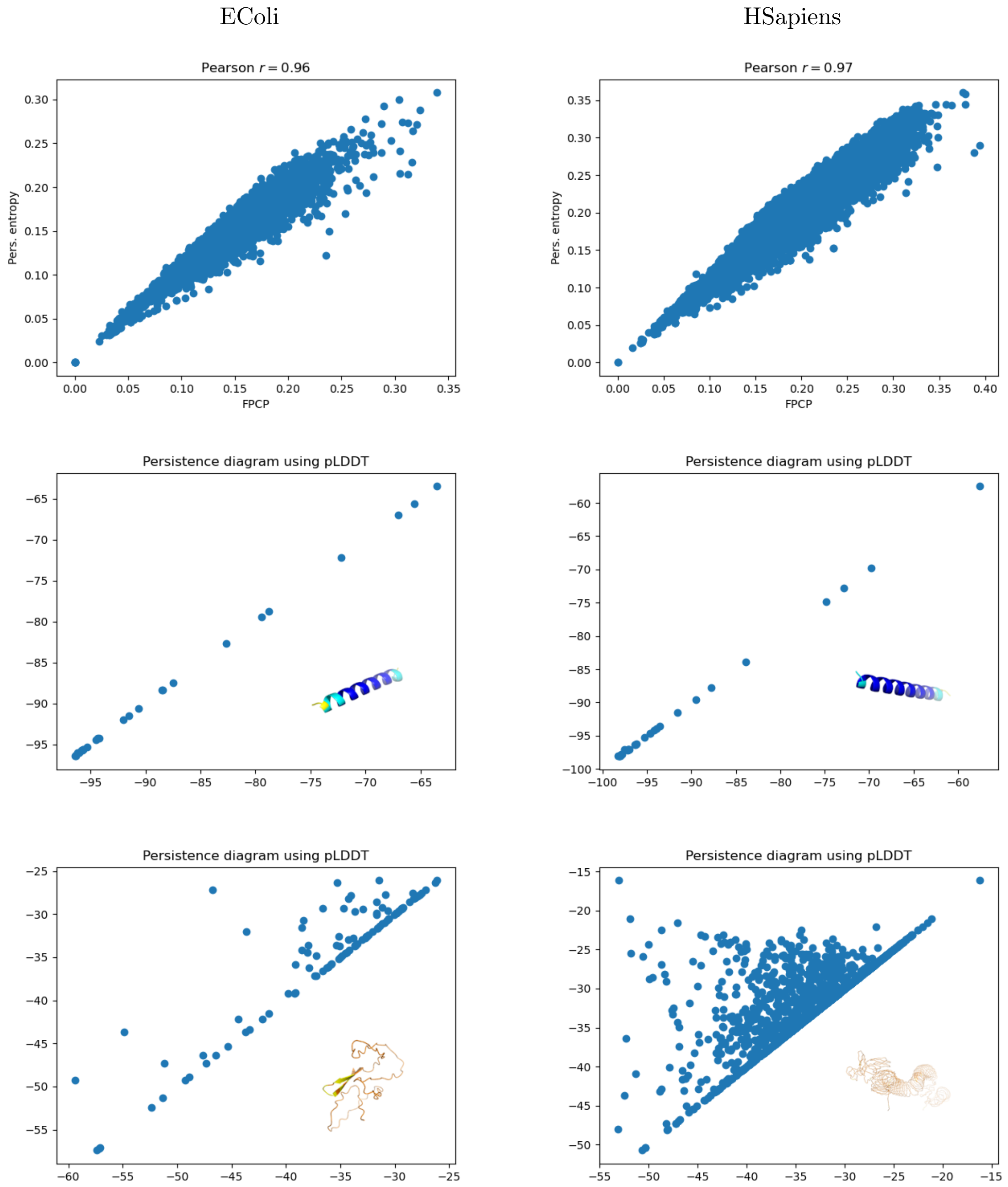

Figure S16: **Persistence entropy $H_{\mathrm{p}}$ and fraction of positive critical points (FPCP) $f_{cp}^{+}$: illustrations per genome, with structures achieving the minimum and maximum values.** Compare with Table 1. **(First row)** Scatter plot fraction of positive critical points $f_{cp}^{+} \times$ persistence entropy $H_{\mathrm{p}}$. **(Second row)** Per organism, persistence diagram and `AlphaFold` model achieving the minimum entropy $H_{\mathrm{p}}$. **(Third row)** Per organism, persistence diagram and `AlphaFold` model achieving the maximum entropy $H_{\mathrm{p}}$. **(Structures for EColi)** AF-A5A617-F1-model_v4, 27 a.a.; AF-P62066-F1-model_v4, 106 a.a. **(Structures for HSapiens)** AF-O00631-F1-model_v4, 31 a.a.; AF-Q02817-F15-model_v4, 1400 a.a.

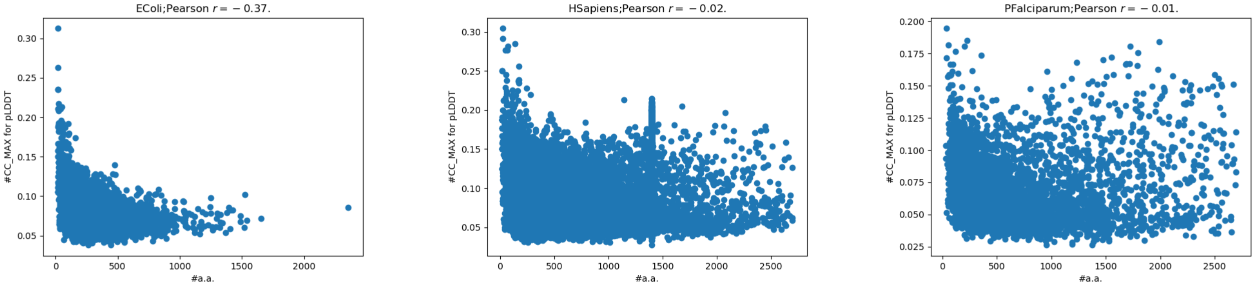

EColi HSapiens PFalciparum

Figure S17: **Relationship between the number of amino acids $n$ and the ratio $N_{\mathbf{cc}}^{max}/n$: illustration for three genomes.** We conjecture that the ratio $N_{\mathrm{cc}}^{max}/n$ for the null model is equal to 1/4.

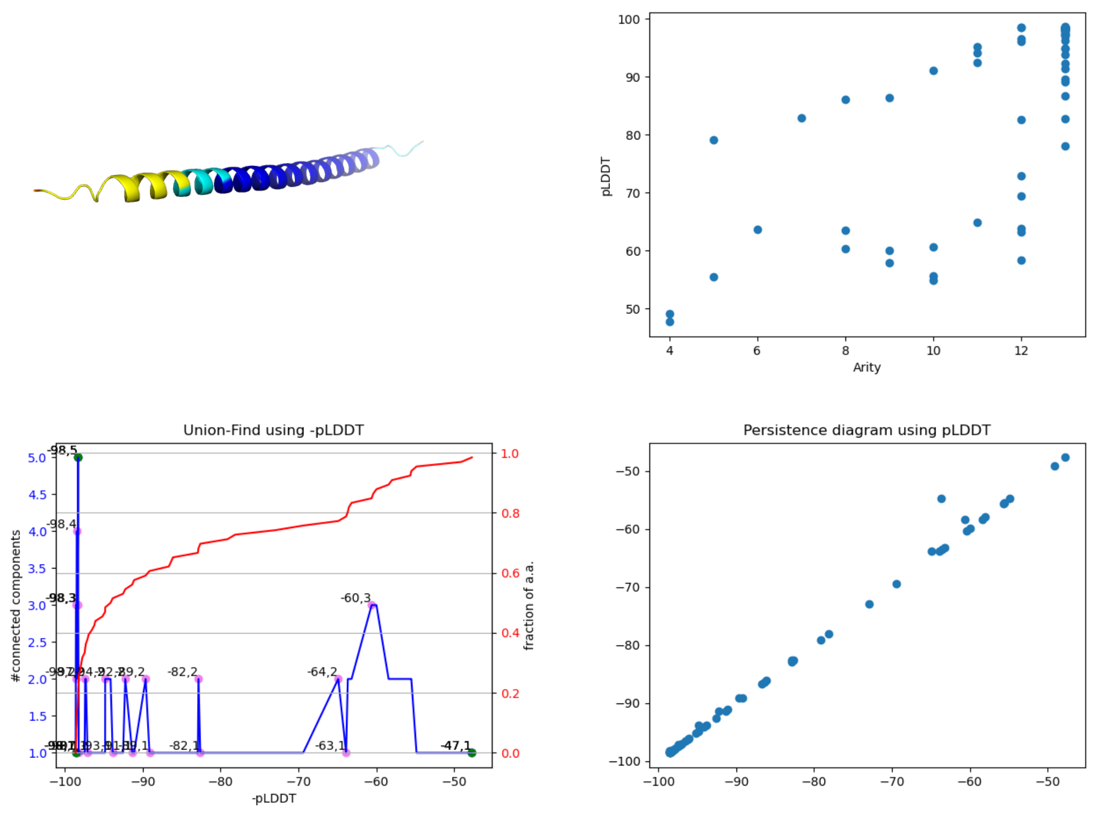

Figure S18: **Function $N_{\mathbf{cc}}$(pLDDT) and its maximum value $N_{\mathbf{cc}}^{max}$ versus persistent local maxima PLM.** Illustration with AF-Q8N300-F1-model_v4

.

### S5 Whole genomes: arity

For arity signatures, we provide two maps: in addition to the arity map discussed in the main text, we also provide a bullet plot of all pairs observed, regardless of the count for a pair $(\texttt{arity\_}C_\alpha(q_1), \texttt{arity\_}C_\alpha(q_2))$.

#### S5.1 AThaliana

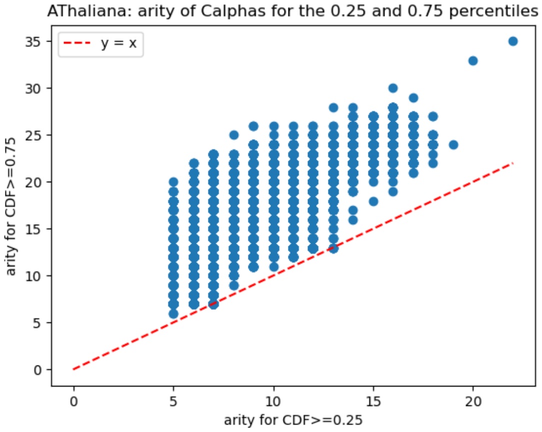

(A) Arity signatures

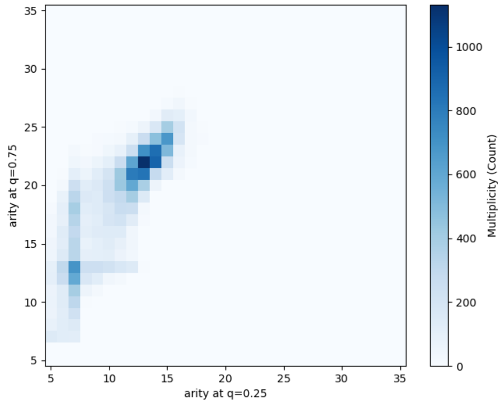

(B) Arity signatures: histogram

Figure S19: **Arity analysis for AThaliana**

#### S5.2 CAlbicans

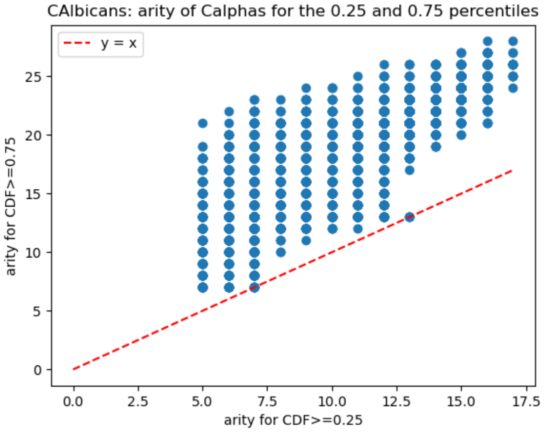

(A) Arity signatures

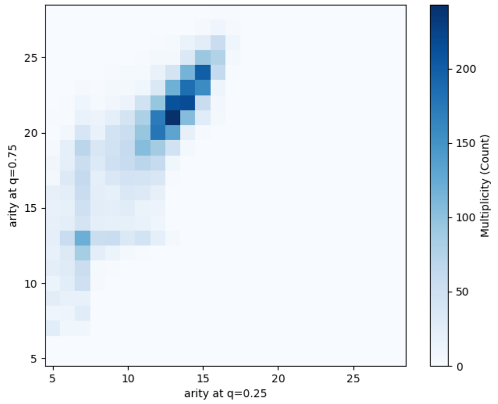

(B) Arity signatures: histogram

Figure S20: **Arity analysis for CAlbicans**

#### S5.3 CElegans

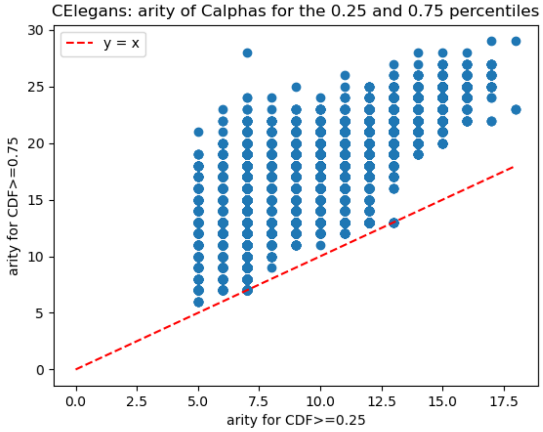

(A) Arity signatures

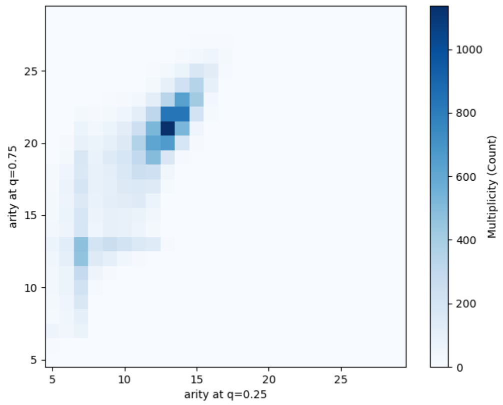

(B) Arity signatures: histogram

Figure S21: **Arity analysis for CElegans**

#### S5.4 DDiscoideum

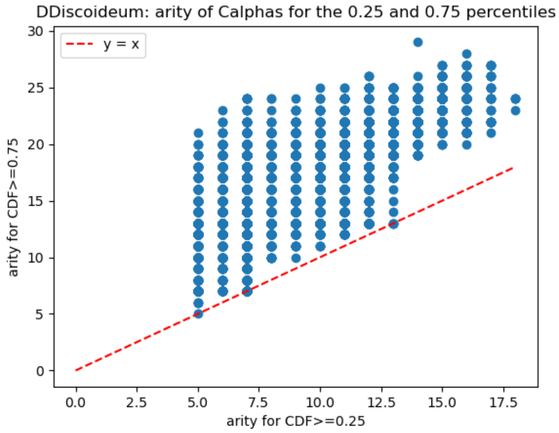

(A) Arity signatures

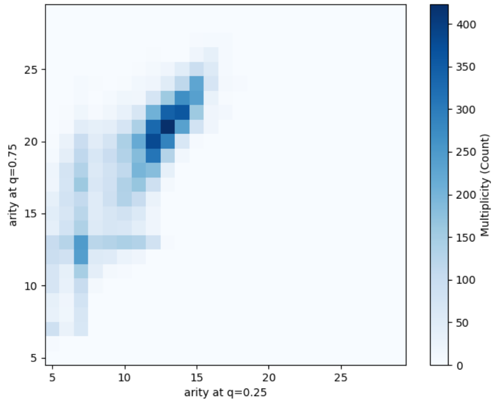

(B) Arity signatures: histogram

Figure S22: **Arity analysis for DDiscoideum**

#### S5.5 DMelanogaster

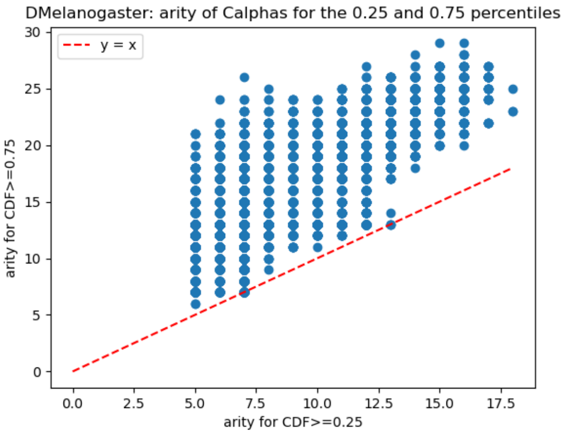

(A) Arity signatures

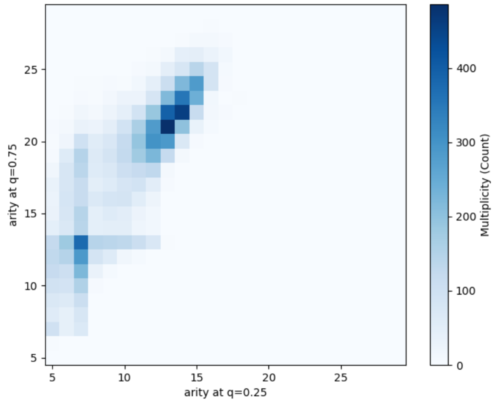

(B) Arity signatures: histogram

Figure S23: **Arity analysis for DMelanogaster**

#### S5.6 DRerio

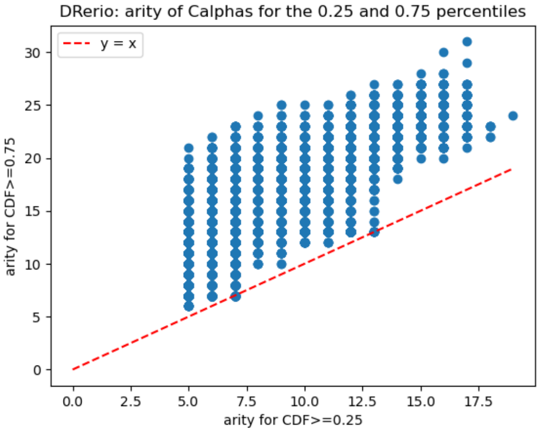

(A) Arity signatures

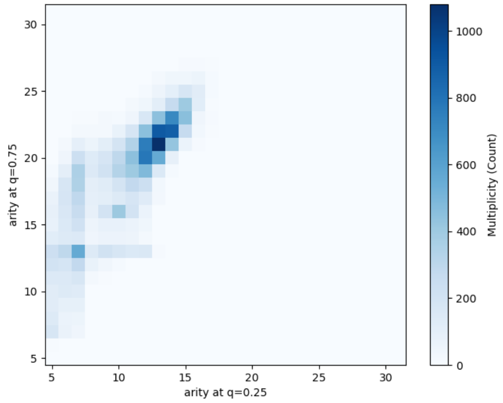

(B) Arity signatures: histogram

Figure S24: **Arity analysis for DRerio**

#### S5.7 EColi

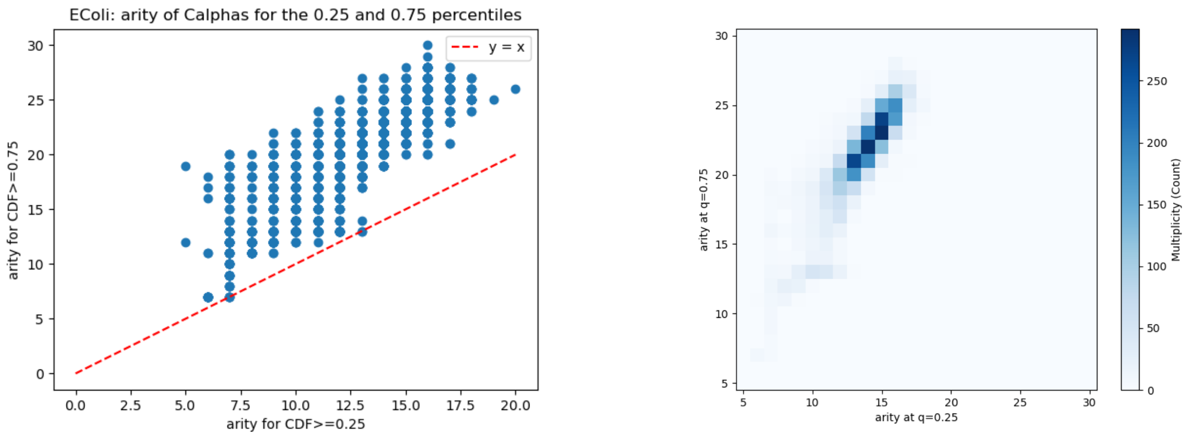

(A) Arity signatures (B) Arity signatures: histogram

Figure S25: **Arity analysis for EColi**

#### S5.8 GMax

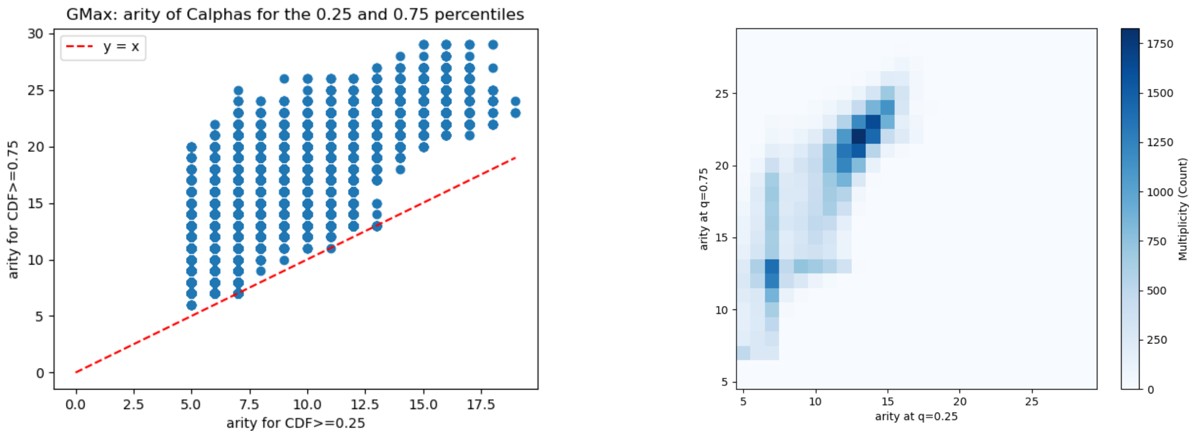

(A) Arity signatures (B) Arity signatures: histogram

Figure S26: **Arity analysis for GMax**

#### S5.9 HPylori

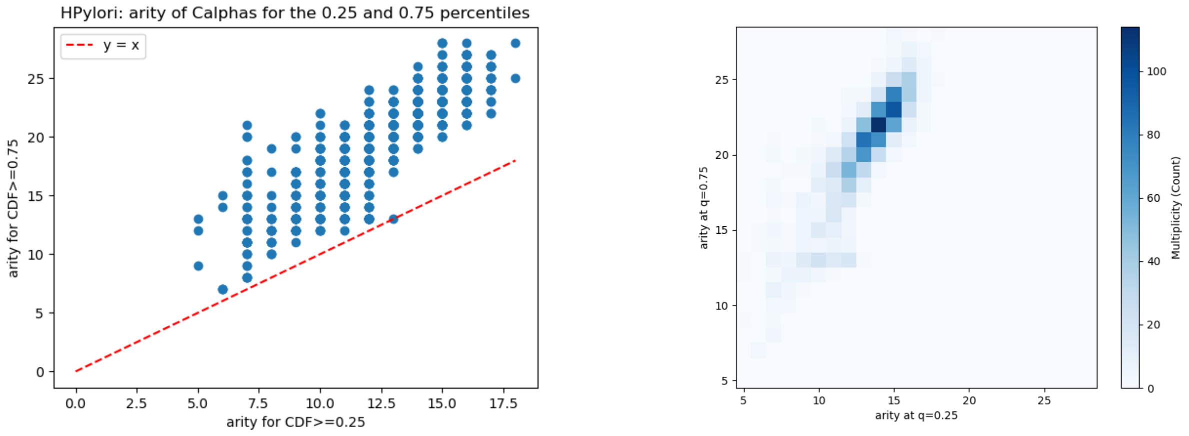

(A) Arity signatures

(B) Arity signatures: histogram

Figure S27: **Arity analysis for HPylori**

#### S5.10 HSapiens

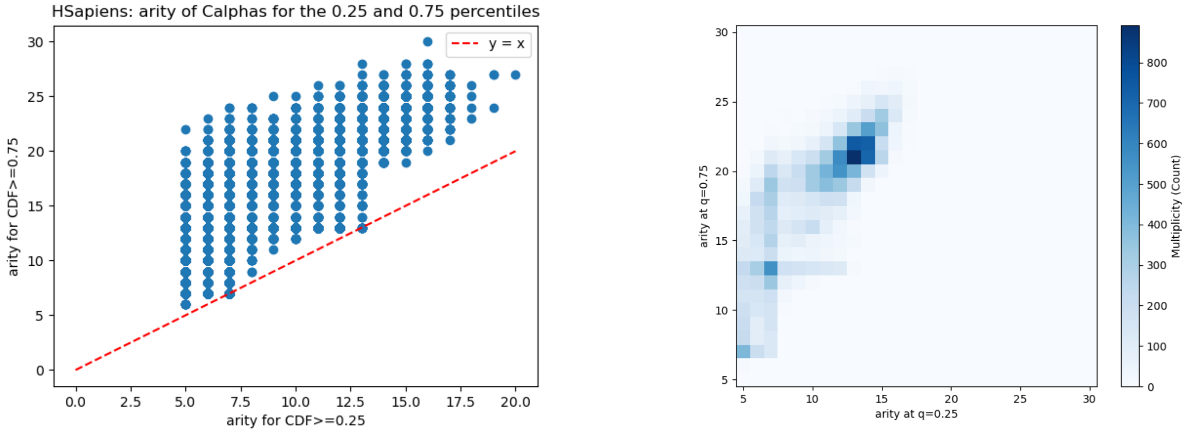

(A) Arity signatures

(B) Arity signatures: histogram

Figure S28: **Arity analysis for HSapiens**

#### S5.11 MJannaschii

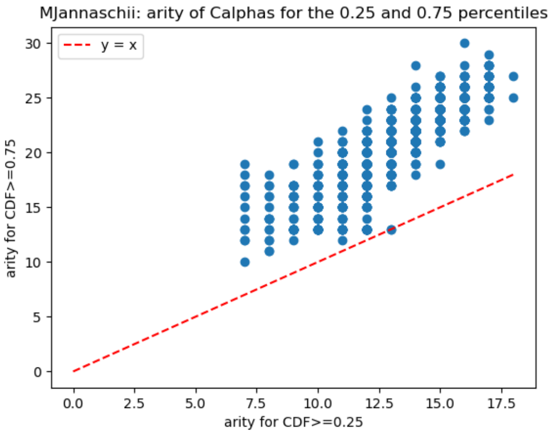

(A) Arity signatures

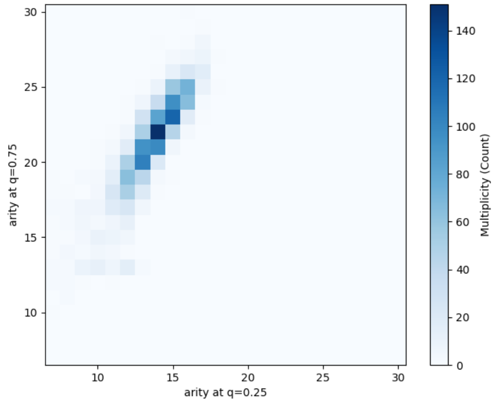

(B) Arity signatures: histogram

Figure S29: **Arity analysis for MJannaschii**

#### S5.12 MMusculus

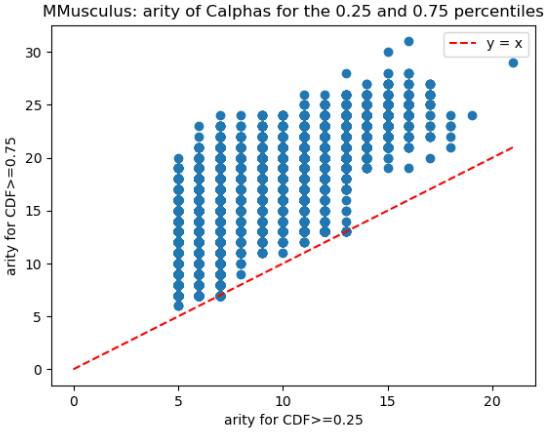

(A) Arity signatures

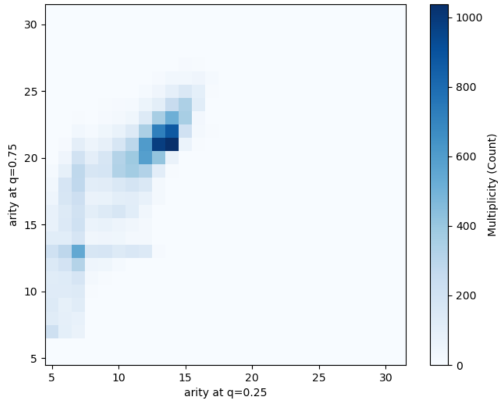

(B) Arity signatures: histogram

Figure S30: **Arity analysis for MMusculus**

#### S5.13 MTuberculosis

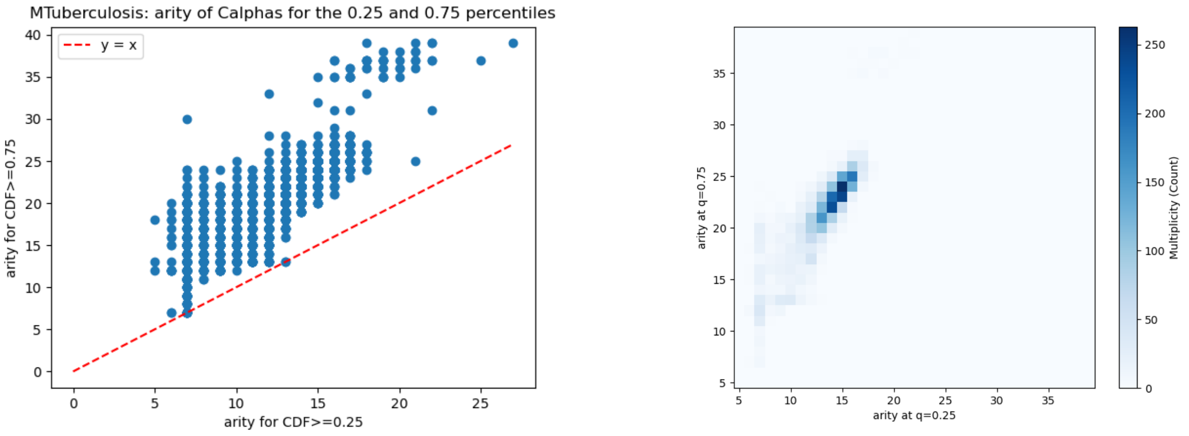

(A) Arity signatures (B) Arity signatures: histogram

Figure S31: **Arity analysis for MTuberculosis**

#### S5.14 OryzaSativa

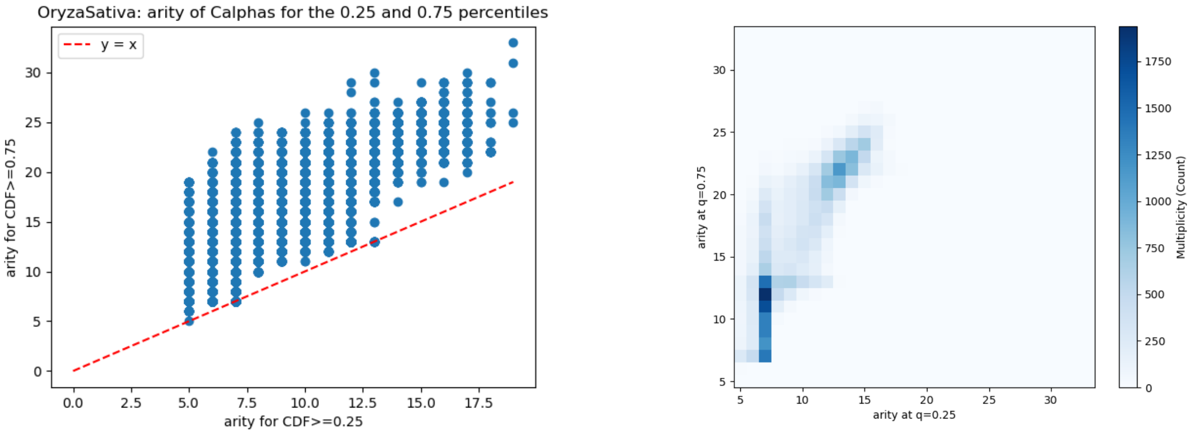

(A) Arity signatures (B) Arity signatures: histogram

Figure S32: **Arity analysis for OryzaSativa**

#### S5.15 PAeruginosa

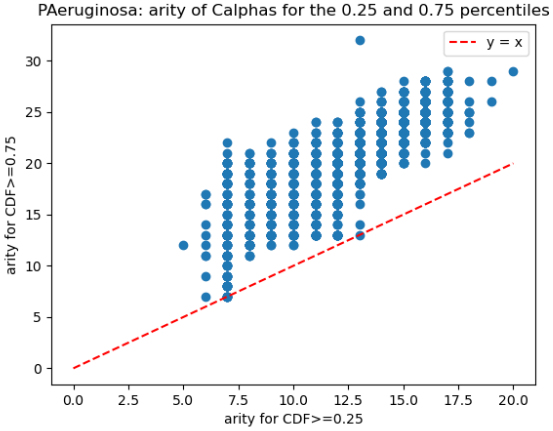

(A) Arity signatures

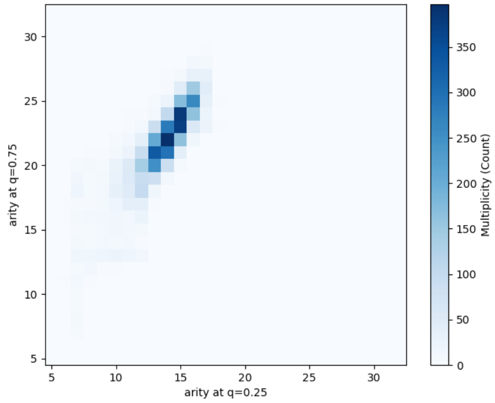

(B) Arity signatures: histogram

Figure S33: **Arity analysis for PAeruginosa**

#### S5.16 PFalciparum

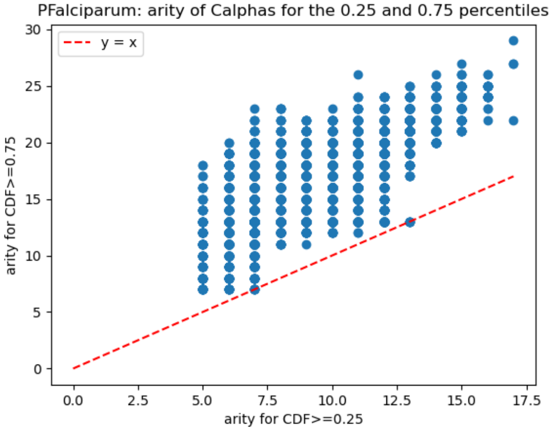

(A) Arity signatures

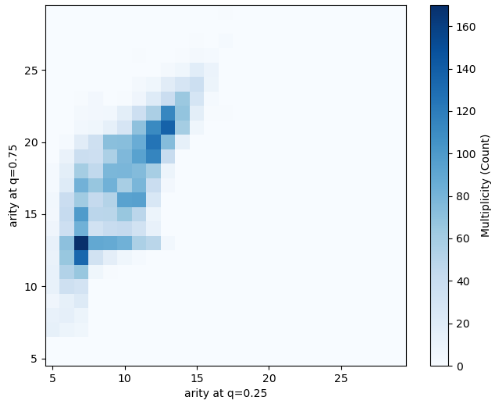

(B) Arity signatures: histogram

Figure S34: **Arity analysis for PFalciparum**

#### S5.17 RattusNorvegicus

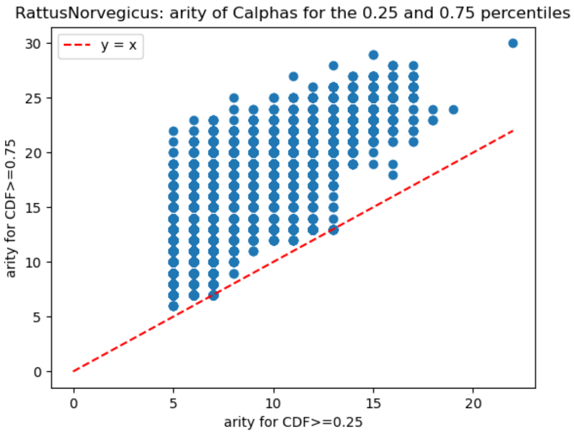

(A) Arity signatures

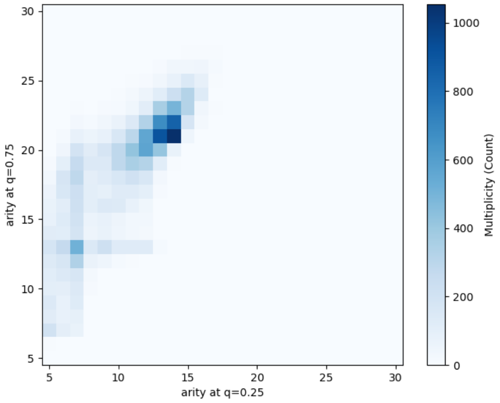

(B) Arity signatures: histogram

Figure S35: **Arity analysis for RattusNorvegicus**

#### S5.18 SAureus

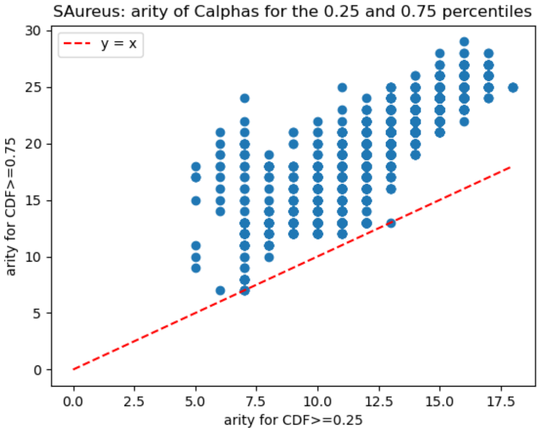

(A) Arity signatures

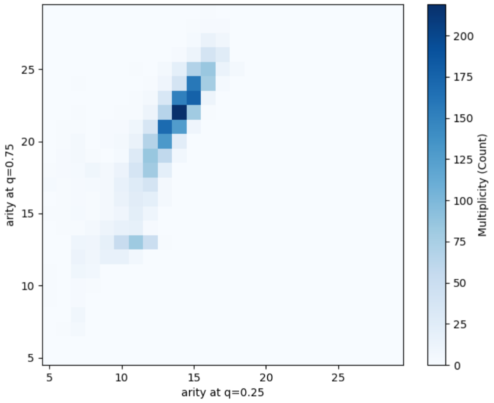

(B) Arity signatures: histogram

Figure S36: **Arity analysis for SAureus**

#### S5.19 SCerevisiae

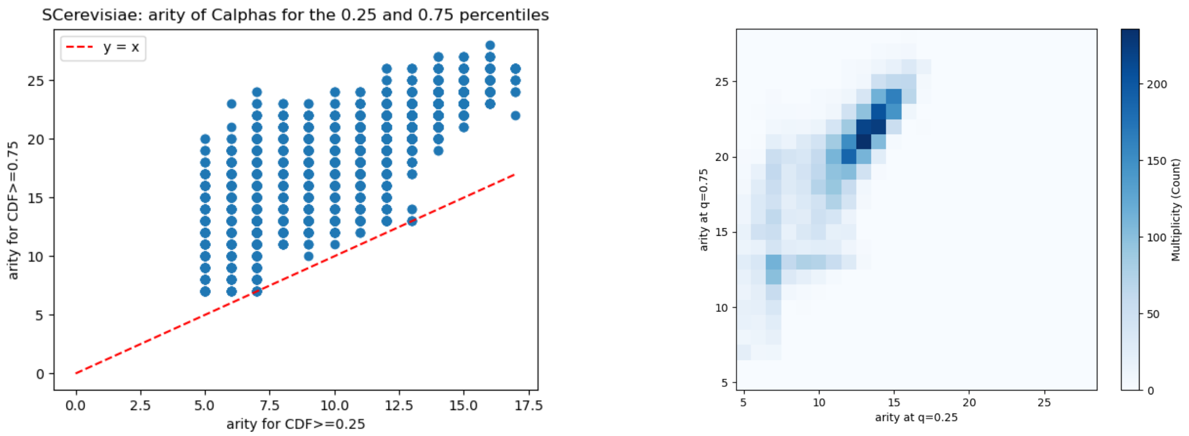

(A) Arity signatures (B) Arity signatures: histogram

Figure S37: **Arity analysis for SCerevisiae**

#### S5.20 SPombe

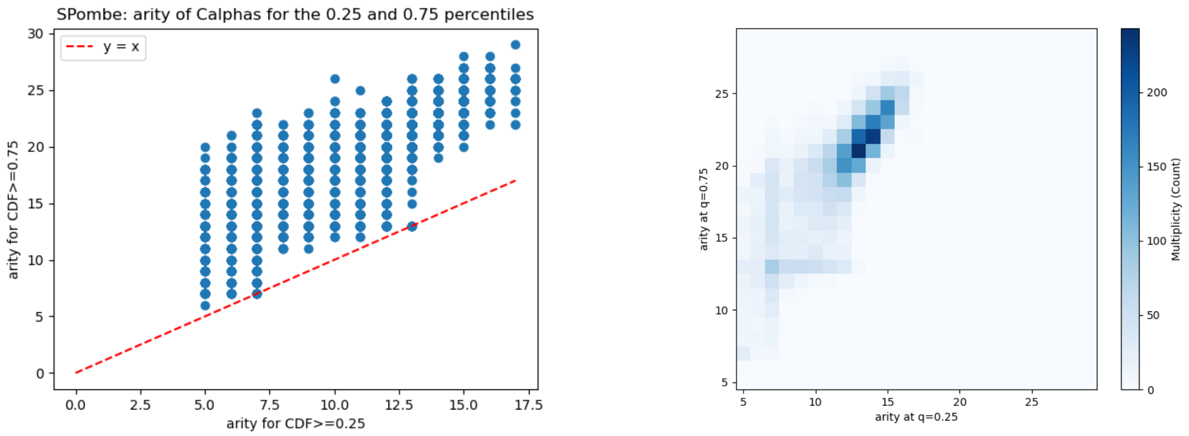

(A) Arity signatures (B) Arity signatures: histogram

Figure S38: **Arity analysis for SPombe**

#### S5.21 ZeaMays

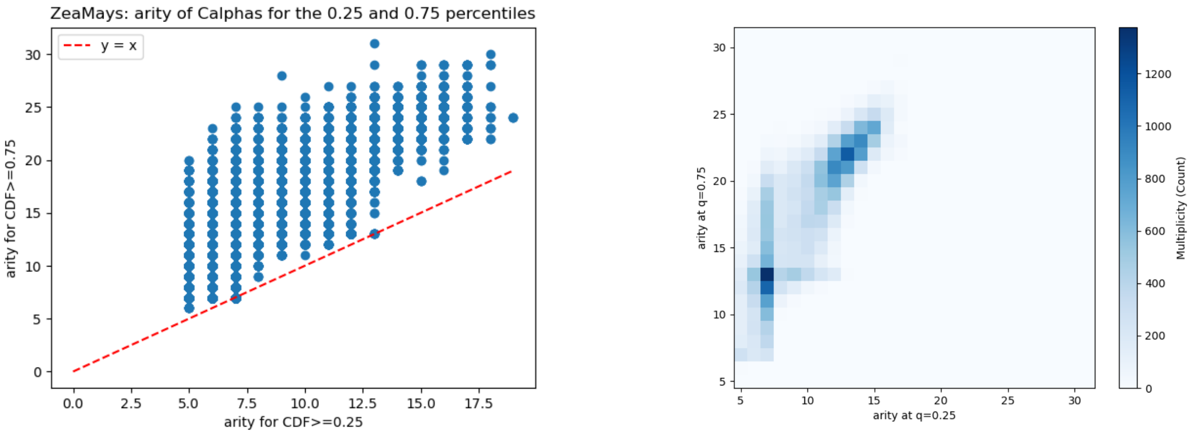

(A) Arity signatures

(B) Arity signatures: histogram

Figure S39: **Arity analysis for ZeaMays**

### S6 Whole genomes: pLDDT

#### S6.1 pLDDT statistics per organism

For the reference, this section provides simple analysis on pLDDT values.

Let us recall the following intervals for pLDDT values [1]: very high: $0.9 \leq$ pLDDT; high: $0.7 \leq$ pLDDT $< 0.9$; low: $0.5 \leq$ pLDDT $< 0.7$; very low pLDDT $< 0.5$.

While the distribution of pLDDT values has been studied in selected cases, in particular for H. Sapiens in the context of conditionally folding proteins [26], we perform instead a systematic study on all genomes. We process all predictions (PDB files from `AlphaFold-DB`) of a given organism, collecting, per prediction, all/the median/the mean pLDDT value(s). These values yield three distributions, namely that for (i) all pLDDT values of all a.a. of all proteins, (ii) all median pLDDT values, and (iii) and all mean pLDDT values (Table 1 and Section S6). For the sake of conciseness, we solely discuss the case of all values.

Per genome statistics show that the fraction of a.a. with pLDDT below 0.7 lies in the range 0.078-0.408 and is rather consistent for model organisms (Table 1). However, the inspection of other organisms shows major discrepancies, with a value as low as 0.086 for P. Aeruginosa and as high as 0.584 for P. Falciparum (Table 1)

#### S6.2 pLDDT statistics per organism

For the reference, this section provides simple analysis on pLDDT values.

Let us recall the following intervals for pLDDT values [1]: very high: $0.9 \leq$ pLDDT; high: $0.7 \leq$ pLDDT $< 0.9$; low: $0.5 \leq$ pLDDT $< 0.7$; very low pLDDT $< 0.5$.

While the distribution of pLDDT values has been studied in selected cases, in particular for H. Sapiens in the context of conditionally folding proteins [26], we perform instead a systematic study on all genomes. We process all predictions (PDB files from `AlphaFold-DB`) of a given organism, collecting, per prediction, all/the median/the mean pLDDT value(s). These values yield three distributions, namely that for (i) all pLDDT values of all a.a. of all proteins, (ii) all median pLDDT values, and (iii) and all mean pLDDT values (Table 1 and Section S6). For the sake of conciseness, we solely discuss the case of all values.

Per genome statistics show that the fraction of a.a. with pLDDT below 0.7 lies in the range 0.078-0.408 and is rather consistent for model organisms (Table 1). However, the inspection of other organisms shows major discrepancies, with a value as low as 0.086 for P. Aeruginosa and as high as 0.584 for P. Falciparum (Table 1)

#### S6.3 AThaliana

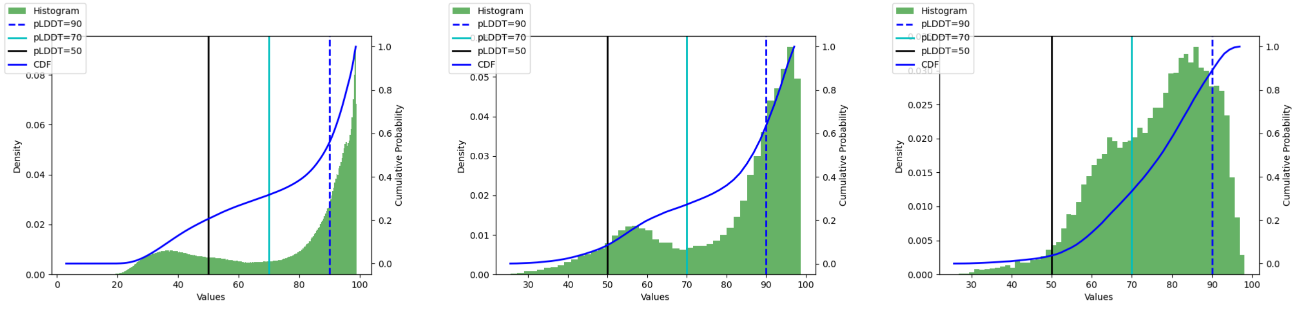

pLDDT all    pLDDT median    pLDDT mean

Figure S40: **AThaliana: pLDDT statistics**

| | 50 | 70 | 90 |
|---|---|---|---|
| AThaliana/all | 0.204 | 0.315 | 0.550 |
| AThaliana/median | 0.071 | 0.261 | 0.564 |
| AThaliana/mean | 0.032 | 0.308 | 0.853 |

Table S1: **AThaliana: CDF pLDDT**

- Worst cases, top 3: ['File AThaliana/AF-F4J1R8-F1-model_v4.pdb has 167 a.a.; pLDDT min/med/-max: 20.21 25.47 44.17', 'File AThaliana/AF-F4HT07-F1-model_v4.pdb has 221 a.a.; pLDDT min/med/-max: 18.70 25.71 31.90', 'File AThaliana/AF-Q9LPG4-F1-model_v4.pdb has 193 a.a.; pLDDT min/med/-max: 19.49 25.81 42.16']
- Median cases, 3 of them: ['File AThaliana/AF-O49680-F1-model_v4.pdb has 951 a.a.; pLDDT min/med/-max: 20.81 88.23 96.79', 'File AThaliana/AF-O64774-F1-model_v4.pdb has 749 a.a.; pLDDT min/med/-max: 23.49 88.23 98.20', 'File AThaliana/AF-O82492-F1-model_v4.pdb has 247 a.a.; pLDDT min/med/-max: 36.31 88.23 96.91']
- Top cases, top 3: ['File AThaliana/AF-Q8RY79-F1-model_v4.pdb has 490 a.a.; pLDDT min/med/-max: 29.56 98.75 98.96', 'File AThaliana/AF-Q9M8R4-F1-model_v4.pdb has 388 a.a.; pLDDT min/med/-max: 41.37 98.77 98.97', 'File AThaliana/AF-Q42578-F1-model_v4.pdb has 335 a.a.; pLDDT min/med/-max: 40.55 98.79 98.96']

#### S6.4 CAlbicans

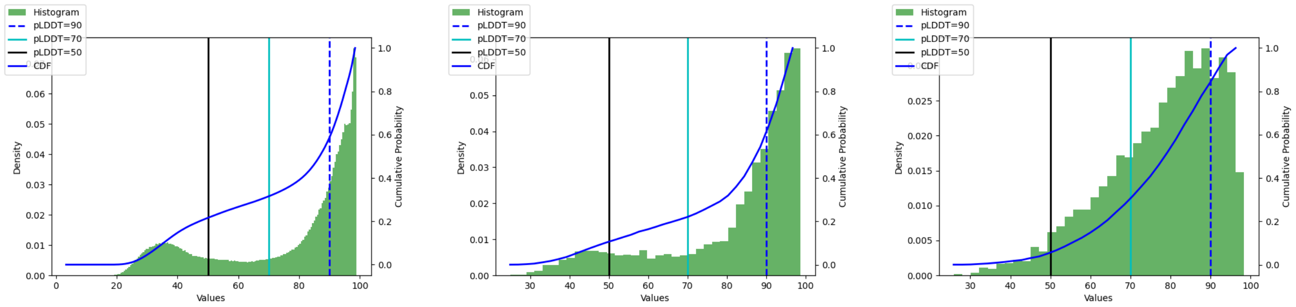

pLDDT all | pLDDT median | pLDDT mean

Figure S41: **CAlbicans: pLDDT statistics**

| | 50 | 70 | 90 |
|---|---|---|---|
| CAlbicans/all | 0.213 | 0.312 | 0.570 |
| CAlbicans/median | 0.094 | 0.209 | 0.525 |
| CAlbicans/mean | 0.042 | 0.267 | 0.783 |

Table S2: **CAlbicans: CDF pLDDT**

- Worst cases, top 3: ['File CAlbicans/AF-A0A1D8PFR1-F1-model_v4.pdb has 409 a.a.; pLDDT min/med/-max: 19.35 24.79 43.75', 'File CAlbicans/AF-Q5A3K4-F1-model_v4.pdb has 459 a.a.; pLDDT min/med/-

max: 19.01 25.39 46.51', 'File CAlbicans/AF-Q5AJV5-F1-model_v4.pdb has 1114 a.a.; pLDDT min/med/-max: 19.17 28.36 75.12']

- Median cases, 3 of them: ['File CAlbicans/AF-A0A1D8PKQ8-F1-model_v4.pdb has 472 a.a.; pLDDT min/med/max: 23.97 89.36 98.84', 'File CAlbicans/AF-Q5A283-F1-model_v4.pdb has 444 a.a.; pLDDT min/med/max: 41.58 89.36 96.83', 'File CAlbicans/AF-Q5AI67-F1-model_v4.pdb has 125 a.a.; pLDDT min/med/max: 52.88 89.36 96.34']
- Top cases, top 3: ['File CAlbicans/AF-A0A1D8PLJ3-F1-model_v4.pdb has 154 a.a.; pLDDT min/med/-max: 89.07 98.70 98.94', 'File CAlbicans/AF-Q5A750-F1-model_v4.pdb has 677 a.a.; pLDDT min/med/-max: 67.37 98.73 98.98', 'File CAlbicans/AF-P29717-F1-model_v4.pdb has 438 a.a.; pLDDT min/med/-max: 28.35 98.77 98.96']

#### S6.5 CElegans

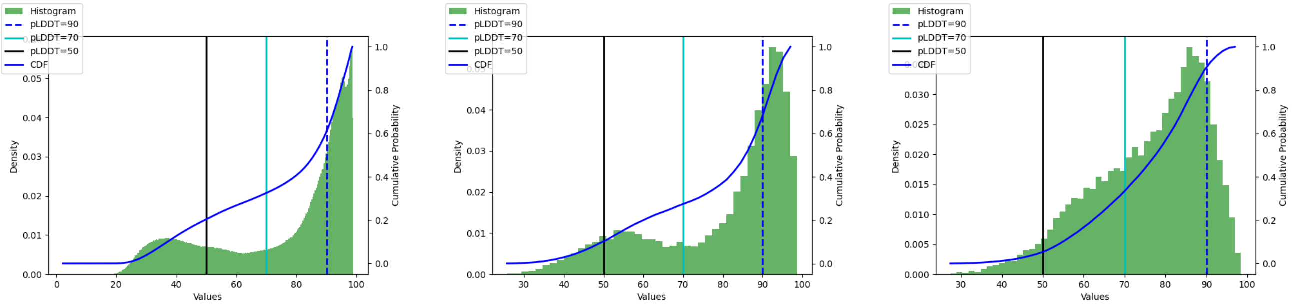

pLDDT all pLDDT median pLDDT mean

Figure S42: **CElegans: pLDDT statistics**

| | 50 | 70 | 90 |
|---|---|---|---|
| CElegans/all | 0.202 | 0.323 | 0.597 |
| CElegans/median | 0.086 | 0.262 | 0.598 |
| CElegans/mean | 0.044 | 0.306 | 0.859 |

Table S3: **CElegans: CDF pLDDT**

- Worst cases, top 3: ['File CElegans/AF-Q09239-F1-model_v4.pdb has 174 a.a.; pLDDT min/med/-max: 17.58 25.67 44.68', 'File CElegans/AF-A0A4V0IL95-F1-model_v4.pdb has 334 a.a.; pLDDT min/med/max: 19.72 25.79 59.21', 'File CElegans/AF-Q95XI1-F1-model_v4.pdb has 337 a.a.; pLDDT min/med/max: 20.36 26.09 47.80']
- Median cases, 3 of them: ['File CElegans/AF-O45170-F1-model_v4.pdb has 367 a.a.; pLDDT min/med/-max: 35.03 87.50 97.68', 'File CElegans/AF-Q20779-F1-model_v4.pdb has 128 a.a.; pLDDT min/med/-max: 46.68 87.50 97.83', 'File CElegans/AF-Q22109-F1-model_v4.pdb has 591 a.a.; pLDDT min/med/-max: 20.74 87.50 98.45']
- Top cases, top 3: ['File CElegans/AF-P41977-F1-model_v4.pdb has 218 a.a.; pLDDT min/med/max: 42.35 98.65 98.94', 'File CElegans/AF-P31161-F1-model_v4.pdb has 221 a.a.; pLDDT min/med/max: 44.21 98.66 98.94', 'File CElegans/AF-P34629-F1-model_v4.pdb has 491 a.a.; pLDDT min/med/max: 76.38 98.78 98.97']

#### S6.6 DDiscoideum

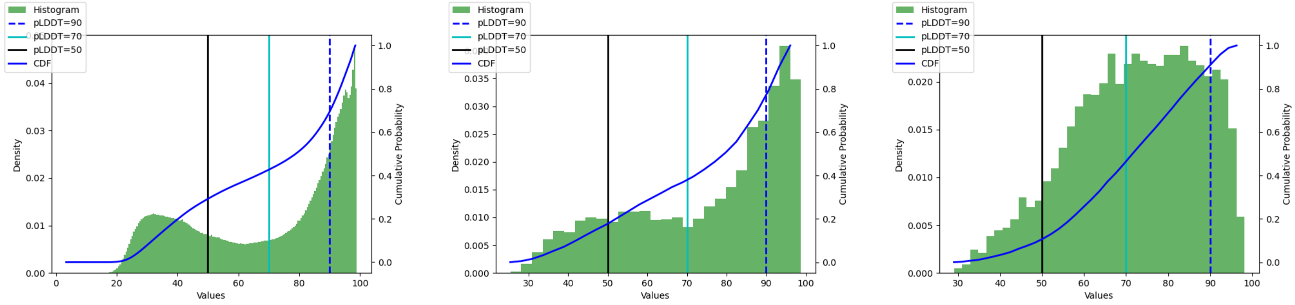

pLDDT all    pLDDT median    pLDDT mean

Figure S43: **DDiscoideum: pLDDT statistics**

| | 50 | 70 | 90 |
|---|---|---|---|
| DDiscoideum/all | 0.288 | 0.423 | 0.681 |
| DDiscoideum/median | 0.153 | 0.356 | 0.685 |
| DDiscoideum/mean | 0.089 | 0.423 | 0.868 |

Table S4: **DDiscoideum: CDF pLDDT**

- Worst cases, top 3: ['File DDiscoideum/AF-Q559X9-F1-model_v4.pdb has 271 a.a.; pLDDT min/med/-max: 20.17 25.36 83.15', 'File DDiscoideum/AF-Q86AT9-F1-model_v4.pdb has 1736 a.a.; pLDDT min/med/max: 17.61 25.88 97.02', 'File DDiscoideum/AF-Q55H25-F1-model_v4.pdb has 235 a.a.; pLDDT min/med/max: 18.94 27.07 46.51']
- Median cases, 3 of them: ['File DDiscoideum/AF-Q54K32-F1-model_v4.pdb has 822 a.a.; pLDDT min/med/max: 24.39 82.11 98.46', 'File DDiscoideum/AF-Q54HJ2-F1-model_v4.pdb has 678 a.a.; pLDDT min/med/max: 25.46 82.12 95.84', 'File DDiscoideum/AF-Q54GD1-F1-model_v4.pdb has 451 a.a.; pLDDT min/med/max: 27.59 82.12 98.76']
- Top cases, top 3: ['File DDiscoideum/AF-Q55CR9-F1-model_v4.pdb has 98 a.a.; pLDDT min/med/-max: 54.58 98.62 98.91', 'File DDiscoideum/AF-P42530-F1-model_v4.pdb has 257 a.a.; pLDDT min/med/max: 57.40 98.63 98.95', 'File DDiscoideum/AF-Q55DL0-F1-model_v4.pdb has 503 a.a.; pLDDT min/med/max: 36.56 98.82 98.97']

#### S6.7 DMelanogaster

| | 50 | 70 | 90 |
|---|---|---|---|
| DMelanogaster/all | 0.281 | 0.387 | 0.623 |
| DMelanogaster/median | 0.125 | 0.291 | 0.579 |
| DMelanogaster/mean | 0.047 | 0.340 | 0.836 |

Table S5: **DMelanogaster: CDF pLDDT**

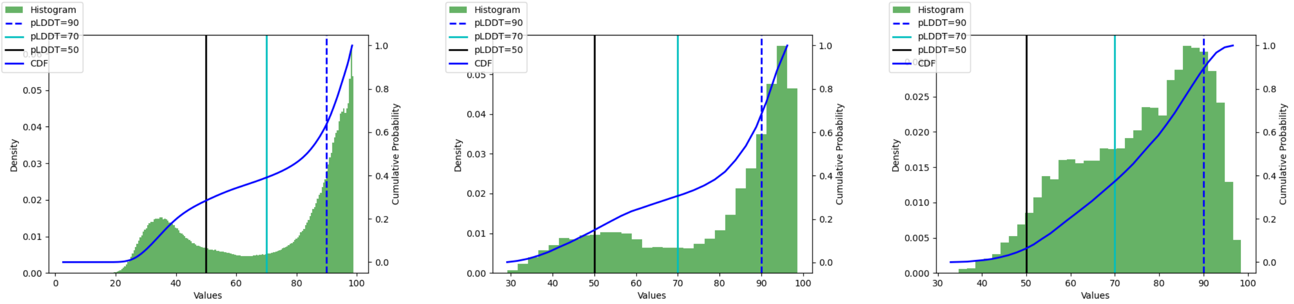

pLDDT all | pLDDT median | pLDDT mean

Figure S44: **DMelanogaster: pLDDT statistics**

- Worst cases, top 3: ['File DMelanogaster/AF-B7Z141-F1-model_v4.pdb has 2684 a.a.; pLDDT min/med/-max: 19.04 29.09 92.06', 'File DMelanogaster/AF-B7Z103-F1-model_v4.pdb has 259 a.a.; pLDDT min/med/max: 20.97 29.17 96.23', 'File DMelanogaster/AF-Q8ST83-F1-model_v4.pdb has 520 a.a.; pLDDT min/med/max: 17.56 29.19 95.79']
- Median cases, 3 of them: ['File DMelanogaster/AF-Q9VYG7-F1-model_v4.pdb has 559 a.a.; pLDDT min/med/max: 27.49 87.50 98.30', 'File DMelanogaster/AF-Q9VC94-F1-model_v4.pdb has 1430 a.a.; pLDDT min/med/max: 20.49 87.50 97.44', 'File DMelanogaster/AF-Q9VP13-F1-model_v4.pdb has 232 a.a.; pLDDT min/med/max: 42.58 87.51 97.94']
- Top cases, top 3: ['File DMelanogaster/AF-Q9VA37-F1-model_v4.pdb has 187 a.a.; pLDDT min/med/-max: 66.48 98.61 98.96', 'File DMelanogaster/AF-Q9VLC5-F1-model_v4.pdb has 520 a.a.; pLDDT min/med/max: 27.37 98.72 98.98', 'File DMelanogaster/AF-Q9VMC9-F1-model_v4.pdb has 300 a.a.; pLDDT min/med/max: 68.22 98.75 98.96']

#### S6.8 DRerio

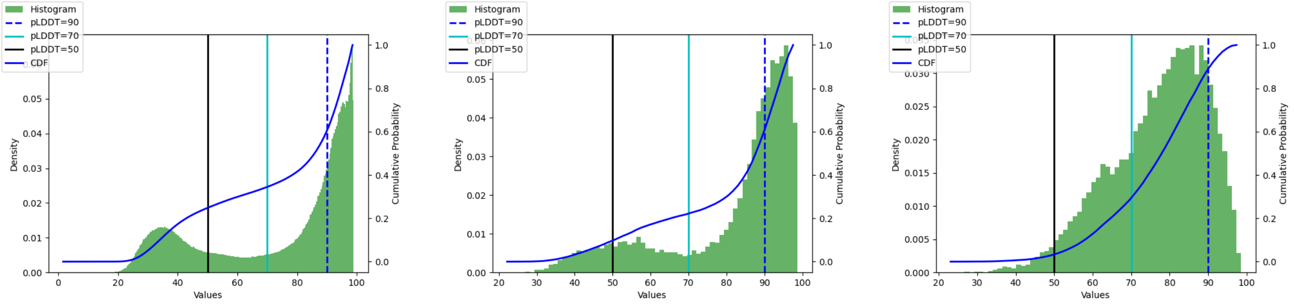

pLDDT all | pLDDT median | pLDDT mean

Figure S45: **DRerio: pLDDT statistics**

- Worst cases, top 3: ['File DRerio/AF-A0A0G2L439-F1-model_v4.pdb has 449 a.a.; pLDDT min/med/-max: 17.39 22.17 41.39', 'File DRerio/AF-A0A0G2KVY8-F1-model_v4.pdb has 349 a.a.; pLDDT min/med/max: 19.86 24.96 57.23', 'File DRerio/AF-A0A0G2KMU6-F1-model_v4.pdb has 374 a.a.; pLDDT min/med/max: 18.37 25.12 49.91']

|  | 50 | 70 | 90 |
|---|---|---|---|
| DRerio/all | 0.246 | 0.343 | 0.593 |
| DRerio/median | 0.089 | 0.216 | 0.549 |
| DRerio/mean | 0.027 | 0.270 | 0.855 |

Table S6: **DRerio: CDF pLDDT**

- Median cases, 3 of them: ['File DRerio/AF-A0A0G2KCZ7-F1-model_v4.pdb has 467 a.a.; pLDDT min/med/max: 29.62 88.93 97.06', 'File DRerio/AF-E7F647-F1-model_v4.pdb has 1015 a.a.; pLDDT min/med/max: 26.64 88.93 96.79', 'File DRerio/AF-E7FBI7-F1-model_v4.pdb has 203 a.a.; pLDDT min/med/max: 33.39 88.93 97.54']
- Top cases, top 3: ['File DRerio/AF-A0A0G2KZR6-F1-model_v4.pdb has 327 a.a.; pLDDT min/med/max: 87.09 98.74 98.96', 'File DRerio/AF-Q90Y03-F1-model_v4.pdb has 518 a.a.; pLDDT min/med/max: 25.60 98.75 98.98', 'File DRerio/AF-A2BGR9-F1-model_v4.pdb has 516 a.a.; pLDDT min/med/max: 25.45 98.78 98.98']

#### S6.9 EColi

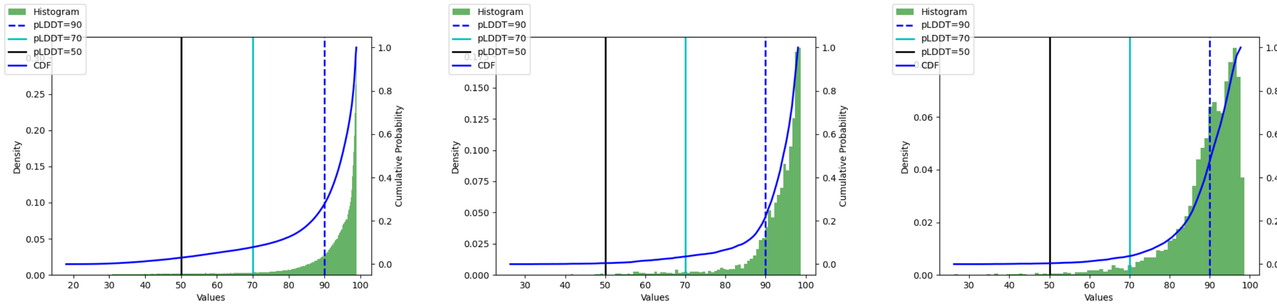

pLDDT all | pLDDT median | pLDDT mean

Figure S46: **EColi: pLDDT statistics**

|  | 50 | 70 | 90 |
|---|---|---|---|
| EColi/all | 0.029 | 0.078 | 0.275 |
| EColi/median | 0.005 | 0.034 | 0.190 |
| EColi/mean | 0.004 | 0.034 | 0.411 |

Table S7: **EColi: CDF pLDDT**

- Worst cases, top 3: ['File EColi/AF-P28697-F1-model_v4.pdb has 161 a.a.; pLDDT min/med/max: 20.16 26.32 31.97', 'File EColi/AF-P76122-F1-model_v4.pdb has 111 a.a.; pLDDT min/med/max: 22.69 32.35 52.00', 'File EColi/AF-P62066-F1-model_v4.pdb has 106 a.a.; pLDDT min/med/max: 26.10 34.23 64.50']
- Median cases, 3 of them: ['File EColi/AF-P37195-F1-model_v4.pdb has 176 a.a.; pLDDT min/med/max: 79.69 95.17 98.41', 'File EColi/AF-P00894-F1-model_v4.pdb has 163 a.a.; pLDDT min/med/-

max: 68.33 95.17 98.69', 'File EColi/AF-P27125-F1-model_v4.pdb has 344 a.a.; pLDDT min/med/-max: 68.75 95.17 98.60']

- Top cases, top 3: ['File EColi/AF-P0AF18-F1-model_v4.pdb has 382 a.a.; pLDDT min/med/max: 89.08 98.84 98.97', 'File EColi/AF-P27302-F1-model_v4.pdb has 663 a.a.; pLDDT min/med/max: 80.19 98.84 98.98', 'File EColi/AF-P77215-F1-model_v4.pdb has 401 a.a.; pLDDT min/med/max: 91.54 98.86 98.97']

#### S6.10 GMax

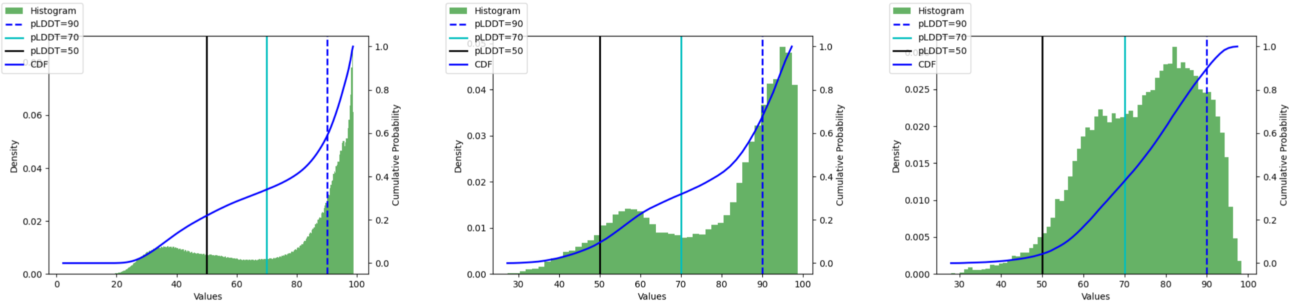

pLDDT all | pLDDT median | pLDDT mean

Figure S47: **GMax: pLDDT statistics**

| | 50 | 70 | 90 |
|---|---|---|---|
| GMax/all | 0.217 | 0.338 | 0.577 |
| GMax/median | 0.079 | 0.306 | 0.618 |
| GMax/mean | 0.036 | 0.356 | 0.873 |

Table S8: **GMax: CDF pLDDT**

- Worst cases, top 3: ['File GMax/AF-A0A0R0H4J2-F1-model_v4.pdb has 423 a.a.; pLDDT min/med/-max: 19.88 27.26 90.64', 'File GMax/AF-K7K902-F1-model_v4.pdb has 201 a.a.; pLDDT min/med/-max: 18.91 27.26 45.67', 'File GMax/AF-K7MPI4-F1-model_v4.pdb has 139 a.a.; pLDDT min/med/-max: 21.07 27.35 38.29']
- Median cases, 3 of them: ['File GMax/AF-I1KSS4-F1-model_v4.pdb has 196 a.a.; pLDDT min/med/-max: 31.72 86.09 97.85', 'File GMax/AF-I1LIH4-F1-model_v4.pdb has 620 a.a.; pLDDT min/med/-max: 28.18 86.09 97.42', 'File GMax/AF-I1LVU3-F1-model_v4.pdb has 604 a.a.; pLDDT min/med/-max: 22.17 86.09 98.78']
- Top cases, top 3: ['File GMax/AF-K7MRF5-F1-model_v4.pdb has 401 a.a.; pLDDT min/med/max: 51.92 98.80 98.97', 'File GMax/AF-C6TK73-F1-model_v4.pdb has 401 a.a.; pLDDT min/med/max: 54.78 98.81 98.98', 'File GMax/AF-I1M4B9-F1-model_v4.pdb has 299 a.a.; pLDDT min/med/max: 46.34 98.82 98.97']

#### S6.11 HPylori

- Worst cases, top 3: ['File HPylori/AF-O25264-F1-model_v4.pdb has 535 a.a.; pLDDT min/med/max: 20.88 33.20 53.61', 'File HPylori/AF-O25381-F1-model_v4.pdb has 449 a.a.; pLDDT min/med/max:

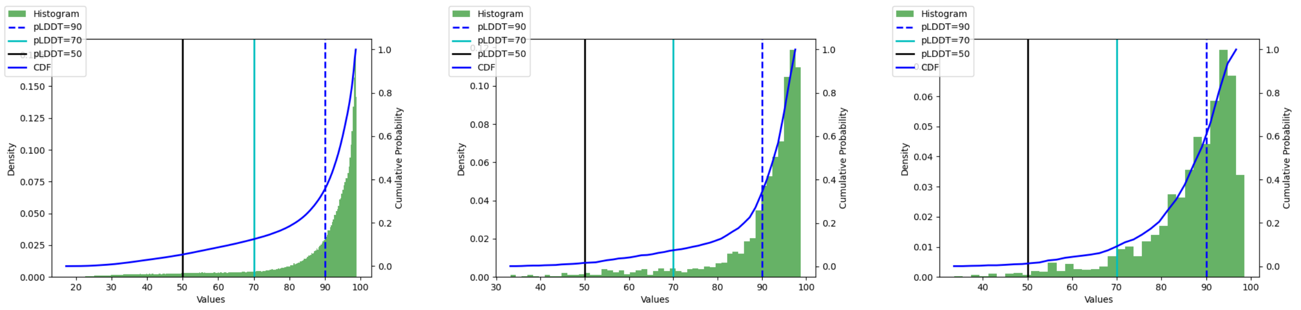

pLDDT all | pLDDT median | pLDDT mean

Figure S48: **HPylori: pLDDT statistics**

| | 50 | 70 | 90 |
|---|---|---|---|
| HPylori/all | 0.053 | 0.123 | 0.347 |
| HPylori/median | 0.014 | 0.070 | 0.280 |
| HPylori/mean | 0.010 | 0.074 | 0.507 |

Table S9: **HPylori: CDF pLDDT**

20.40 34.06 86.43', 'File HPylori/AF-O25271-F1-model_v4.pdb has 306 a.a.; pLDDT min/med/max: 21.60 36.29 88.71']

- Median cases, 3 of them: ['File HPylori/AF-O25499-F1-model_v4.pdb has 114 a.a.; pLDDT min/med/max: 40.28 93.95 98.78', 'File HPylori/AF-O25546-F1-model_v4.pdb has 202 a.a.; pLDDT min/med/max: 35.54 93.96 98.28', 'File HPylori/AF-P60325-F1-model_v4.pdb has 74 a.a.; pLDDT min/med/max: 36.35 93.96 96.70']
- Top cases, top 3: ['File HPylori/AF-P77872-F1-model_v4.pdb has 505 a.a.; pLDDT min/med/max: 43.77 98.79 98.95', 'File HPylori/AF-O24890-F1-model_v4.pdb has 330 a.a.; pLDDT min/med/max: 85.98 98.81 98.96', 'File HPylori/AF-O25452-F1-model_v4.pdb has 292 a.a.; pLDDT min/med/max: 90.66 98.83 98.97']

#### S6.12 HSapiens

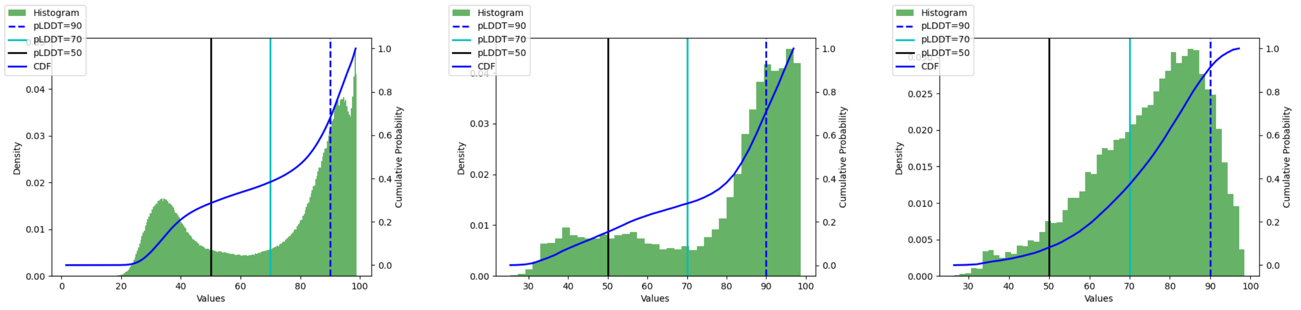

pLDDT all | pLDDT median | pLDDT mean

Figure S49: **HSapiens: pLDDT statistics**

| | 50 | 70 | 90 |
|---|---|---|---|
| HSapiens/all | 0.284 | 0.382 | 0.666 |
| HSapiens/median | 0.139 | 0.274 | 0.630 |
| HSapiens/mean | 0.071 | 0.344 | 0.877 |

Table S10: **HSapiens: CDF pLDDT**

- Worst cases, top 3: ['File HSapiens/AF-Q96MU5-F1-model_v4.pdb has 243 a.a.; pLDDT min/med/max: 19.88 25.35 38.27', 'File HSapiens/AF-Q6ZR03-F1-model_v4.pdb has 302 a.a.; pLDDT min/med/max: 21.04 26.27 46.20', 'File HSapiens/AF-Q96M85-F1-model_v4.pdb has 177 a.a.; pLDDT min/med/max: 20.02 26.32 42.34']
- Median cases, 3 of them: ['File HSapiens/AF-P56159-F1-model_v4.pdb has 465 a.a.; pLDDT min/med/max: 25.40 86.54 98.58', 'File HSapiens/AF-Q96N96-F1-model_v4.pdb has 652 a.a.; pLDDT min/med/max: 24.37 86.54 98.41', 'File HSapiens/AF-Q9H009-F1-model_v4.pdb has 215 a.a.; pLDDT min/med/max: 36.28 86.54 95.84']
- Top cases, top 3: ['File HSapiens/AF-Q99497-F1-model_v4.pdb has 189 a.a.; pLDDT min/med/max: 67.04 98.81 98.97', 'File HSapiens/AF-P00352-F1-model_v4.pdb has 501 a.a.; pLDDT min/med/max: 28.66 98.82 98.98', 'File HSapiens/AF-P21549-F1-model_v4.pdb has 392 a.a.; pLDDT min/med/max: 58.71 98.83 98.98']

#### S6.13 MJannaschii

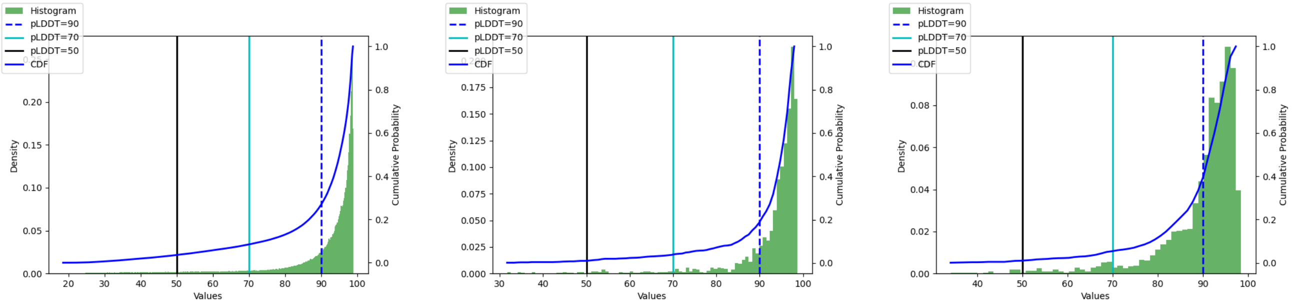

pLDDT all | pLDDT median | pLDDT mean

Figure S50: **MJannaschii: pLDDT statistics**

| | 50 | 70 | 90 |
|---|---|---|---|
| MJannaschii/all | 0.035 | 0.084 | 0.264 |
| MJannaschii/median | 0.010 | 0.036 | 0.168 |
| MJannaschii/mean | 0.009 | 0.050 | 0.326 |

Table S11: **MJannaschii: CDF pLDDT**

- Worst cases, top 3: ['File MJannaschii/AF-Q58686-F1-model_v4.pdb has 312 a.a.; pLDDT min/med/max: 19.67 31.70 84.81', 'File MJannaschii/AF-Q58912-F1-model_v4.pdb has 278 a.a.; pLDDT min/med/max: 19.48 32.08 65.08', 'File MJannaschii/AF-Q60308-F1-model_v4.pdb has 237 a.a.; pLDDT min/med/max: 22.90 34.61 71.98']

- Median cases, 3 of them: ['File MJannaschii/AF-Q57707-F1-model_v4.pdb has 202 a.a.; pLDDT min/med/max: 45.71 95.86 98.84', 'File MJannaschii/AF-Q57706-F1-model_v4.pdb has 404 a.a.; pLDDT min/med/max: 56.43 95.87 98.87', 'File MJannaschii/AF-Q58280-F1-model_v4.pdb has 620 a.a.; pLDDT min/med/max: 35.71 95.87 98.72']

- Top cases, top 3: ['File MJannaschii/AF-Q57764-F1-model_v4.pdb has 165 a.a.; pLDDT min/med/-max: 49.70 98.74 98.95', 'File MJannaschii/AF-Q60358-F1-model_v4.pdb has 396 a.a.; pLDDT min/med/-max: 64.69 98.76 98.98', 'File MJannaschii/AF-Q58691-F1-model_v4.pdb has 218 a.a.; pLDDT min/med/-max: 88.41 98.77 98.97']

#### S6.14 MMusculus

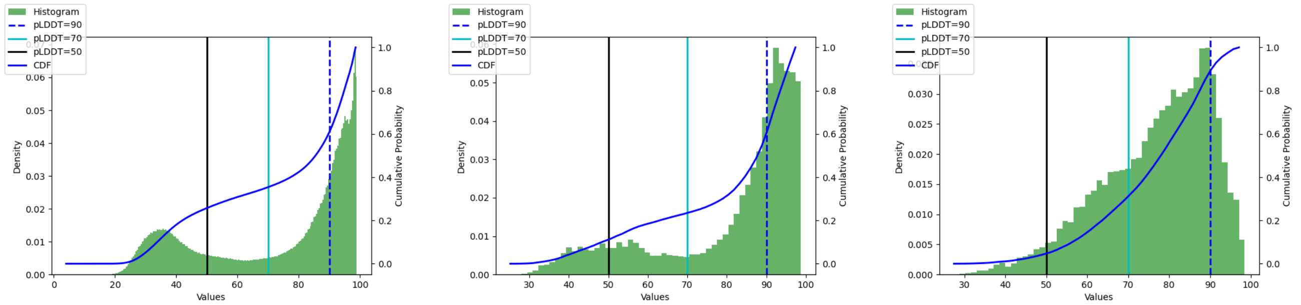

pLDDT all　　pLDDT median　　pLDDT mean

Figure S51: **MMusculus: pLDDT statistics**

| | 50 | 70 | 90 |
|---|---|---|---|
| MMusculus/all | 0.255 | 0.352 | 0.597 |
| MMusculus/median | 0.102 | 0.228 | 0.534 |
| MMusculus/mean | 0.040 | 0.285 | 0.842 |

Table S12: **MMusculus: CDF pLDDT**

- Worst cases, top 3: ['File MMusculus/AF-Q3UUJ8-F1-model_v4.pdb has 172 a.a.; pLDDT min/med/-max: 20.56 25.16 50.36', 'File MMusculus/AF-Q8BQQ8-F1-model_v4.pdb has 176 a.a.; pLDDT min/med/max: 20.19 26.95 49.81', 'File MMusculus/AF-Q569Q2-F1-model_v4.pdb has 183 a.a.; pLDDT min/med/max: 17.71 28.42 40.28']

- Median cases, 3 of them: ['File MMusculus/AF-Q80VL1-F1-model_v4.pdb has 560 a.a.; pLDDT min/med/max: 25.35 89.19 98.09', 'File MMusculus/AF-Q8K5B1-F1-model_v4.pdb has 716 a.a.; pLDDT min/med/max: 24.58 89.19 97.79', 'File MMusculus/AF-Q9CQR5-F1-model_v4.pdb has 170 a.a.; pLDDT min/med/max: 44.38 89.19 96.99']

- Top cases, top 3: ['File MMusculus/AF-Q8VC28-F1-model_v4.pdb has 323 a.a.; pLDDT min/med/-max: 27.86 98.78 98.97', 'File MMusculus/AF-Q99LX0-F1-model_v4.pdb has 189 a.a.; pLDDT min/med/-max: 65.04 98.78 98.97', 'File MMusculus/AF-Q62148-F1-model_v4.pdb has 518 a.a.; pLDDT min/med/-max: 28.41 98.80 98.98']

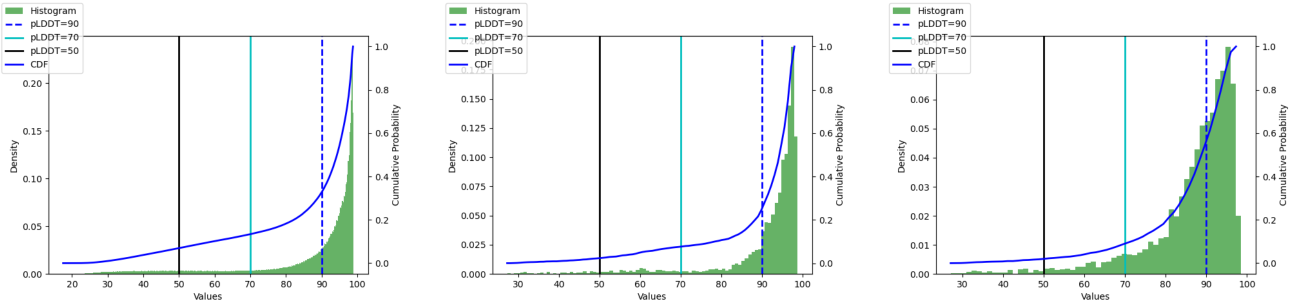

pLDDT all    pLDDT median    pLDDT mean

Figure S52: **MTuberculosis: pLDDT statistics**

| | 50 | 70 | 90 |
|---|---|---|---|
| MTuberculosis/all | 0.069 | 0.133 | 0.322 |
| MTuberculosis/median | 0.022 | 0.075 | 0.220 |
| MTuberculosis/mean | 0.018 | 0.082 | 0.497 |

Table S13: **MTuberculosis: CDF pLDDT**

#### S6.15 MTuberculosis

- Worst cases, top 3: ['File MTuberculosis/AF-P9WLQ3-F1-model_v4.pdb has 173 a.a.; pLDDT min/med/-max: 18.69 27.23 49.23', 'File MTuberculosis/AF-P95044-F1-model_v4.pdb has 203 a.a.; pLDDT min/med/max: 17.77 27.38 38.24', 'File MTuberculosis/AF-P9WLX7-F1-model_v4.pdb has 253 a.a.; pLDDT min/med/max: 20.85 29.51 58.62']
- Median cases, 3 of them: ['File MTuberculosis/AF-P72007-F1-model_v4.pdb has 532 a.a.; pLDDT min/med/max: 38.35 95.19 98.88', 'File MTuberculosis/AF-P95233-F1-model_v4.pdb has 372 a.a.; pLDDT min/med/max: 28.08 95.19 98.74', 'File MTuberculosis/AF-P9WJK9-F1-model_v4.pdb has 410 a.a.; pLDDT min/med/max: 26.08 95.19 98.89']
- Top cases, top 3: ['File MTuberculosis/AF-P9WQD9-F1-model_v4.pdb has 416 a.a.; pLDDT min/med/-max: 66.52 98.80 98.97', 'File MTuberculosis/AF-P9WGI9-F1-model_v4.pdb has 438 a.a.; pLDDT min/med/max: 42.72 98.82 98.97', 'File MTuberculosis/AF-I6XHI4-F1-model_v4.pdb has 391 a.a.; pLDDT min/med/max: 82.88 98.84 98.98']

#### S6.16 OryzaSativa

| | 50 | 70 | 90 |
|---|---|---|---|
| OryzaSativa/all | 0.257 | 0.408 | 0.623 |
| OryzaSativa/median | 0.199 | 0.486 | 0.717 |
| OryzaSativa/mean | 0.155 | 0.520 | 0.911 |

Table S14: **OryzaSativa: CDF pLDDT**

- Worst cases, top 3: ['File OryzaSativa/AF-Q6K7L0-F1-model_v4.pdb has 173 a.a.; pLDDT min/med/-max: 19.39 23.67 85.80', 'File OryzaSativa/AF-A0A0P0VJ79-F1-model_v4.pdb has 259 a.a.; pLDDT

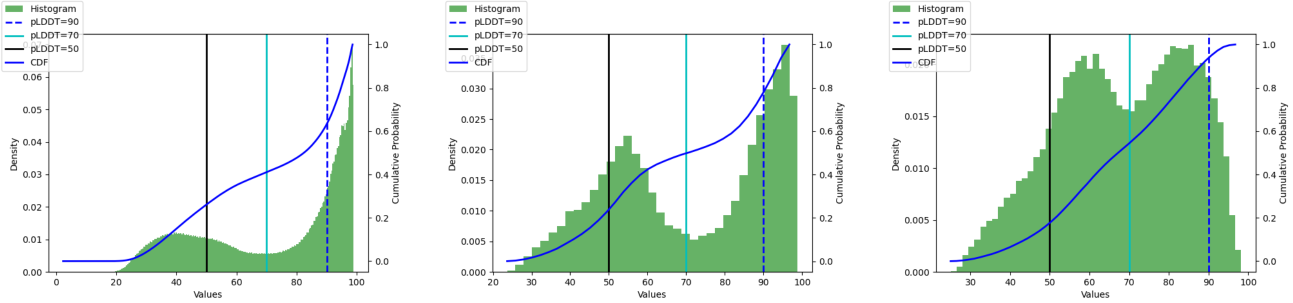

pLDDT all pLDDT median pLDDT mean

Figure S53: **OryzaSativa: pLDDT statistics**

min/med/max: 17.43 24.27 36.13', 'File OryzaSativa/AF-A0A0P0WVM6-F1-model_v4.pdb has 504 a.a.; pLDDT min/med/max: 18.68 24.84 78.76']

- Median cases, 3 of them: ['File OryzaSativa/AF-A0A0N7KPX1-F1-model_v4.pdb has 262 a.a.; pLDDT min/med/max: 35.91 72.61 97.84', 'File OryzaSativa/AF-A0A0P0WQU0-F1-model_v4.pdb has 77 a.a.; pLDDT min/med/max: 40.54 72.61 96.23', 'File OryzaSativa/AF-Q7Y0W5-F1-model_v4.pdb has 341 a.a.; pLDDT min/med/max: 26.78 72.61 97.53']

- Top cases, top 3: ['File OryzaSativa/AF-Q8L7J2-F1-model_v4.pdb has 521 a.a.; pLDDT min/med/-max: 28.69 98.78 98.96', 'File OryzaSativa/AF-Q6ZFI6-F1-model_v4.pdb has 402 a.a.; pLDDT min/med/-max: 45.10 98.80 98.98', 'File OryzaSativa/AF-Q75I93-F1-model_v4.pdb has 504 a.a.; pLDDT min/med/-max: 40.54 98.85 98.97']

#### S6.17 PAeruginosa

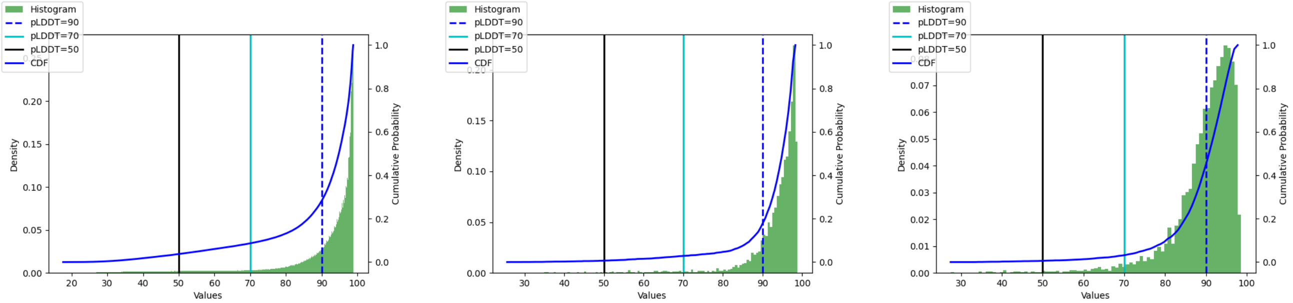

pLDDT all pLDDT median pLDDT mean

Figure S54: **PAeruginosa: pLDDT statistics**

- Worst cases, top 3: ['File PAeruginosa/AF-Q9I728-F1-model_v4.pdb has 198 a.a.; pLDDT min/med/-max: 20.78 25.56 47.94', 'File PAeruginosa/AF-Q9I2K8-F1-model_v4.pdb has 327 a.a.; pLDDT min/med/-max: 20.37 26.57 89.72', 'File PAeruginosa/AF-Q9I665-F1-model_v4.pdb has 224 a.a.; pLDDT min/med/-max: 21.50 29.81 73.31']

- Median cases, 3 of them: ['File PAeruginosa/AF-Q9HTB7-F1-model_v4.pdb has 479 a.a.; pLDDT min/med/max: 47.61 95.54 98.73', 'File PAeruginosa/AF-Q9HVR7-F1-model_v4.pdb has 303 a.a.;

| | 50 | 70 | 90 |
|---|---|---|---|
| PAeruginosa/all | 0.036 | 0.086 | 0.278 |
| PAeruginosa/median | 0.006 | 0.028 | 0.156 |
| PAeruginosa/mean | 0.005 | 0.030 | 0.396 |

Table S15: **PAeruginosa: CDF pLDDT**

pLDDT min/med/max: 39.46 95.54 98.50', 'File PAeruginosa/AF-Q9HXS6-F1-model_v4.pdb has 267 a.a.; pLDDT min/med/max: 37.32 95.54 98.81']

- Top cases, top 3: ['File PAeruginosa/AF-Q9HTP2-F1-model_v4.pdb has 497 a.a.; pLDDT min/med/-max: 54.80 98.85 98.98', 'File PAeruginosa/AF-Q9HWU0-F1-model_v4.pdb has 461 a.a.; pLDDT min/med/max: 42.05 98.85 98.98', 'File PAeruginosa/AF-Q9I121-F1-model_v4.pdb has 159 a.a.; pLDDT min/med/max: 90.82 98.86 98.96']

#### S6.18 PFalciparum

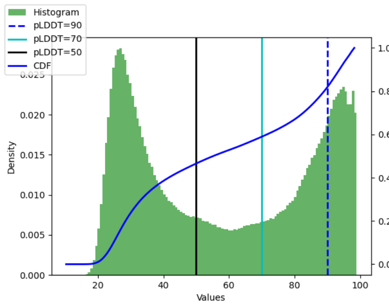

pLDDT all

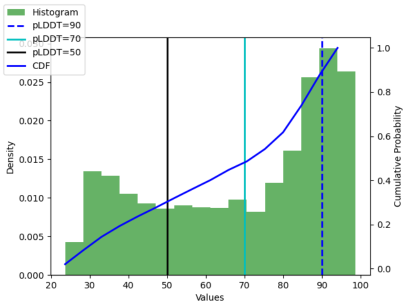

pLDDT median

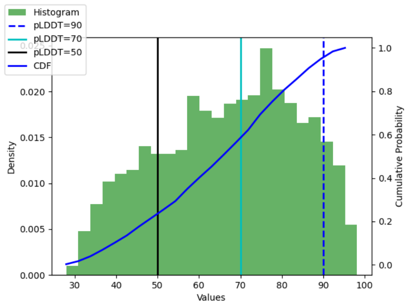

pLDDT mean

Figure S55: **PFalciparum: pLDDT statistics**

| | 50 | 70 | 90 |
|---|---|---|---|
| PFalciparum/all | 0.460 | 0.584 | 0.804 |
| PFalciparum/median | 0.261 | 0.441 | 0.757 |
| PFalciparum/mean | 0.197 | 0.529 | 0.915 |

Table S16: **PFalciparum: CDF pLDDT**

- Worst cases, top 3: ['File PFalciparum/AF-Q8ILT4-F1-model_v4.pdb has 2280 a.a.; pLDDT min/med/-max: 15.91 23.66 96.49', 'File PFalciparum/AF-C0H4Q1-F1-model_v4.pdb has 2580 a.a.; pLDDT min/med/max: 17.34 24.41 96.81', 'File PFalciparum/AF-C6KT74-F1-model_v4.pdb has 2404 a.a.; pLDDT min/med/max: 16.33 24.84 98.69']
- Median cases, 3 of them: ['File PFalciparum/AF-Q8IFL1-F1-model_v4.pdb has 231 a.a.; pLDDT min/med/max: 33.35 76.56 93.89', 'File PFalciparum/AF-Q8I419-F1-model_v4.pdb has 366 a.a.; pLDDT min/med/max: 27.79 76.59 97.05', 'File PFalciparum/AF-Q8IIM7-F1-model_v4.pdb has 698 a.a.; pLDDT min/med/max: 22.18 76.60 95.19']

- Top cases, top 3: ['File PFalciparum/AF-Q8IKK7-F1-model_v4.pdb has 337 a.a.; pLDDT min/med/-max: 69.22 98.66 98.94', 'File PFalciparum/AF-Q8IAY6-F1-model_v4.pdb has 198 a.a.; pLDDT min/med/max: 66.94 98.69 98.94', 'File PFalciparum/AF-Q8I3X4-F1-model_v4.pdb has 245 a.a.; pLDDT min/med/max: 49.70 98.76 98.97']

#### S6.19 RattusNorvegicus

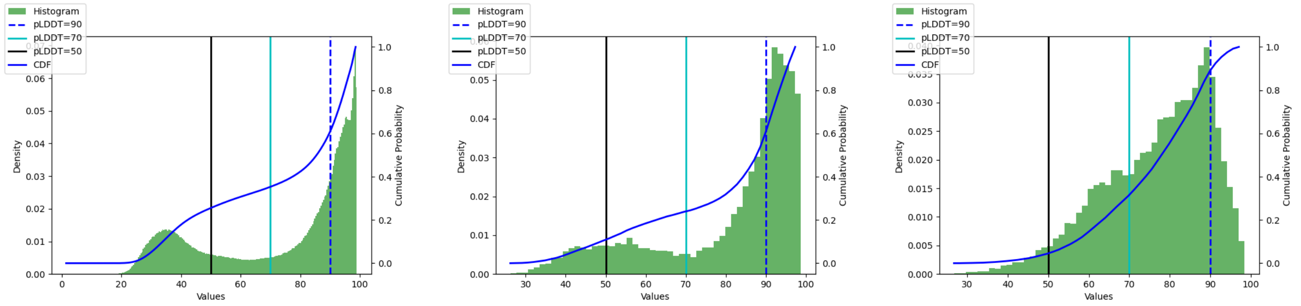

pLDDT all    pLDDT median    pLDDT mean

Figure S56: **RattusNorvegicus: pLDDT statistics**

| | 50 | 70 | 90 |
|---|---|---|---|
| RattusNorvegicus/all | 0.253 | 0.351 | 0.596 |
| RattusNorvegicus/median | 0.098 | 0.232 | 0.535 |
| RattusNorvegicus/mean | 0.038 | 0.290 | 0.843 |

Table S17: **RattusNorvegicus: CDF pLDDT**

- Worst cases, top 3: ['File RattusNorvegicus/AF-D4A5K8-F1-model_v4.pdb has 164 a.a.; pLDDT min/med/max: 20.30 26.16 36.12', 'File RattusNorvegicus/AF-F7F730-F1-model_v4.pdb has 218 a.a.; pLDDT min/med/max: 19.72 26.42 60.73', 'File RattusNorvegicus/AF-Q6QI15-F1-model_v4.pdb has 396 a.a.; pLDDT min/med/max: 20.68 26.84 48.50']

- Median cases, 3 of them: ['File RattusNorvegicus/AF-Q63258-F1-model_v4.pdb has 1135 a.a.; pLDDT min/med/max: 22.15 89.17 98.63', 'File RattusNorvegicus/AF-B2RZ71-F1-model_v4.pdb has 240 a.a.; pLDDT min/med/max: 32.07 89.17 98.66', 'File RattusNorvegicus/AF-D3ZQK4-F1-model_v4.pdb has 819 a.a.; pLDDT min/med/max: 19.17 89.18 98.32']

- Top cases, top 3: ['File RattusNorvegicus/AF-P51647-F1-model_v4.pdb has 501 a.a.; pLDDT min/med/-max: 35.91 98.78 98.98', 'File RattusNorvegicus/AF-P11884-F1-model_v4.pdb has 519 a.a.; pLDDT min/med/max: 30.63 98.79 98.96', 'File RattusNorvegicus/AF-Q03336-F1-model_v4.pdb has 299 a.a.; pLDDT min/med/max: 56.82 98.79 98.98']

#### S6.20 SAureus

- Worst cases, top 3: ['File SAureus/AF-Q2G012-F1-model_v4.pdb has 340 a.a.; pLDDT min/med/-max: 17.98 25.04 54.26', 'File SAureus/AF-Q2FUW1-F1-model_v4.pdb has 2271 a.a.; pLDDT min/med/-max: 21.21 29.66 98.86', 'File SAureus/AF-Q2G249-F1-model_v4.pdb has 159 a.a.; pLDDT min/med/-max: 22.92 34.85 52.05']

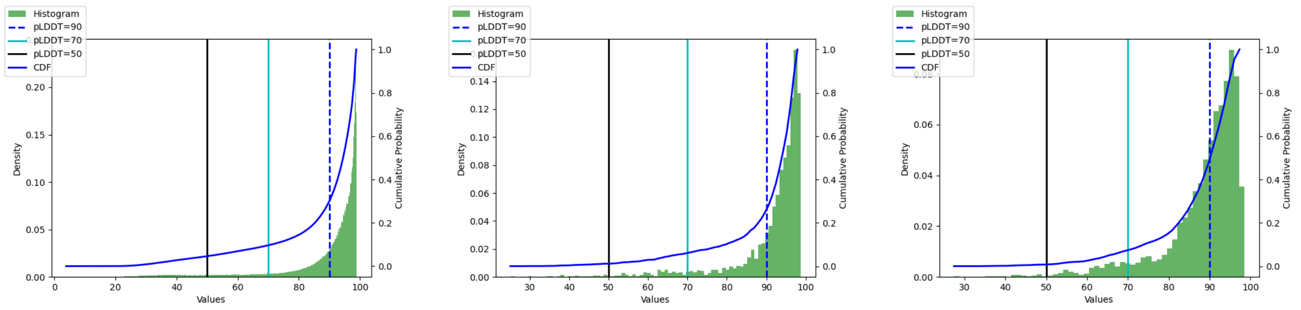

pLDDT all pLDDT median pLDDT mean

Figure S57: **SAureus: pLDDT statistics**

| | 50 | 70 | 90 |
|---|---|---|---|
| SAureus/all | 0.045 | 0.096 | 0.295 |
| SAureus/median | 0.012 | 0.057 | 0.229 |
| SAureus/mean | 0.007 | 0.068 | 0.421 |

Table S18: **SAureus: CDF pLDDT**

- Median cases, 3 of them: ['File SAureus/AF-Q2G2X8-F1-model_v4.pdb has 232 a.a.; pLDDT min/med/max: 26.38 94.82 98.55', 'File SAureus/AF-Q2FVK7-F1-model_v4.pdb has 288 a.a.; pLDDT min/med/max: 59.70 94.83 98.60', 'File SAureus/AF-Q2FVU5-F1-model_v4.pdb has 176 a.a.; pLDDT min/med/max: 38.29 94.83 98.41']
- Top cases, top 3: ['File SAureus/AF-P0A0K3-F1-model_v4.pdb has 171 a.a.; pLDDT min/med/max: 63.37 98.80 98.97', 'File SAureus/AF-Q2FV24-F1-model_v4.pdb has 560 a.a.; pLDDT min/med/max: 55.84 98.80 98.98', 'File SAureus/AF-Q2G160-F1-model_v4.pdb has 293 a.a.; pLDDT min/med/max: 46.53 98.83 98.95']

#### S6.21 SCerevisiae

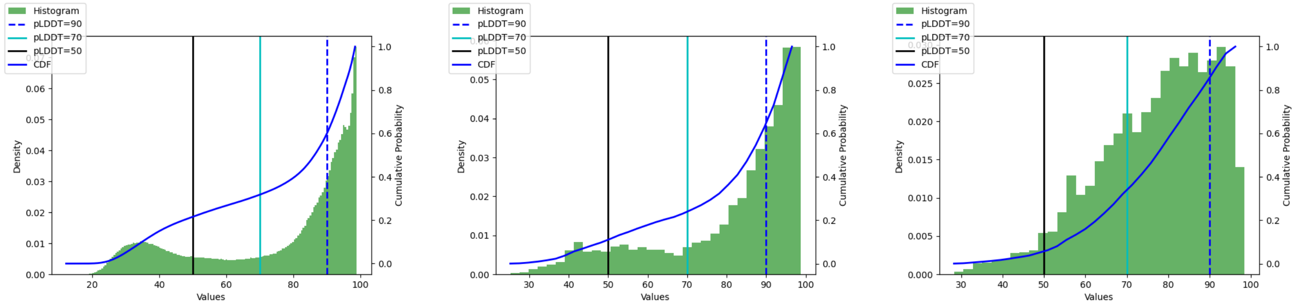

pLDDT all pLDDT median pLDDT mean

Figure S58: **SCerevisiae: pLDDT statistics**

- Worst cases, top 3: ['File SCerevisiae/AF-P38299-F1-model_v4.pdb has 523 a.a.; pLDDT min/med/max: 18.42 25.22 56.94', 'File SCerevisiae/AF-P38083-F1-model_v4.pdb has 404 a.a.; pLDDT min/med/-

| | 50 | 70 | 90 |
|---|---|---|---|
| SCerevisiae/all | 0.213 | 0.314 | 0.582 |
| SCerevisiae/median | 0.095 | 0.222 | 0.559 |
| SCerevisiae/mean | 0.045 | 0.294 | 0.791 |

Table S19: **SCerevisiae: CDF pLDDT**

max: 19.89 25.65 52.05', 'File SCerevisiae/AF-P53963-F1-model_v4.pdb has 612 a.a.; pLDDT min/med/-max: 18.67 26.82 81.53']

- Median cases, 3 of them: ['File SCerevisiae/AF-Q03516-F1-model_v4.pdb has 953 a.a.; pLDDT min/med/max: 22.63 88.32 98.50', 'File SCerevisiae/AF-Q06147-F1-model_v4.pdb has 687 a.a.; pLDDT min/med/max: 25.84 88.32 98.67', 'File SCerevisiae/AF-P36023-F1-model_v4.pdb has 863 a.a.; pLDDT min/med/max: 22.36 88.33 98.43']
- Top cases, top 3: ['File SCerevisiae/AF-P0CW40-F1-model_v4.pdb has 589 a.a.; pLDDT min/med/-max: 47.78 98.79 98.97', 'File SCerevisiae/AF-P53051-F1-model_v4.pdb has 589 a.a.; pLDDT min/med/-max: 50.17 98.81 98.96', 'File SCerevisiae/AF-Q03558-F1-model_v4.pdb has 400 a.a.; pLDDT min/med/-max: 68.18 98.82 98.98']

#### S6.22 SPombe

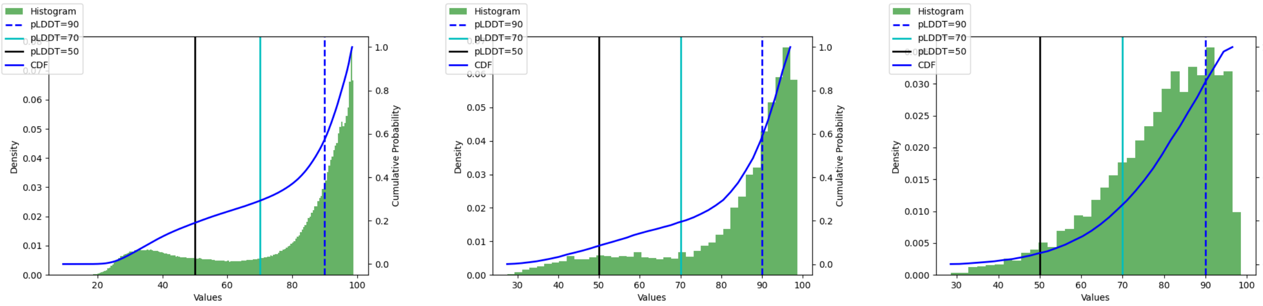

pLDDT all pLDDT median pLDDT mean

Figure S59: **SPombe: pLDDT statistics**

| | 50 | 70 | 90 |
|---|---|---|---|
| SPombe/all | 0.187 | 0.289 | 0.558 |
| SPombe/median | 0.074 | 0.183 | 0.503 |
| SPombe/mean | 0.041 | 0.236 | 0.764 |

Table S20: **SPombe: CDF pLDDT**

- Worst cases, top 3: ['File SPombe/AF-Q9P7E7-F1-model_v4.pdb has 166 a.a.; pLDDT min/med/max: 21.73 27.41 51.55', 'File SPombe/AF-Q09862-F1-model_v4.pdb has 339 a.a.; pLDDT min/med/max: 18.99 27.98 46.68', 'File SPombe/AF-Q9Y7R9-F1-model_v4.pdb has 283 a.a.; pLDDT min/med/max: 17.64 29.33 53.72']

- Median cases, 3 of them: ['File SPombe/AF-P32834-F1-model_v4.pdb has 533 a.a.; pLDDT min/med/-max: 20.51 89.94 98.59', 'File SPombe/AF-O13799-F1-model_v4.pdb has 1030 a.a.; pLDDT min/med/-max: 22.20 89.94 98.09', 'File SPombe/AF-Q09807-F1-model_v4.pdb has 701 a.a.; pLDDT min/med/-max: 35.87 89.95 98.79']
- Top cases, top 3: ['File SPombe/AF-P18253-F1-model_v4.pdb has 162 a.a.; pLDDT min/med/max: 77.46 98.65 98.93', 'File SPombe/AF-O42652-F1-model_v4.pdb has 409 a.a.; pLDDT min/med/max: 77.99 98.78 98.97', 'File SPombe/AF-Q10356-F1-model_v4.pdb has 191 a.a.; pLDDT min/med/max: 87.96 98.81 98.96']

#### S6.23 ZeaMays

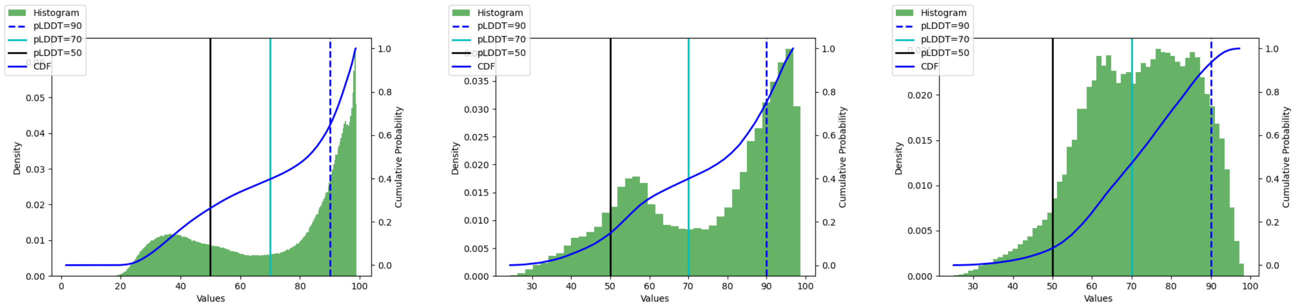

pLDDT all    pLDDT median    pLDDT mean

Figure S60: **ZeaMays: pLDDT statistics**

| | 50 | 70 | 90 |
|---|---|---|---|
| ZeaMays/all | 0.260 | 0.394 | 0.635 |
| ZeaMays/median | 0.123 | 0.382 | 0.687 |
| ZeaMays/mean | 0.068 | 0.443 | 0.911 |

Table S21: **ZeaMays: CDF pLDDT**

- Worst cases, top 3: ['File ZeaMays/AF-A0A1D6HY84-F1-model_v4.pdb has 348 a.a.; pLDDT min/med/-max: 19.54 24.33 41.44', 'File ZeaMays/AF-A0A1D6H3E1-F1-model_v4.pdb has 266 a.a.; pLDDT min/med/max: 19.77 24.50 37.39', 'File ZeaMays/AF-A0A1D6I485-F1-model_v4.pdb has 393 a.a.; pLDDT min/med/max: 18.10 24.60 69.80']
- Median cases, 3 of them: ['File ZeaMays/AF-B4FZ77-F1-model_v4.pdb has 174 a.a.; pLDDT min/med/-max: 39.15 81.90 91.30', 'File ZeaMays/AF-A0A1D6I1I2-F1-model_v4.pdb has 422 a.a.; pLDDT min/med/max: 22.95 81.91 98.22', 'File ZeaMays/AF-A0A1D6QK68-F1-model_v4.pdb has 50 a.a.; pLDDT min/med/max: 51.82 81.91 88.41']
- Top cases, top 3: ['File ZeaMays/AF-P00874-F1-model_v4.pdb has 476 a.a.; pLDDT min/med/max: 45.05 98.75 98.96', 'File ZeaMays/AF-B6TKZ3-F1-model_v4.pdb has 300 a.a.; pLDDT min/med/max: 41.45 98.77 98.97', 'File ZeaMays/AF-B6T171-F1-model_v4.pdb has 403 a.a.; pLDDT min/med/max: 48.91 98.79 98.98']

**S2 Q1. Contacts, packing, and whole genome predictions** 22

**S3 Q2. Predicted domains and their quality** 33

**S4 Q4. pLDDT values and fragmentation of AlphaFold reconstructions** 35

**S5 Whole genomes: arity** 38
S5.1 AThaliana . . . 38
S5.2 CAlbicans . . . 38
S5.3 CElegans . . . 39
S5.4 DDiscoideum . . . 39
S5.5 DMelanogaster . . . 39
S5.6 DRerio . . . 40
S5.7 EColi . . . 40
S5.8 GMax . . . 41
S5.9 HPylori . . . 41
S5.10HSapiens . . . 42
S5.11MJannaschii . . . 42
S5.12MMusculus . . . 43
S5.13MTuberculosis . . . 43
S5.14OryzaSativa . . . 44
S5.15PAeruginosa . . . 44
S5.16PFalciparum . . . 45
S5.17RattusNorvegicus . . . 45
S5.18SAureus . . . 46
S5.19SCerevisiae . . . 46
S5.20SPombe . . . 47
S5.21ZeaMays . . . 47

**S6 Whole genomes: pLDDT** 49
S6.1 pLDDT statistics per organism . . . 49
S6.2 pLDDT statistics per organism . . . 49
S6.3 AThaliana . . . 49
S6.4 CAlbicans . . . 50
S6.5 CElegans . . . 51
S6.6 DDiscoideum . . . 52
S6.7 DMelanogaster . . . 52
S6.8 DRerio . . . 53
S6.9 EColi . . . 54
S6.10GMax . . . 55
S6.11HPylori . . . 55
S6.12HSapiens . . . 56
S6.13MJannaschii . . . 57
S6.14MMusculus . . . 58
S6.15MTuberculosis . . . 59
S6.16OryzaSativa . . . 59
S6.17PAeruginosa . . . 60
S6.18PFalciparum . . . 61
S6.19RattusNorvegicus . . . 62
S6.20SAureus . . . 62
S6.21SCerevisiae . . . 63
S6.22SPombe . . . 64
S6.23ZeaMays . . . 65
